# Rapid proteotyping reveals cancer biology and drug response determinants in the NCI-60 cells

**DOI:** 10.1101/268953

**Authors:** Tiannan Guo, Augustin Luna, Vinodh N Rajapakse, Ching Chiek Koh, Zhicheng Wu, Michael P Menden, Yongran Cheng, Laurence Calzone, Loredana Martignetti, Alessandro Ori, Murat Iskar, Ludovic Gillet, Qing Zhong, Sudhir Varma, Uwe Schmitt, Peng Qiu, Yaoting Sun, Yi Zhu, Peter J Wild, Mathew J Garnett, Peer Bork, Martin Beck, Julio Saez-Rodriguez, William C. Reinhold, Chris Sander, Yves Pommier, Ruedi Aebersold

## Abstract

We describe the rapid and reproducible acquisition of quantitative proteome maps for the NCI-60 cancer cell lines and their use to reveal cancer biology and drug response determinants. Proteome datasets for the 60 cell lines were acquired in duplicate within 30 working days using pressure cycling technology and SWATH mass spectrometry. We consistently quantified 3,171 SwissProt proteotypic proteins across all cell lines, generating a data matrix with 0.1% missing values, allowing analyses of protein complexes and pathway activities across all the cancer cells. Systematic and integrative analysis of the genetic variation, mRNA expression and proteomic data of the NCI-60 cancer cell lines uncovered complementarity between different types of molecular data in the prediction of the response to 240 drugs. We additionally identified novel proteomic drug response determinants for clinically relevant chemotherapeutic and targeted therapies. We anticipate that this study represents a landmark effort toward the translational application of proteotypes, which reveal biological insights that are easily missed in the absence of proteomic data.

## Introduction

To date, mainly owing to the maturity and availability of high throughput DNA-and RNA-based techniques, forays into the molecular landscape of diseases, in particular cancers, have primarily focused on genomics and transcriptomics ^1–3^. Protein-level measurements, although showing great potential for providing the granularity and details necessary for personalized therapeutic decisions, are underutilized due to technical hurdles. Advances in data-dependent acquisition (DDA) mass spectrometry (MS) have permitted quantitative proteomic profiling of about 100 tumor samples using multi-dimensional fractionated MS analyses of each sample ^4–6^, demonstrating the added value of protein measurement in classifying tumor samples. Nevertheless, such DDA workflows suffer from relatively lower sample-throughput, relatively higher sample consumption and technical complexity, precluding their routine use in clinically relevant applications (*e.g.* drug response prediction) on the speed and scale achieved by genomic and transcriptomic approaches ^2, 3^.

To achieve reproducible and high throughput proteomic profiling, we have developed a workflow ^7, 8^ integrating pressure cycling technology (PCT), an emerging sample preparation method that accelerates and standardizes sample preparation for proteomic profiling ^9^, together with SWATH-MS, an MS-based proteomic technique that consists of data independent acquisition (DIA) and a targeted data analysis strategy with unique advantages over other MS-based proteomic methods ^10, 11^. With this technique all MS-measurable peptides of a sample are fragmented and recorded in a recursive fashion, thus generating digital proteome maps that can be used to reproducibly detect and quantify proteins across high numbers of samples without the need of isotope labeling. The PCT-SWATH technique thus significantly increases the sample throughput and data reproducibility providing excellent quantitative accuracy, and in the meantime reduces sample consumption to ca. 1 microgram of total peptide mass per sample ^7, 8^.

In this study, we describe the acquisition of proteome maps of the NCI-60 cell lines in duplicate by PCT-SWATH. The 120 proteome maps were acquired within 30 working days on a single instrument and each sample consumed ca. 1 microgram of total peptide mass. We consistently quantified 3,171 SwissProt proteotypic proteins across all cell lines, generating a data matrix (120 proteomes vs. 3171 proteins) with 0.1% missing values. Raw signals of each peptide and protein in each sample were curated with an expert system. The NCI-60 human cancer cell line panel contains 60 lines from 9 different tissue types ^12^. The NCI-60 have been molecularly and pharmacologically characterized with unparalleled depth and coverage, offering a prime *in vitro* model to further our understanding of cancer biology and cellular responses to anti-cancer agents ^12, 13^. Discoveries enabled by the NCI-60 in recent years include the development of the FDA approved drugs oxaliplatin for the treatment of colon cancers ^14^, eribulin for metastatic breast cancers ^12^, bortezomib for the treatment of multiple myeloma ^15^, and rhomidepsin for cutaneous T-cell lymphomas ^16^. The sensitivity of the NCI-60 has been measured for over 100,000 synthetic or natural compounds derived from a wide range of academic and industrial sources ^12^, constructing the most comprehensive resource for cancer pharmacological research. The proteomic data complement the existing NCI-60 molecular landscapes, allowing systematic investigation of the complementarity among genomics, transcriptomics and proteomics in a number of applications.

The proteome of the NCI-60 cells has been analyzed previously by data dependent analysis (DDA), a commonly used discovery mass spectrometry technique ^17^. Whereas the study reported the cumulative identification of 10,350 IPI proteins from about 1,000 fractionated and kinase-enriched sample runs, only 492 IPI proteins were quantified across the NCI-60 cell lines without missing value. The present study thus extends the number of consistently quantified proteins, in duplicates, to 3,171, with a ca. six-fold increase. The high quality proteomic data were used for pharmacoproteomic analysis of the response of the cell panel to 240 anti-cancer drugs, resulting in the identification of novel proteomic drug response determinants for clinically relevant chemotherapeutic and targeted therapies.

## Results

### Acquisition of the NCI-60 proteome maps

We applied the PCT-SWATH workflow ^7^ to generate quantitative proteome maps of the NCI-60 cell lines in technical replicates, resulting in the generation of 120 SWATH maps with high reproducibility at the raw data level (**Supplementary Fig. 1**). The PCT-assisted sample preparation took about 18 working days and the SWATH-MS analyses consumed about 12 working days. Thus, the entire process, from sample preparation to data acquisition, was accomplished within 30 working days, at an unprecedented sample-throughput compared to other cancer proteomic research of comparable scale ^4–6, 17^, which is due to the elimination of multiple dimensional fractionation, using one barocycler and one mass spectrometer (**Supplementary Fig. 1**, **Supplementary Table 1**).

SWATH proteome maps contain fragment ion chromatograms from all MS-measurable peptides, albeit in a highly convoluted form. To interpret the SWATH maps, we built a human cancer cell line spectral library containing 86,209 proteotypic peptides, *i.e.* peptides that uniquely identify a specific protein, from 8,056 SwissProt proteins (**Supplementary Table 1**). Using this library and the OpenSWATH software ^11^, we identified 6,556 protein groups, covering 81% of the library (**Supplementary Fig. 2**). To avoid ambiguity of peptide/protein quantification, we limited our analyses to canonical and proteotypic peptides and proteins by excluding protein isoforms, un-reviewed protein sequences, and peptide/protein sequence variants.

We evaluated the technical variation of each measurement through manual inspection of the OpenSWATH results based on the replicated measurement for each cell line and observed in multiple cases substantial technical variation. This is probably due to the fact that cell type-specific interfering signals leads to invalid SWATH assays, and the presence of irregular liquid chromatography (LC) and MS behavior of certain peptides in the highly variable proteomic context of the NCI-60 cells. These phenomena have also been observed previously in selected reaction monitoring (SRM)-based targeted proteomics studies ^18^.

To obtain high accuracy quantitative data for the cell lines, we further developed an expert system, *i.e.* DIA-expert (see Methods), to refine the peptide identification and quantification provided by automated analysis tools like OpenSWATH (Fig. 1A). We thus excluded proteins and peptides that were not reproducibly quantified in technical replicates and focused our analyses on a shorter list of 22,554 proteotypic peptides from 3,171 proteins, with 8% missing values at the peptide level and 0.1% missing values at the protein level across all MS runs (**Supplementary Table 1**). On average, 7 peptide precursors and 6 unique peptide sequences were identified for each protein (Fig. 1B). Several proteins were identified with over 200 peptides (Fig. 1C). The proteins excluded by DIA-expert may not be incorrect identifications, but rather proteins that could not achieve reproducible quantification by the existing algorithm across all cell lines due to either technical issues, for instance the signal-to-noise ratio, or biological issues such as post-translational modifications and splicing variants. Improved computational methods will likely rescue some of them in the future.

**Figure 1.**
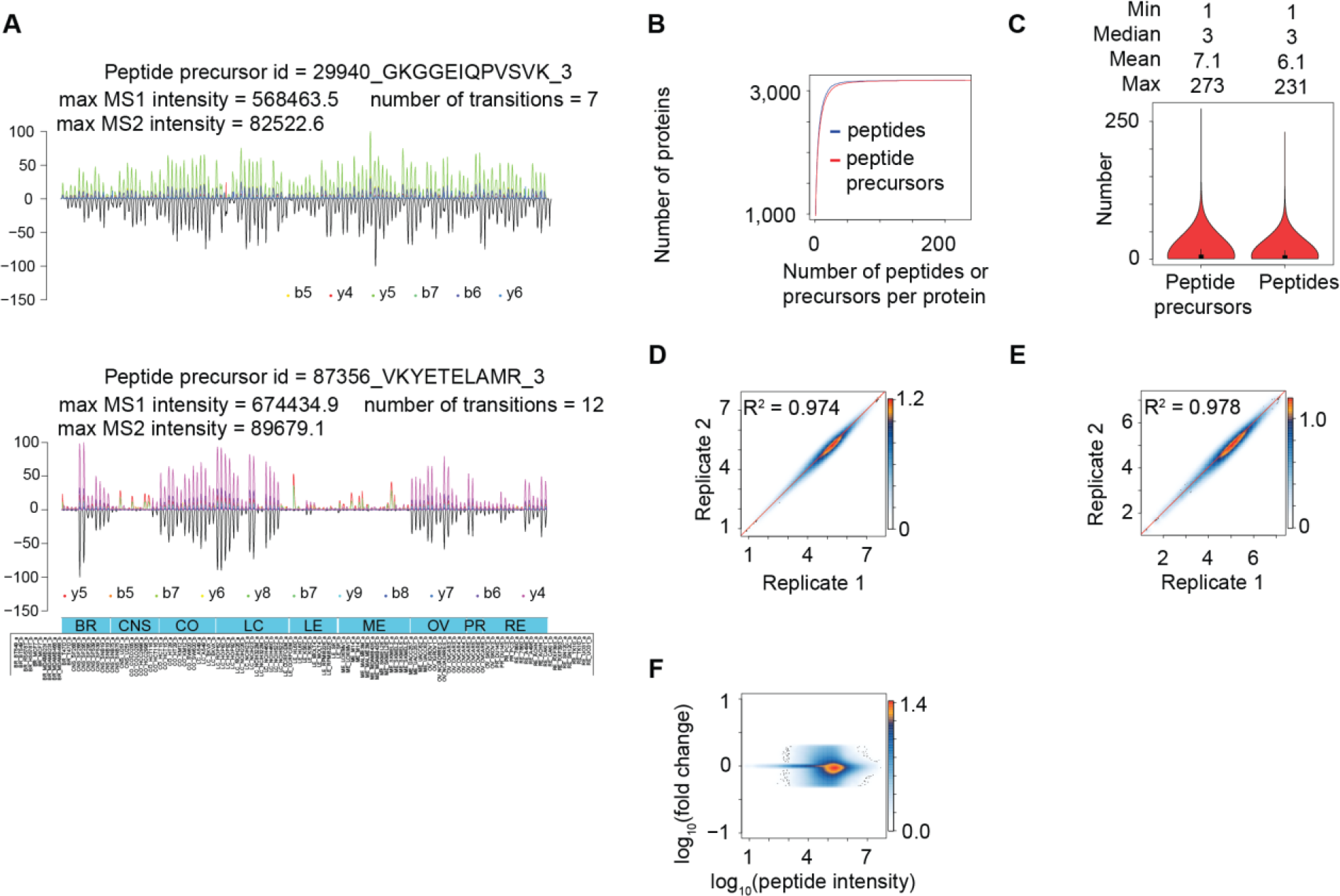
The acquisition of NCI-60 proteotype. (**A**) Representative peptide signals as curated and visualized by the DIA-expert software. (**B**) The cumulative number of peptide and peptide precursors identified for each protein. (**C**) The distribution of peptide precursors and peptides per protein. The overall Pearson correlation between technical replicates at the peptide level (**D**) and the protein level (**E**). Here, the log10 transformed intensity of each peptide/protein in each cell line technical replicate is plotted in the heatmap. (**F**) Dynamic range of the MS signals for 22,968 proteotypic peptides.

Most peptides for the 3,171 proteins were consistently quantified in all cell lines at both MS1 and MS2 levels. Two representative peptides are shown in Fig. 1A. The coefficient of determination (R^2^) between technical replicates, for overall expression of peptides (Fig. 1D) and proteins (Fig. 1E), were 0.974 and 0.978, respectively, with a dynamic range over 5 orders of magnitude (Fig. 1F). We provide the raw MS signals for each quantitative value in **Supplementary File 1**, allowing visual inspection of the MS signal for every peptide in each sample. When we limited the minimal peptide number per protein to 2, 3 and 4, fewer proteins were quantified however the quantitative accuracy did not substantially improve (**Supplementary Figure 3**).

### Characterization of the NCI-60 quantitative proteomes

The landscape of the 120 thus measured proteotypes is displayed in Fig. 2A. All technical replicates were clustered together using an unsupervised method based on the quantified proteotypes, confirming high quantitative accuracy. In most cases, the proteotypes are not strikingly different across different cancer cell lines, in sharp contrast with the distinct proteomes of tumor versus non-tumor kidney tissues ^7^. The median coefficient of variation (CV) of the protein intensity in different cells was 48%. The CV demonstrated a low dependence on protein abundance, as evident from the distribution of its values for different expression level quantile groups of the measured proteins (Fig. 2B). We then compared our data with the previously reported proteome of the NCI-60 cells using DDA-MS ^17^. While the DDA data reported comparable number of IPI protein groups to the SwissProt proteotypic protein number from this SWATH data set, the SWATH data exhibited much higher degree of consistency (**Supplementary Table 2 and Supplementary Fig. 5**) and better quantitative accuracy (**Supplementary Fig. 6-7**).

**Figure 2.**
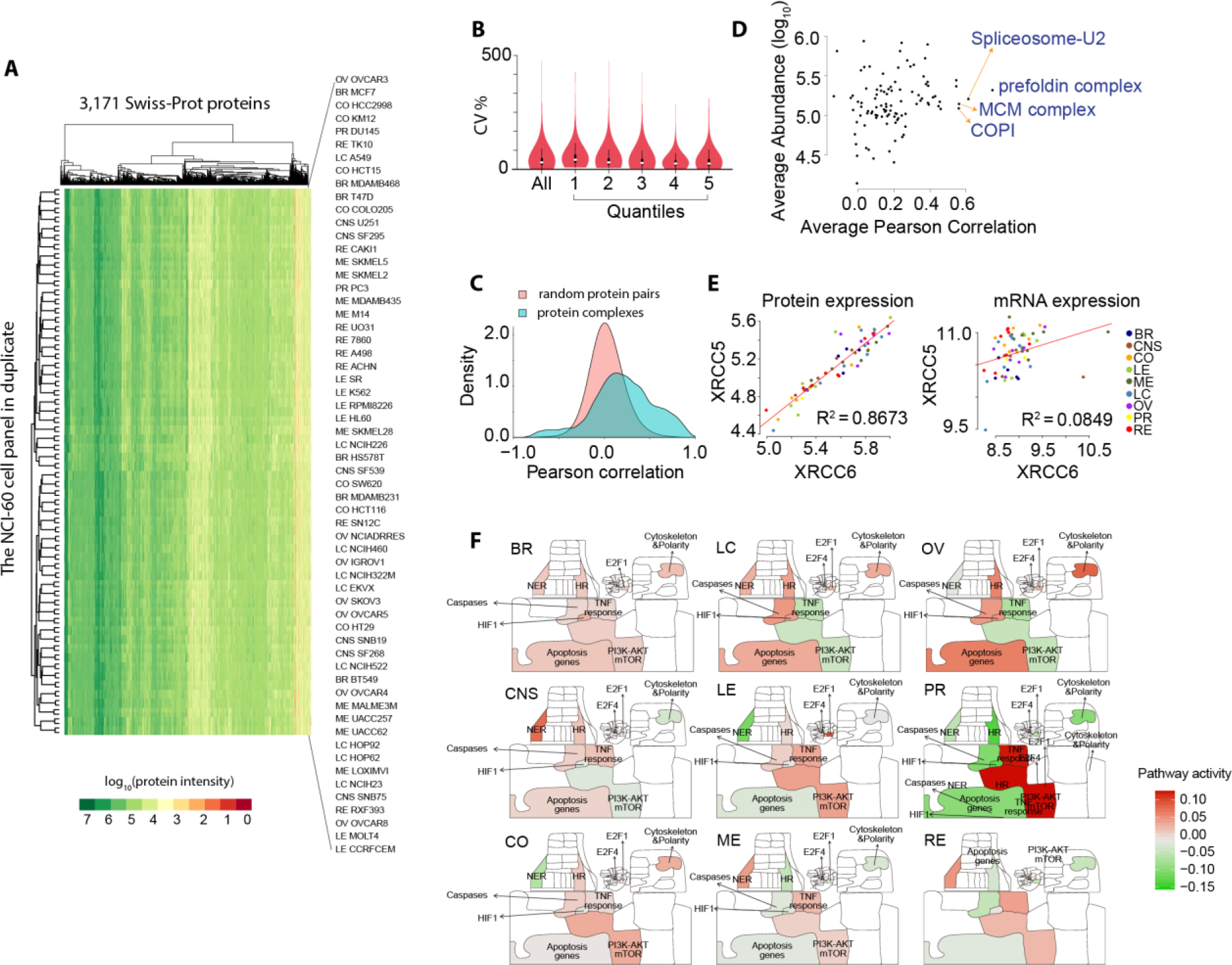
Characterizing NCI-60 quantitative proteomes. (**A**) Heatmap overview of NCI-60 proteotype data matrix. 3,171 Swiss-Prot proteins were quantified in 120 SWATH runs. (**B**) Variation of protein expression, for all proteins (All) and proteins in each abundance quantile group (from low abundance to high abundance). (**C**) Density plot of correlation of determination between pairs of random proteins versus pairs of proteins within a complex. (**D**) Stoichiometry variation of protein complexes in the NCI-60 cells. The x-axis shows the average Pearson correlation of each protein complex across the NCI-60. The y-axis shows the average abundance of proteins in a complex. Stable complexes tend to show higher values of average Pearson correlation. (**E**) Protein and mRNA expression of XRCC6/Ku70 and XRCC5/Ku80. (**F**) Visualization of pathway activity in NCI-60 proteotypes. More detailed pathway annotations for this Google map are provided in **Supplementary File 2**.

### Quantification of drug-responsiveness related proteins

The proteotypes covered 105 protein targets for FDA-approved anti-cancer compounds, 661 protein drug targets annotated in DrugBank ^19^ (including 68 drug metabolizing enzymes, 5 drug carriers, and 15 drug transporters), 694 proteins known to participate in human diseases ^19, 20^, and 58 human protein kinases, in addition to proteins involved in various biological functions (**Supplementary Table 3**). Some kinases were found to be broadly expressed in most cells with high abundance, including MST4 and WNK1 (**Supplementary Fig. 4**), consistent with previous reports ^21, 22^. Other kinases were highly expressed in specific cell lines, for example, EGFR in the breast cancer cell line MDAMB468, ERBB2 in SKOV3 cells, and CDK6 in MOLT4 cells, in agreement with previous studies using antibody-based methods ^20, 23^.

One unique benefit of our proteomic data set, compared to genomic and transcriptomic data, is its capacity to reveal more accurate information about the abundance of protein complexes and their stoichiometry ^24^. Our measurements included 101 protein complexes comprising 1,045 proteins (**Supplementary Table 4**) from a curated resource ^24^. Significantly higher Pearson correlation coefficients for pairs of proteins that are part of a complex further supported the quantitative accuracy of our data matrix (Fig. 2C). We applied our computational pipeline for analyzing co-expression of protein complex numbers ^24^ to the NCI-60 proteotype data and confirmed conserved stoichiometry of protein complexes such as the prefoldin and MCM complexes in various cell lines (Fig. 2D). In a specific case, we observed a high correlation between the protein expression of XRCC6/Ku70 and XRCC5/Ku80, a critical heterodimer involved in DNA repair and responsible for resistance to radiotherapy and chemotherapy. Ku80 is degraded when not bound to Ku70 ^25, 26^. Remarkably, this correlation is not detectable using mRNA measurements (Fig. 2E), indicating that expression of Ku80 is tightly regulated by protein degradation mechanisms independent of cancer types. Indeed, a recent report has shown that RNF8, an E3 ubiquitin ligase, regulates the expression of Ku80 via its removal from DNA double strand break sites and its degradation through ubiquitination ^27^.

### Google-map-based visualization of cancer signaling pathways

The NCI-60 proteotypes cover 648 proteins in the Atlas of Cancer Signaling Networks (ACSN), a manually curated pathway database presenting published facts about biochemical reactions involved in cancer using a Google-Maps-style visualization (**Supplementary Fig. 8**) ^28^. When mapping the mean protein expression per cancer type, we found that multiple pathways in different cell types, including apoptosis, cell survival, motility and DNA repair among others, displayed a similar pattern (**Supplementary File 2**), consistent with the fact that the immortal cells retain cancer hallmarks after artificial culturing ^29^. An example of a clear proteotypic pattern is the delta isoenzyme of protein kinase C, *i.e.* PRKCD, involved in DNA repair and a drug target that has been tested in various cancers ^30^. It was reported to be absent in four renal clear cell carcinoma lines ^31^. In agreement, this protein stood out in our visualization, with significantly lower protein expression in renal carcinoma, relative to the average expression in the NCI-60 panel. We provided detailed instructions on how to navigate through the atlas and explore protein abundance in each cancer cell line (see **Supplementary File 2)**.

We next compared the activity of cellular pathways using ROMA (Representation and quantification Of Module Activities) ^32^ (Fig. 2F), a gene-set-based quantification algorithm. This approach revealed substantial diversity of pathway activity between different proteotypes as evidenced by two-tailed *t*-tests of activity scores (*P*-value < 0.05). When mapping activity scores onto ACSN, some tissue specificities were revealed, with particular cell line proteotypes displaying distinct patterns of pathway activity. For instance, the activity of apoptosis (with both Caspases and Apoptosis Genes modules) was found to be significantly higher in ovarian cell lines (see **Supplementary Table 5**). Although there are only two prostate cancer cell lines in the panel, our analysis was able to highlight modules including “AKT-mTOR” and “Apoptosis”, whose differential activity can be attributed to HSP90AA1 and PRDX. The latter protein has been independently reported to be overexpressed in prostate tumors ^33^.

### Accessibility of the NCI-60 proteotypes

To enable easy data access, visualization, and comparison with other NCI-60 data sets, we have incorporated the SWATH data into the CellMiner database ^13, 34^. CellMiner allows the direct download of the data, as well as comparative and integrative analyses with other molecular data and pharmacological data, *e.g.* sensitivity of each cell line to over 20,000 compounds, and the manual inspection of specific genes, up to 150 per query. The detailed instructions for using this resource are provided on the project website (https://discover.nci.nih.gov/cellminer/) and in **Supplementary Fig. 9**. We have also deposited raw data and processed data matrices of the NCI-60 proteotype in public databases, including PRIDE ^35^ and ExpressionArray ^36^.

### Predicting drug responsiveness

The robust, quantitative proteomic data, with almost no missing values, permitted systematic investigation of whether integration of the SWATH-based proteotype with existing genomic and transcriptomic features improves the prediction of drug responsiveness (**Supplementary Table 6**). We generated various combinations of molecular features, and evaluated their predictive power using the Pearson correlation between predicted and observed drug response values for 240 FDA-approved or investigational compounds in CellMiner ^13, 34, 37^. Each compound is assigned a NSC (National Service Center) identifier upon submission to the National Cancer Institute for evaluation in the NCI-60 panel. The largest groups of drugs with target annotations are those that interfere with DNA synthesis and the DNA damage response, including topoisomerase inhibitors. The drug set also contains dozens of targeted agents, including 18 serine and threonine kinase inhibitors and 18 tyrosine kinase inhibitors (Fig. 3A).

**Figure 3.**
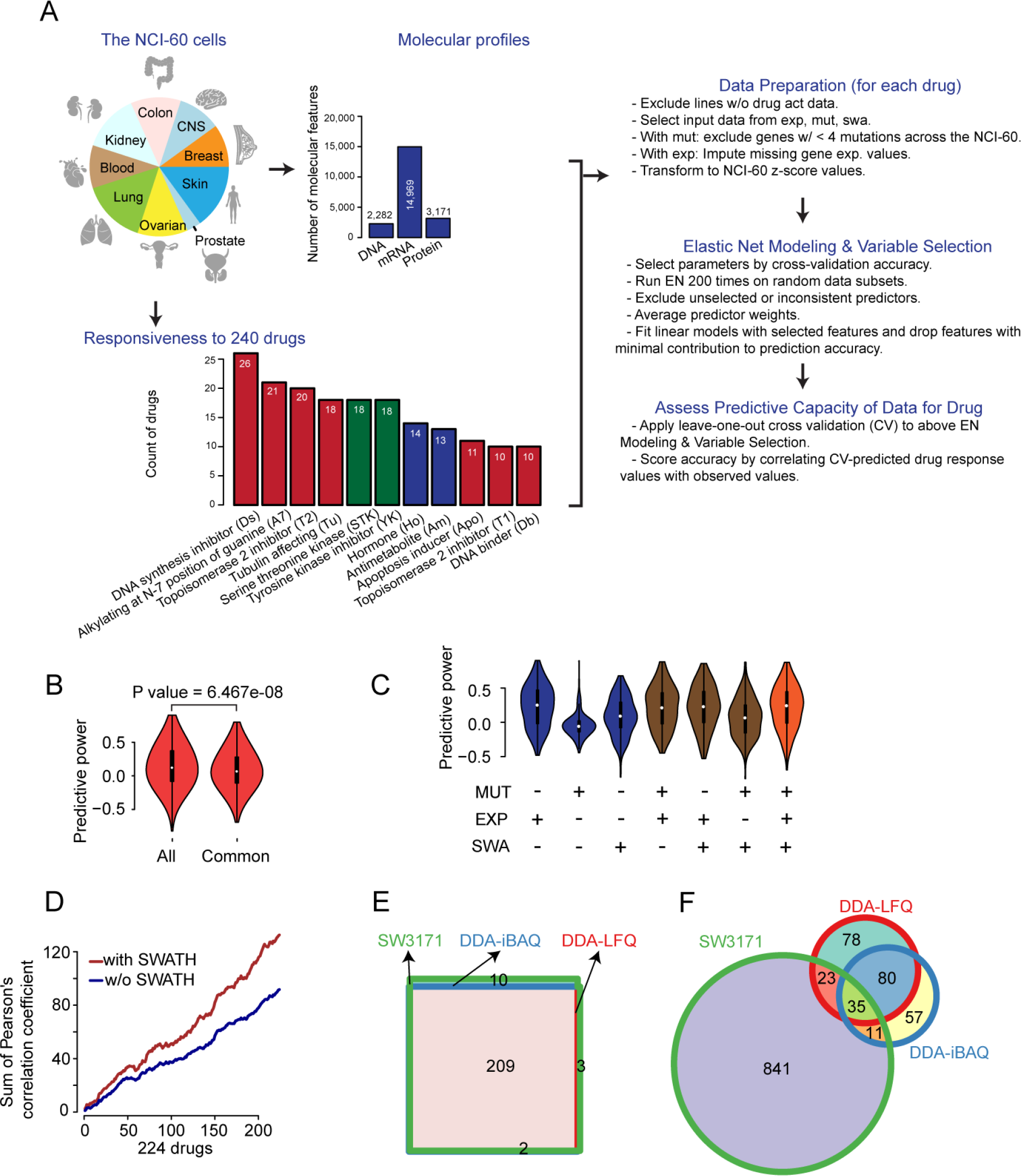
Prediction of drug responsiveness. **(A)** Workflow for drug responsiveness prediction. Drug groups with at least ten drugs are shown. (**B**) Distribution of predictive power (Pearson’s correlation of cross-validation predicted vs. observed response) for 240 compounds using all molecular features (All) versus common features (Common) available for all molecular data types. (**C**) Distribution of predictive power for different molecular data sets and their combinations. (**D**) Cumulative sum of Pearson correlation coefficients from drug responsiveness prediction in 224 drugs. (**E**) Venn diagram of drugs successfully modeled using elastic net using our SWATH data containing 3171 proteins (SW3171), and the DDA data based on iBAQ (DDA-iBAQ) and LFQ (DDA-LFQ). (**F**) Venn diagram of protein predictors using the SWATH and DDA data sets.

Using the elastic net algorithm, we then developed multivariate linear models to predict the NCI-60 response for each compound based on genomic, transcriptomic and proteomic features. The Pearson’s correlation between observed drug response values and leave-one-out cross validation-predicted response values was applied to evaluate the performance of each predictive model.

As different numbers of features were measured for each omics data set, two strategies were adopted in the modeling analyses. First, we used all omics features (2,282 DNA mutations, 14,969 mRNAs and 3,171 proteins), separately and combinatorically, as inputs to evaluate the general performance. Second, we selected 1,566 features that were available for all three molecular data types (denoted as common features). In both cases, we obtained valid models for 224 (93%) of the drugs. The predictive power achieved with all features was slightly higher than that obtained using the common features for all three data types (Fig. 3A); a likely reason for this is that the latter excluded some genomic and transcriptomic features not detected at the protein level. We accordingly derived our main analysis results from data including all available molecular features. Our modeling led to the discovery of valid biomarkers for drug responsiveness prediction. For instance, we found that the mRNA expression of SLFN11, strikingly responsible for the sensitivity of 45 compounds, out of which 39 were FDA-approved drugs including topoisomerase inhibitors, alkylating agents, and DNA synthesis inhibitors, was the most dominant indicator, in agreement with our previous report ^38^ (**Supplementary Table 7**). Fourteen ATP-binding cassette family transporters, detected as mutation, transcript or protein levels, were found responsible for sensitivity prediction of 51 compounds including chemotherapeutic agents and protein-targeting agents such as HDAC inhibitor Depsipeptide, HSP90 inhibitor Alvespimycin, mTOR inhibitor Temsirolimus and BCR-ABL inhibitor Nilotinib (**Supplementary Table 7**).

For ease of reproducibility of data analysis, we developed a Docker container (described in **Methods**) that includes our code and other essential dependencies, allowing all analyses to be replicated and extended for this and other omics data sets.

### Synergies among mutations, transcripts and proteins

Our pipeline led to the identification of valid models for 224 compounds (**Supplementary Table 8**). Given the relatively small sample size, it was not surprising that accurate predictive models could not be found for every drug, particularly those with limited numbers of responsive lines. We found that the SWATH-MS derived proteotypes displayed higher percentage of predictive features than mutations and transcripts. 1,090 (34%) out of 3,171 SWATH features are predictive, while 284 (12%) out of 2,282 features for mutations and 1,976 (13%) out of 14,969 transcripts were selected in the models. In general, the SWATH data outperformed the mutation data, however, the mRNA expression data set has about a five to six-fold higher number of features than the protein and mutation data sets (Fig. 3A) and exhibited better overall performance (Fig. 3C).

Our analyses revealed notable synergies among the different molecular measurements. Each type of molecular data set demonstrated indispensable benefits in predicting the response to certain drugs/compounds. The responsiveness of 35 compounds (16%) out of 224 was best predicted with SWATH data, whereas 107 compounds (48%) were best predicted by SWATH data or by combining SWATH data with transcripts and/or DNA data. The most accurate models for over half of the compounds required at least two different types of molecular features. We then computed accumulative sum of Pearson correlation coefficient based on drug responsiveness prediction and observed significant contribution of SWATH data (Fig. 3D). We also compared the predictive power of the DDA data to the SWATH data. While the DDA data were able to generate elastic net models for comparable number of drugs (Fig. 3E), the number of protein predictors is much lower than SWATH data over some overlap (Fig. 3F).

### Drug responsiveness prediction

Based on the integration of various data sets, global drug response patterns were predicted for the 158 well-modeled drugs (Fig. 4, see Methods), with predictive molecular features for individual compounds provided in **Supplementary Table 8**. The data generated from this computational pipeline were validated by the recovery of established pharmacogenomic knowledge. For instance, the mutational status of BRAF was the top predictive molecular feature for sensitivity to BRAF inhibitors, *e.g.* vemurafenib (NSC 761431) and dabrafenib (NSC 764134), and this association was particularly evident in melanomas. Activated BRAF mutational status also sensitized cells to the MEK inhibitor hypothemycin (NSC: 354462), as has been previously described ^39^.

**Figure 4.**
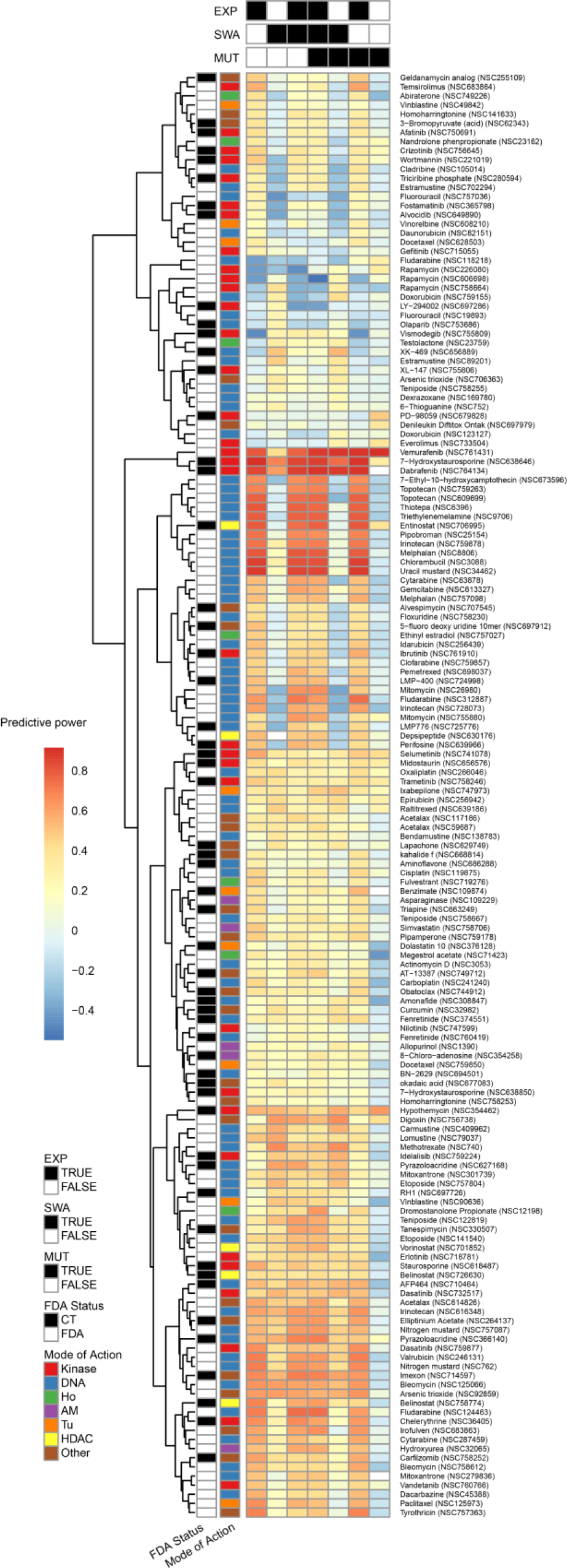
Predictive power for 224 compounds using different types of omics data. We applied elastic net and cross validation to evaluate the drug response predictive accuracy for each omics data set and combinations of data sets for 224 drugs which could be effectively modeled. Drug response prediction accuracies across input data types are clustered without supervision. MoA of compounds and clinical status of the compounds are colored. Each column indicates an input data type or combination of types; each row represents a compound. The color indicates the predictive power measured by Pearson correlation of cross-validation predicted versus observed drug response values. Black indicates that a valid elastic net model could not be obtained.

Sensitivity to the antimetabolite 6-thioguanine (6-TG, NSC: 752) (Fig. 5A) was predicted by protein expression of NUDT5 and MAT2B within an elastic net model composed of 5 proteomic features: NUDT5, MAT2B, CD47, STX12 and GFAP. The cross-validation accuracy with this compound and the SWATH-MS data was relatively low (r = 0.27), probably due to instability in the selected predictive features with limited sample size. Still, we find that for the two strongest predictors in the model, NUDT5 and MAT2B, the expression data were significantly correlated with the activity of 6-TG (Fig. 5B and 5C). Additionally, we were able to relate the inter-connected activities of these two proteins to the mechanism of action for 6-TG. In the purine salvage pathway, HPRT1 catalyzes synthesis of inosine monophosphate from hypoxanthine and phosphoribosyl pyrophosphate (PRPP), with production of the latter stimulated by NUDT5. 6-TG can substitute for hypoxanthine, ultimately yielding altered nucleotides that are toxic upon incorporation into DNA ^40^. PRPP is still required, so low NUDT5 expression could possibly induce 6-TG resistance. This is consistent with our NCI-60 data and recent experimental work showing that depletion of NUDT5 confers resistance to 6-TG ^41^. As noted in Fig. 5A, a metabolite of 6-TG, thioguanosine monophosphate (TGMP) can be inactivated by methylation. Production of the methyl group donor, S-adenosylmethionine (SAMe), is catalyzed by the methionine adenosyltransferase IIα (MAT2A) enzyme. The MAT2B protein, exhibiting high correlation with MAT2A (Fig. 5D), is a regulatory component of MAT which may enhance feedback inhibition by SAMe ^42^. Increased MAT inhibition and diminished TGMP methylation may shunt more TGMP toward DNA incorporation, enhancing the 6-TG response. In spite of its relatively low cross-validation accuracy, the presented model may provide a starting point for further exploration, in light of the supporting prior research.

**Figure 5.**
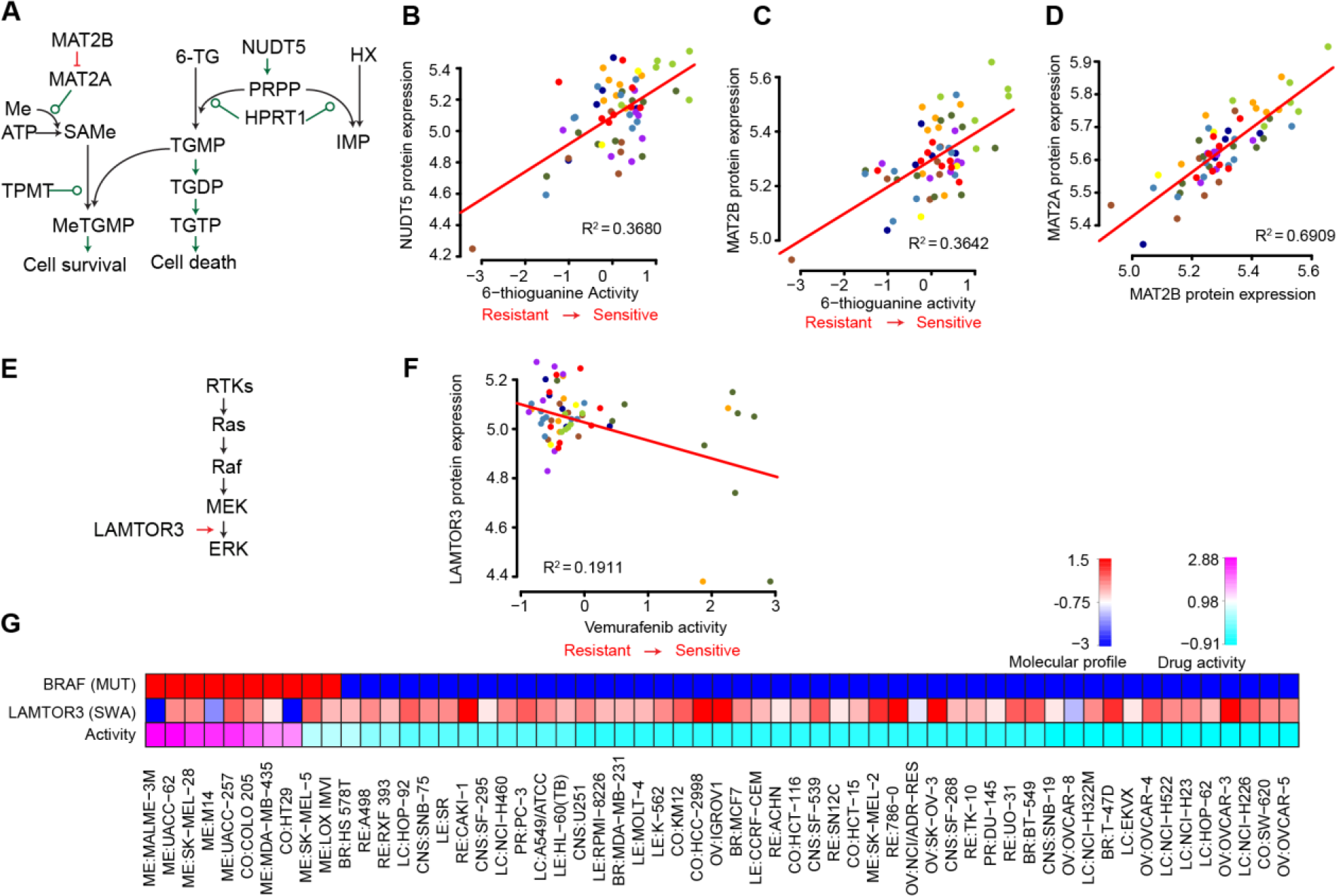
Drug responsiveness predicted by SWATH data. **(A)** molecular mechanisms of 6TG. **(B)** correlation between NUDT5 protein expression and 6-TG activity. **(C)** correlation between MAT2B protein expression and 6-TG activity. **(D)** correlation between MAT2B and MAT2A protein expression. **(E)** LAMTOR3 facilitates MEK/ERK pathway activation by binding MEK and ERK. **(F)** correlation between LAMTOR3 protein expression and Vemurafenib activity. **(G)** Association of BRAF mutation and LAMTOR3 protein expression with Vemurafenib activity.

Analysis of the protein kinase inhibitor vemurafenib (NSC 761431) yielded a multivariate model based on BRAF V600E activating mutation status ^43^ and the protein expression level of LAMTOR3. LAMTOR3 (MP1) is part of an endosomal scaffolding complex that interacts with components of the RAF/MEK/ERK mitogenic signaling pathway (Fig. 5E). In particular, LAMTOR3 binds MEK1 and ERK1, facilitating activation of the latter protein ^44^. Elevated LAMTOR3 protein expression was correlated with vemurafenib resistance (r= 0.44, Fig. 5F), consistent with the hypothesis that LAMTOR3 has the capacity to enhance RAF/MEK/ERK pathway signaling downstream from RAF. In particular, increased protein expression of LAMTOR3 was observed in two BRAF mutant cell lines, ME:SK-MEL-5 and ME:LOXIMVI, which are relatively resistant to Vemurafenib (Fig. 5G). Due to the limited number of BRAF mutant cell lines exhibiting relative drug resistance (*i.e.* 2 cell lines), additional statistical analyses with sufficient power are not possible. Robust statistical validation of this model may possible when larger cell line databases (e.g. the Sanger and Broad resources) expand to include proteomic coverage of LAMTOR3. Still, this finding remains relevant in light of the recent research into the activity of LAMTOR3, including the observation that reduced LAMTOR3 protein levels decreased the activation of MEK1/2 and ERK1/2 ^44, 45^. Additionally, LAMTOR3 has been shown to affect proliferation of pancreatic and breast cancers ^46, 47^, and has been patented as a diagnostic biomarker for breast cancer ^47^.

Our elastic net analysis also produced multiple recurrent predictors with plausible drug response associations. ABCC4 was a negatively weighted predictor for several alkylating agents, including chlorambucil (NSC: 3088), uracil mustard (NSC: 34462), nitrogen mustard (NSC: 762), consistent with its established role as a drug efflux pump ^48^. Another recurrent, negatively-weighted predictor was CTNND1, which was identified for several compounds, including bendamustine (NSC: 138783), etoposide (NSC: 141540), valrubicin (NSC: 246131), and carmustine (NSC: 409962). CTNND1 encodes delta-catenin, whose overexpression promotes cell survival through activation of Wnt pathway signaling ^49^. The resulting inhibition of apoptosis ^50^ could plausibly confer resistance to the mentioned DNA-damage inducing drugs.

## Discussion

Due to the complementarity of protein and transcript data ^4–6, 51^, it can be expected that the rapid and consistent quantification of thousands of proteins across a large sample cohort will revel new biological information that is not apparent from the commonly used transcript profiles. However, due to technical limitations, such proteomic cohort datasets have been challenging to acquire. Here, using the NCI-60 cell line compendium, we demonstrate the ability of the PCT-SWATH proteomic technique to consistently quantify in excess of 3000 proteins across the 60 cell lines measured in duplicate. The data were acquired in 30 working days on a single mass spectrometer and for each sample measurement ca. 1 microgram of total peptide mass was consumed. This has been enabled by the pressure cycling technology which minimizes samples consumption and the data-independent MS data acquisition using SWATH-MS ^7^. The data generated and their use to reveal cancer biology and drug response determinants represent a significant advance in the field.

The proteome of the NCI-60 cells has been previously measured by extensive sample fractionation and DDA-MS analysis of over 1,000 fractionated samples ^17^. In this study, data acquisition for each cell line required an average of about 29.16 hours MS instrument time. That shotgun proteomics study reported the cumulative identification of 10,350 IPI proteins over the NCI-60 cell lines. However, only 492 proteins were quantified in all cell lines without missing value. The PCT-SWATH methodology adopted in this study offers an over 10-fold increase in sample-throughput, which has allowed us to acquire the proteotype for each cell in the NCI-60 panel in duplicate, with standardized sample preparation, within 30 working days. In addition, our data have 0.1% amount of missing values at protein level owing to the data acquisition strategy and improvements in bioinformatics analysis. This study demonstrates that the human proteotype can be obtained with a throughput comparable to genomic and transcriptomic analyses, though still at relatively lower coverage.

Two aspects of our workflow ensure robust and quantitatively accurate protein expression measurements. First, we obtained technical duplicates for the entire set of NCI-60 proteotypes, which was feasible due to the unparalleled high sample-throughput of the PCT-SWATH methodology which is now gaining popularity in proteomic profiling of clinical specimens. In addition, we developed an expert system software (manuscript in preparation) to further curate peptide and protein identification and quantification. Applying stringent criteria, 3,171 proteins were included for further analyses. The raw MS signal for each of these quantified proteins, in each cell line, was inspected by the expert system, simulating manual inspection, and is available for visual inspection in the supplementary data. We further compared the expression of a few proteins with known expression in certain cell lines, obtaining good agreement. Nevertheless, we cannot conclude that the peptides and proteins that failed to pass curation by the expert system are not biological signals, due to the unpredictable degree of biological heterogeneity, and the fact that we did not analyze non-canonical peptide variants and post-translational modification. The latter can be potentially dissected and quantitated by future *in silico* analyses of our SWATH maps. Since the NCI-60 cell lines are widely used in cell biology, we anticipate broad utility of this highly curated proteomic data. Additionally, our rapid proteotype acquisition pipeline using PCT-SWATH requires little biological material, making it suitable for clinical settings and in precision medicine efforts ^7, 8, 52^.

Compared to other omics data, the proteotypes obtained here offered unique insights into the coordinated expression of protein complexes. Interactions amongst their component subunits contribute to our understanding of protein function, as well as human diseases ^24, 53–55^. Several protein complexes have been identified as biomarkers of disease, including cancer progression ^56^. Our high quality proteomic data allowed systematic investigation of the composition of 101 protein complexes in 60 cell lines. We expect that this represents a proof-of-principle for a generic, high-throughput approach, applicable to clinical specimens ^7^, for exploring the association between protein complexes and biological/disease phenotypes.

The NCI-60 continues to enable important contributions that have come and continue to come from this resource, and often emerging technologies are first tested on this cell line panel due to its diversity and depth of surrounding knowledge ^3, 12, 57–59^. Each cancer cell line in the NCI-60 has been tested against tens of thousands of compounds, including the 240 FDA-approved and investigational drugs featured in our analyses. With the addition of the SWATH proteomic data, the NCI-60 remains positioned as one of most comprehensive models for cancer research and drug discovery ^12, 15^. It uniquely enabled our thorough, integrative analysis of different molecular profiles (genomic, transcriptomic, and proteomic) in predicting drug responsiveness. Our findings strengthen the body of work highlighting the importance of integrative omic approaches in understanding drug mechanisms and establish the benefit of large-scale proteomic measurements. Therefore, we expect this work to become a seminal work in the area of pharmacoproteomics, the benefit of which will grow with anticipated expansion of sample size, proteomic coverage, including extension to phosphoproteomic expression, as well as extension to mouse models ^60^ and human specimens ^7^.

The existing SWATH data specifically enabled the use of advanced analysis techniques to produce multivariate models of drug response. Great effort was put into making our work accessible to a large audience through data submission to the NCI-60 CellMiner database and availability through an accompanying R package, rcellminer. We expect this pipeline based on the widely used elastic net method will continue to evolve and enable future studies on additional data sets and phenotypes. And while the strengths of the elastic net method over other related methods have been previously described ^61, 62^, the resulting models still require careful scrutiny by individual researchers. The interpretation of the models developed here, and by others using our pipeline, should be guided by understanding of the biological activities of the associated predictors in the context of the mechanisms of action for the input drugs. From the models generated by the current analyses, we identified several potential determinants of drug responses, including NUDT5 and MAT2B protein levels for the antimetabolite 6-TG, as well as complementary markers, such as LAMTOR3 protein levels in conjunction with BRAF mutational status for Vemurafenib and other BRAF inhibitors. These determinants may provide clinically relevant insights toward understanding mechanisms of resistance to these and other agents. Together, these results invite further investigation of this unique proteomic data resource. For example, in the current study’s analysis of protein complexes, we identified discrepancies between data at the transcriptomic and proteomic levels. This observation has been similarly made in tumor samples, with additional variation across tissue types ^63^. These differences can be used in future studies to develop drug response models with non-redundant predictor sets including both data types. However, due to the tissue diversity of the NCI-60 cells and the limited number of cell lines, data from more cancer cell lines of specific tissue type and extension to clinical specimens are required to advance our findings to clinical applications.

## Acknowledgements

We thank Margot Sunshine who developed CellMiner and the NCI-DTP team (Dr. Jerry Collins and Dr. James H. Doroshow) for the drug data and support for data acquisition, Emanuel Gonçalves for comments to the manuscript. The work was supported by the SystemsX.ch project PhosphoNetX PPM (to R.A.), the Swiss National Science Foundation (grant no. 3100A0-688 107679 to R.A.), the European Research Council (grants no. ERC-2008-AdG 233226 and ERC-20140AdG 670821 to R.A.), European Union’s Horizon 2020 research and innovation programme under grant agreement No 668858 (to R.A., J.S.-R., L.C., P.W.), the Ruth L. Kirschstein National Research Service Award (grant no. F32 CA192901 to A.L.), the National Resource for Network Biology (NRNB) from the National Institute of General Medical Sciences (NIGMS) (grant no. P41 GM103504 to C.S.), and the Center for Cancer Research, Intramural Program of the National Cancer Institute (grant no. Z01 BC006150 to Y.P), and the Wellcome Trust Award (102696) to M.J.G.. We thank An Guo for help in preparing the graphics.

## Authors contributions

T.G. designed and coordinated the project with supervision from R.A. C.C.K. processed the samples. L.G., C.C.K. and T.G. acquired the SWATH data. T.G. performed the SWATH data interpretation and benchmarking with help from C.C.K., and the expert system analysis with help from U.S. A.L., V.N.R. and Z.W. performed the drug response prediction analysis, and developed the reproducible research infrastructure, with critical inputs from M.P.M., J.S.R., M.J.G., S.V., W.C.R., C.S, and Y.P.. L.C. and L.M. performed the pathway analysis. A.L., V.N.R., W.C.R. and S.V. integrated the SWATH data into rcellminer and CellMiner. A.O., M.I. and R.C. performed the protein complex analysis, with help from A.L., Z.W., Y.C., V.N.R, C.S., Y.S., Y.Z., Y.P.. P.Q. and Q.Z. contributed to the data analysis. T.G., A.L. and V.N.R. wrote the manuscript with inputs from all co-authors. P.J.W., P.B., M.R., J.S.R., W.C.R., C.S., Y.P. and R.A. supervised the project.

## Materials and Methods

### PCT-assisted sample preparation for MS analyses

The NCI-60 cells were obtained as frozen, non-viable cell pellets from the Developmental Therapeutics Program (DTP), National Cancer Institute (NCI-NIH) and processed using Barocycler^®^ NEP2320 (PressureBioSciences Inc, South Easton, MA). The IDs of the NCI-60 cells in our study matching to the IDs in Cellminer and a previous proteomic study by the Kuster group are provided in **Supplementary Table 1**. Briefly, cell pellets were lysed in a buffer containing 8M urea, 0.1M ammonium bicarbonate, and Complete^TM^ protease inhibitor using barocycler program (20 seconds 45 kpsi, 10 seconds 0 kpsi, 120 cycles) at 35°C ^7^. Whole cell lysates were sonicated for 25 seconds with 1 min interval on ice for 3 times. Cellular debris was removed by centrifugation and sample protein concentration was determined by BCA assay prior to protein reduction with 10 mM TCEP for 20 min at 35°C, and alkylation with 40 mM iodoacetamide in the dark for 30 min at room temperature. Lys-C digestion (1/50, w/w) was performed in 6 M urea using PCT program (25 seconds 25 kpsi, 10 seconds 0 kpsi 75 cycles) at 35°C; whereas trypsin digestion (1/30, w/w) was performed in further diluted urea (1.6M) using PCT program (25 seconds 25 kpsi, 10 seconds 0 kpsi, 160 cycles) at 35°C. Digestion was stopped by acidification with trifluoroacetic acid to a final pH of around 2 before C18 column desalting using SEP-PAK C18 cartridges (Waters Corp., Milford, MA, USA).

### Off-gel electrophoresis

To create a comprehensive spectral library for SWATH-MS analysis, we pooled 20-40% of desalted peptide solutions from each NCI-60 sample and performed off-gel fractionation. Briefly, pooled peptides were resolubilised in OGE buffer containing 5% (v/v) glycerol, 0.7% (v/v) acetonitrile (ACN) and 1% (v/v) carrier ampholytes mixture (IPG buffer pH 3.0-10.0, GE Healthcare). Fractionation was performed on a 3100 OFFGEL (OGE) Fractionator (Agilent Technologies) using a 24 cm pH3-10 IPG strip (Immobilised pH Gradient strip from GE Healthcare) according to manufacturer’s instructions using a program of 1 h rehydration at a maximum of 500 V, 50 µA and 200 mW followed by separation at a maximum of 8000 V, 100 µA and 300 mW until 50 kVh were reached. Each of 24 fraction was recovered and cleaned up by C18 reversed-phase MicroSpin columns (The Nest Group Inc.). Based on the sample complexity (based on Nanodrop, A280 measurement), for each strip, the following fractions were pooled into 12 samples for MS injections: pool 1 (fraction 1-2), pool 2 (fraction 3), pool 3 (fraction 4), pool 4 (fraction 5), pool 5 (fraction 6-7), pool 6 (fraction 8-9), pool 7 (fraction 10-11), pool 8 (fraction 12-15), pool 9 (fraction 16-19), pool 10 (fraction 20-21), pool 11 (fraction 22), pool 12 (fraction 23-24). Those were injected in quadruplicate, resulting in 48 DDA injections of fractionated samples.

### DDA MS for spectral library generation

For spectral library generation, a SCIEX TripleTOF 5600 System mass spectrometer was operated essentially as described before ^64^: all samples were analyzed on an Eksigent nanoLC (AS-2/1Dplus or AS-2/2Dplus) system coupled with a SWATH-MS-enabled AB SCIEX TripleTOF 5600 System. The HPLC solvent system consisted of buffer A (2% ACN and 0.1% formic acid, v/v) and buffer B (95% ACN with 0.1% formic acid, v/v). Samples were separated in a 75 μm diameter PicoTip emitter (New Objective) packed with 20 cm of Magic 3 μm, 200A C18 AQ material (Bischoff Chromatography). The loaded material was eluted from the column at a flow rate of 300 nL min^−1^ with the following gradient: linear 2 - 35% B over 120 min, linear 35 - 90% B for 1 min, isocratic 90% B for 4 min, linear 90 - 2% B for 1 min and isocratic 2% solvent B for 9 min. The mass spectrometer was operated in DDA mode using a top20 method, with 500 ms and 150 ms acquisition time for the MS1 and MS2 scans respectively, and 20 s dynamic exclusion for the fragmented precursors. Rolling collision energy using the following equation (0.0625 × m/z - 3.5) with a collision energy spread of 15 eV was used for fragmentation regardless of the charge state of the precursors, to mimic as close as possible the fragmentation conditions of the precursors in SWATH-MS mode. Altogether, we had 66 DDA-MS injections, including the 48 OGE samples and another 18 pooled peptide samples from the unfractionated cell lysate of the NCI-60 cells.

### Spectral and assay library generation

All raw instrument data were centroided using Proteowizard msconvert (version 2.0). The assay library was generated using an established protocol ^64^. In short, the shotgun data sets were searched individually using X!Tandem ^65^ (2011.12.01.1) with k-score plugin ^66^, Myrimatch ^67^ (2.1.138), OMSSA ^68^ (2.1.8) and Comet ^69^ (2013.02r2) against the reviewed UniProtKB/Swiss-Prot (2014_02) protein sequence database containing 20,270 proteins appended with 11 iRT peptides and decoy sequences. Carbamidomethyl was used as a fixed modification and oxidation as the variable modification. Maximally two missed cleavages were allowed. Peptide mass tolerance was set to 50 ppm, fragment mass error to 0.1 Da. The search identifications were combined and statistically scored using PeptideProphet ^70^ and iProphet ^71^ available within the Trans-Proteomics Pipeline (TPP) toolset (version 4.7.0) ^72^. MAYU ^73^ (v. 1.07) was used to determine the iProphet cutoff (0.999354) corresponding to a protein FDR of 1.03%. SpectraST was used in library generation mode with CID-QTOF settings and iRT normal-isation at import against the iRT Kit ^74^ peptide sequences (-c_IRTirt.txt -c_IRR) and a consensus library was consecutively generated. An in-house python script, spec-trast2tsv.py31 (msproteomicstools 0.2.2) was then used to generate the assay library with the following settings: -l 350,2000 -s b,y -x 1,2 -o 6 -n 6 -p 0.05 -d -e -w swath32.txt -k openswath (fragment ions between 350 and 2000 m/z, b and y ions authorized, fragment charges 1+ and 2+, 6 most intense transitions, precision of fragment ion retrieved 0.05 Da, exact fragment ion mass calculated, exclude fragments in the swath window). The OpenSWATH tool, ConvertTSVToTraML converted the TSV file to TraML format; OpenSwathDecoyGenerator generated the decoy assays in shuffle mode and appended them to the TraML assay library. In this study, we built a SWATH assay library containing 86,209 proteotypic peptide precursors in 8,056 proteotypic SwissProt proteins. This library is supplied in PRIDE project PXD003539.

### SWATH-MS

The SWATH-MS data acquisition in a Sciex TripleTOF 5600 mass spectrometer was performed as described before ^10^, using 32 windows of 25 Da effective isolation width (with an additional 1 Da overlap on the left side of the window) and with a dwell time of 100 ms to cover the mass range of 400 - 1200 *m/z* in 3.3 s. The collision energy for each window was set using the collision energy of a 2+ ion centered in the middle of the window (equation: 0.0625 x *m/z* - 3.5) with a spread of 15 eV. The sequential precursor isolation window setup was as follows: [400-425], [424-450], [449-475], …, [1174-1200].

### Protein identification using OpenSWATH

We analyzed the SWATH data using OpenSWATH software ^11^ using parameters as described previously ^24^. We identified 48,374 peptides from 6,556 protein groups from the NCI-60 panel with < 1% false discovery rate at both peptide and protein level evaluated by OpenSWATH ^11^ and Mayu ^75^ (supplied in PRIDE project PXD003539).

### DIA-expert analyses

The DIA-expert software read OpenSWATH output result file which contains statistical scores (*i.e.* mProphet score or mScore) indicating the confidence of identification for each peptide precursor in each sample, and from there selected the sample in which a peptide precursor was identified with highest confidence. It then obtained extracted ion chromatograms (XICs) for the target peptide precursor and all associated annotated *b* and *y* fragments in the reference sample, and refined fragments based on the peak shape of each fragment and its peak boundary. The refined fragments and precursor XIC traces from each of the rest samples were subsequently compared with the reference peak group using empirical expert rules, based on which the best matched peak group in each sample was picked and visualized. Duplicated measurements were used to evaluate the accuracy of peptide and protein quantification. The protein quantity was normalized based on total ion chromatography of the MS1 spectra from each raw SWATH file. All codes are provided in Github https://github.com/tiannanguo/dia-expert.

### Protein complexes analysis

For this analysis, technical replicates were averaged to generate the NCI-60 proteotypes. To assess the coverage of protein complexes by NCI-60 proteotypes, we retrieved a large resource of mammalian protein complexes assembled from CORUM ^76^, COMPLEAT ^77^ and literature-curated complexes ^24, 78^. This resource contains 2,041 proteins as members of 279 distinct complexes and it is available at http://variablecomplexes.embl.de/. 101 complexes were represented in the NCI-60 proteotypes with at least 5 members quantified. These complexes, in total, contain 1,045 distinct proteins quantified in the NCI-60 proteotypes. Pearson’s correlation coefficient was calculated for all the pairwise comparisons of 3,171 proteins across the NCI-60 cell lines. All pairwise comparisons were classified into two categories: either two proteins were members of the same complex or not. Average abundance, standard deviation and average Pearson correlation of each complex were calculated based on the abundance of complex members in the NCI-60 proteotypes.

For this analysis, technical replicates were averaged to generate the NCI-60 proteotypes. To assess the coverage of protein complexes by NCI-60 proteotypes, we retrieved a large resource of mammalian protein complexes assembled from CORUM ^76^, COMPLEAT ^77^ and literature-curated complexes ^24, 78^. This resource contains 2041 proteins as members of 279 distinct complexes and it is available at http://variablecomplexes.embl.de/. 158 complexes were represented in the NCI-60 proteotypes with at least 5 members quantified. These complexes, in total, contain 1,045 distinct proteins quantified in the NCI-60 proteotypes. Pearson’s correlation coefficient was calculated for all the pairwise comparisons of 3,171 proteins across the NCI-60 cell lines. All pairwise comparisons were classified into two categories: either two proteins were members of the same complex or not. Average abundance and standard deviation of each complex were calculated based on the mean abundance of complex members in the NCI-60 proteotypes.

### Pathway activity analysis

The activity of pathways, as they are described in ACSN, has been computed using ROMA ^32^. Among all the modules defined in ACSN, only 11 show a significant dispersion over the data set: AKT_MTOR, HR (Homologous Recombination), NER (nucleotide Excision Repair), TNF response, Death Receptors regulators, Apoptosis, caspases, E2F3 and E2F4 targets, HIF1 and cytoskeleton polarity. For these modules, the mean activity score for each type of cancer cell lines was computed and mapped onto the atlas (from bright green for low values to bright red for high values). To assess module differential activity between proteotypes, we computed a *t*-test on the activity scores in cell lines of a cancer type versus the activity of all other cancer cell lines. The definition of genes composing each module can be found in http://acsn.curie.fr

### Drug sensitivity prediction using elastic net

The elastic net regularized regression algorithm was applied to predict drug response for 240 FDA-approved or investigational NSC-designated compounds. Some widely studied drugs are represented by more than one NSC identifier, with each identifier associated with a distinct compound sample and series of NCI-60 drug activity assays. For each compound, 7 combinations of input data were evaluated. These included NCI-60 mRNA expression, gene-level mutation, and SWATH-MS protein expression, both alone and in all possible combinations. mRNA expression data was available for 14,969 genes, and derived from CellMiner, with missing values imputed using the impute.knn function (with default parameters) of the Bioconductor impute package. Gene-level mutation profiles were available for 2,282 genes, and were obtained from CellMiner exome sequencing data, with values indicating the percent conversion to a variant form for the case of expected function-impacting alterations (frameshift, nonsense, splice-sense, missense mutations by SIFT/PolyPhen2 analysis). SWATH-MS based protein expression data was available for 3,171 proteins.

Elastic net analysis was done using the glmnet R package ^79^. The elastic net analysis was conducted using a multi-step pipeline involving cross-validations performed in a nested manner. The “outer” cross-validation is a leave-one-out cross validation that is conducted over all computational steps present in the “inner” pipeline, and it is used to validate model performance. The “inner” cross-validation are conducted to select elastic net hyperparameters (alpha and lambda) and for predictor set trimming, using data from a set of ∼59 cell lines.

The elastic net parameters alpha and lambda were selected by minimizing the cross-validation error (average of 10 replicates of 10-fold cross-validation) within the “inner” pipeline. The selected alpha and lambda parameters were then applied to 200 runs of the elastic net algorithm, each using a random data subset derived from 90% of the available cell lines. The 200 resulting coefficient vectors were then averaged, and predictors were ranked by the magnitude of their average coefficient weight. To select a limited number of predictors with potential to generalize to new data, top k-element predictor sets (by average coefficient weight magnitude) were evaluated using standard linear regression and 10-fold cross-validation. The appropriate k was set to the smallest value yielding a cross-validation error within one standard deviation of the minimum cross-validation error.

To obtain a robust estimate of performance on unseen data, leave-one-out cross-validation was applied to the overall procedure as part of the “outer” pipeline. Specifically, drug response for each cell line was predicted using an elastic net model derived using the remaining held out data (and the steps outlined above). The vector of predicted response values was then correlated with the actual response values, with the Pearson’s correlation coefficient providing an estimate of the predictive value of the applied input data combination. More details of the elastic net algorithm are provided in File S3.

Elastic net analysis was done using the rcellminerElasticNet R package (https://bitbucket.org/cbio_mskcc/rcellminerelasticnet), which facilitates the application of the glmnet R package (which provides the elastic net algorithm code) to data from the rcellminer and rcellminerData packages ^80^. rcellminerElasticNet also provides utility functions for summarizing and visualizing elastic net results.

Results for the elastic net analysis are available from this URL: https://discover.nci.nih.gov/cellminerreviewdata/swath_analysis/swathOutput_062316_all.tar.gz. This compressed file contains results for the analysis run with all features and selected common features. Each drug compound has three files for each combination of molecular features used in a particular run of the elastic net algorithm: 1) a knitr report R Markdown (.Rmd) file containing the code that was run, 2) an RData (.Rdata) file containing the results of each elastic net run (see elasticNet() documentation in the rcellminerElasticNet package), 3) the rendered knitr report as a webpage (.html).

Beyond the knitr report containing code, the elastic net pipeline is made reproducible using a Docker image. Docker (www.docker.com) is an emerging platform for conducting reproducible research in the biomedical research community. All necessary software and dependencies to run the described analysis have been embedded in the available Docker container to provide readers an environment that runs on all major operating systems (including Windows, OSX, and Linux), making Docker containers self-contained, portable, and capable of performing at levels similar to the host system.

The Docker container is available at the Docker Hub repository: cannin/swath (https://hub.docker.com/r/cannin/swath/). Key dependencies installed, include: RStudio Server (https://www.rstudio.com/), rcellminer/rcellminerData ^80^, and rcellminerElasticNet. With these installed dependencies, readers have the opportunity to 1) re-run analysis for specific drug compounds and modify the code in order to extend the analysis using RStudio Server, a web-based version of the RStudio R editor, and 2) use an R Shiny app web-based data explorer to further understand described results. Instructions on the usage of the Docker container are located at the rcellminerElasticNet project page (https://bitbucket.org/cbio_mskcc/rcellminerelasticnet).

### Data deposition

The NCI-60 SWATH data sets and SWATH assay library has been deposited in PRIDE. Project Name: NCI60 proteome by PCT-SWATH; Project accession: PXD003539. Reviewer account details:

Username:
Password: dWdyptzf

The protein data matrix has also been deposited in ArrayExpress. Project accession: E-PROT-2. Project title: Proteomic profiling of NCI60 cell lines from Cancer Cell Line Encyclopedia.

Reviewer account details:

Username: Reviewer_E-PROT-2
Password: gdgywGco

The protein data matrix is also accessible in CellMiner website ^13^ and R package rcellminer ^37^.

