## Supplementary material for "Rapid proteotyping reveals cancer biology and drug response determinants in the NCI-60 cells": supple file

### Supplementary Files

### Supplementary Figure 1

A

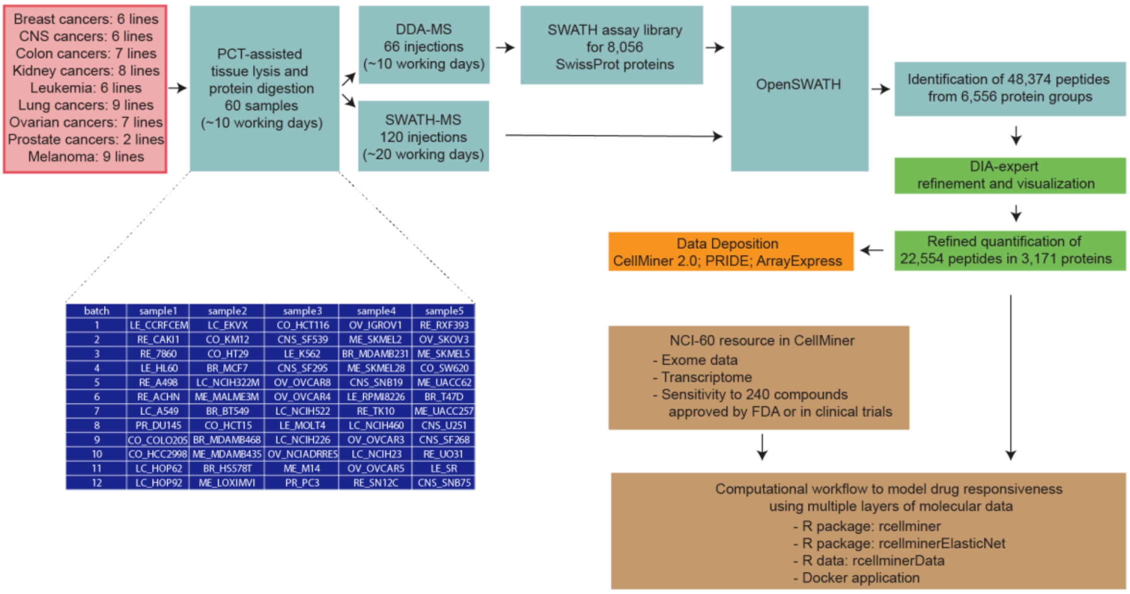


| batch | sample1 | sample2 | sample3 | sample4 | sample5 |
|---|---|---|---|---|---|
| 1 | LE_CCRFCEM | LC_EKVX | CO_HCT116 | OV_IGROV1 | RE_RXF393 |
| 2 | RE_CAKI1 | CO_KM12 | CNS_SF539 | ME_SKMEL2 | OV_SKOV3 |
| 3 | RE_7860 | CO_HT29 | LE_K562 | BR_MDAMB231 | ME_SKMEL5 |
| 4 | LE_HL60 | BR_MCF7 | CNS_SF295 | ME_SKMEL28 | CO_SW620 |
| 5 | RE_A498 | LC_NCIH322M | OV_OVCAR8 | CNS_SNB19 | ME_UACC62 |
| 6 | RE_ACHN | ME_MALME3M | OV_OVCAR4 | LE_RPMI8226 | BR_T47D |
| 7 | LC_A549 | BR_BT549 | LC_NCIH522 | RE_TK10 | ME_UACC257 |
| 8 | PR_DU145 | CO_HCT15 | LE_MOLT4 | LC_NCIH460 | CNS_U251 |
| 9 | CO_COLO205 | BR_MDAMB468 | LC_NCIH226 | OV_OVCAR3 | CNS_SF268 |
| 10 | CO_HCC2998 | ME_MDAMB435 | OV_NCIADRRES | LC_NCIH23 | RE_UO31 |
| 11 | LC_HOP62 | BR_HS578T | ME_M14 | OV_OVCAR5 | LE_SR |
| 12 | LC_HOP92 | ME_LOXIMVI | PR_PC3 | RE_SN12C | CNS_SNB75 |

B

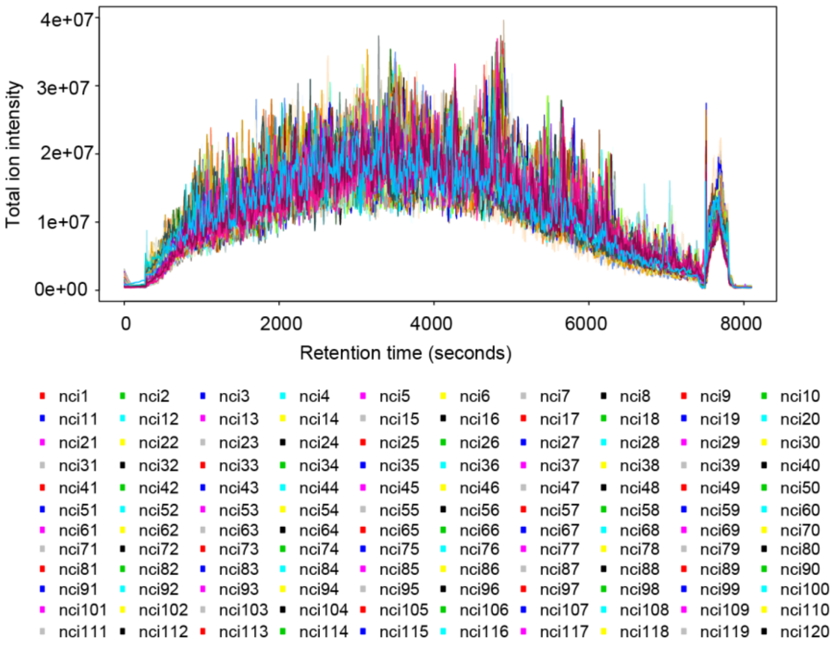


**Supplementary Figure 1. Workflow for generating the NCI-60 proteome maps and predicting phenotypes. (A)** Flowchart of experimental design. The NCI-60 cell pellets were divided into 12 batches, lyzed and digested using the PCT method. The peptides were first analyzed in DDA mode to build a SWATH assay library. In total we performed 66 DDA injections either from whole cell lysate or fractionated samples. Each sample was analyzed in SWATH mode twice. The SWATH data were processed using software tools including OpenSWATH and DIA-expert in sequence. Our data were deposited in several public databases including CellMiner 2.0. Subsequently, we developed a computational workflow to model drug responsiveness using multiple layers of molecular data. The generation of a spectral library specifically for the NCI-60 cells consumed ca. 10 working days. For studies of this type this step is optional because similar results can be obtained from the use of publicly accessible, extensive human spectral libraries such as the pan-human library [1]. (**B**) raw mass spectrometric signal for the 120 SWATH runs. Total ion chromatography graphs are shown. The index of the 120 NCI SWATH files are explained in **Supplementary Table 1**.

**Supplementary Figure 2**

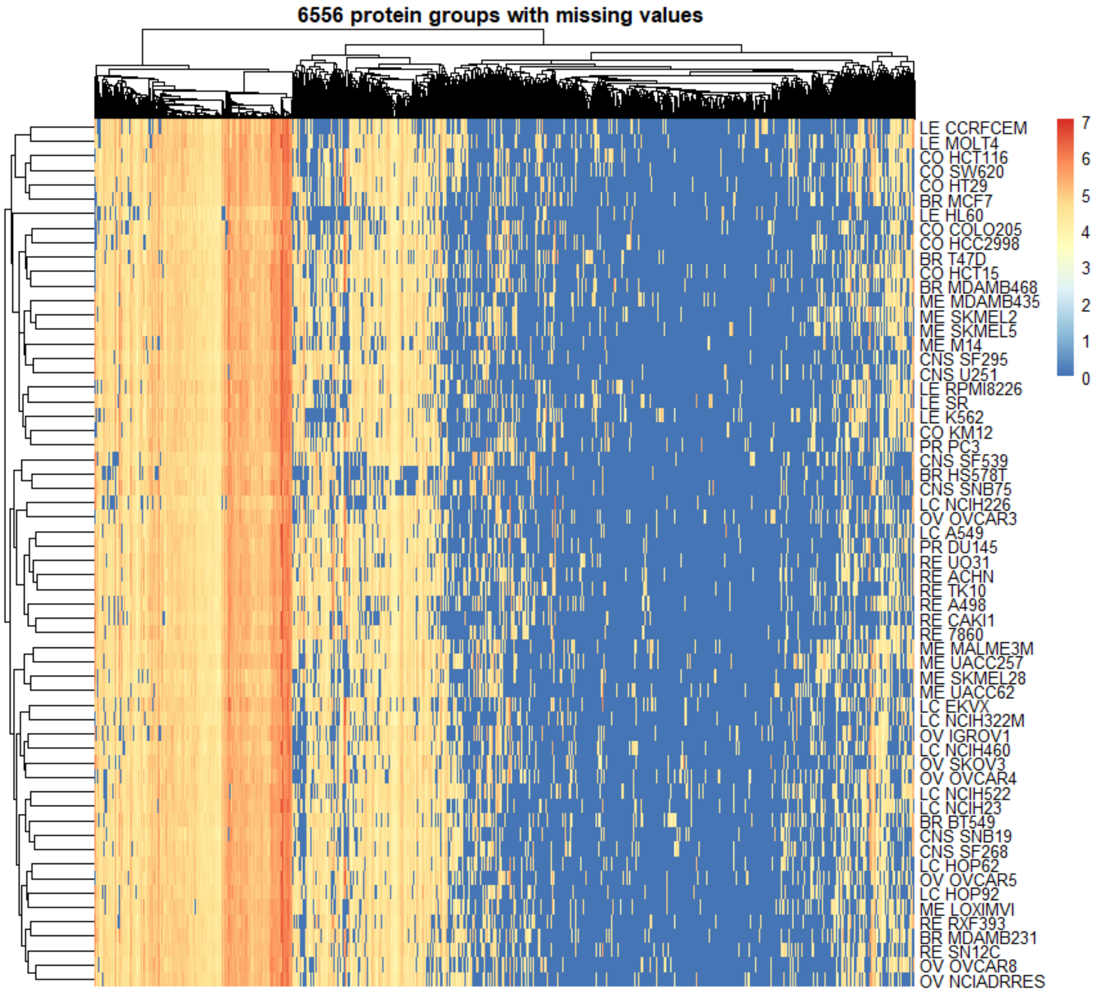


**Supplementary Figure 2. Unsupervised clustering of 6556 protein groups identified and quantified in the NCI-60 cells.** Using the SWATH library containing 8056 protein groups, we displayed the identified and quantified protein groups after unsupervised clustering of both cells and proteins based on their log10 transformed intensity values.

**Supplementary Figure 3**

a

| Min number of peptides | Number of proteins | R2 |
|---|---|---|
| 1 | 3171 | 0.9872 |
| 2 | 2200 | 0.9800 |
| 3 | 1741 | 0.9815 |
| 4 | 1428 | 0.9823 |

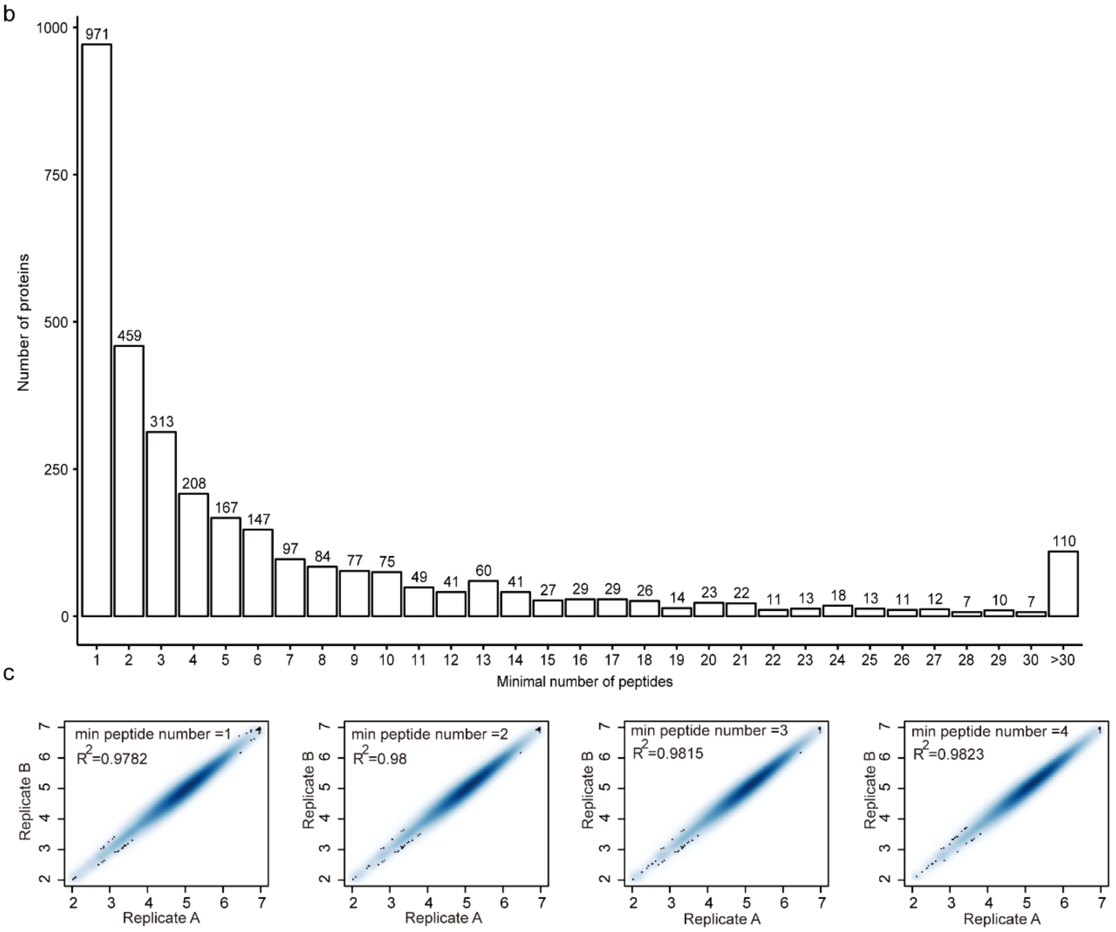


**Supplementary Figure 3. Quantitative accuracy of the NCI-60 proteome as a function of the number of peptides quantified per protein.** (**a**) Number of proteins quantified when minimally 1, 2, 3 and 4 peptides were quantified per protein. The R2 values of technical replicates are computed. (**b**) Distribution of protein numbers based on increasing number of peptides. (**c**) The heatmap scatter plot of proteins quantified in two technical replicates when the minimal peptide number is limited to 1, 2, 3 and 4.

**Supplementary Figure 4**

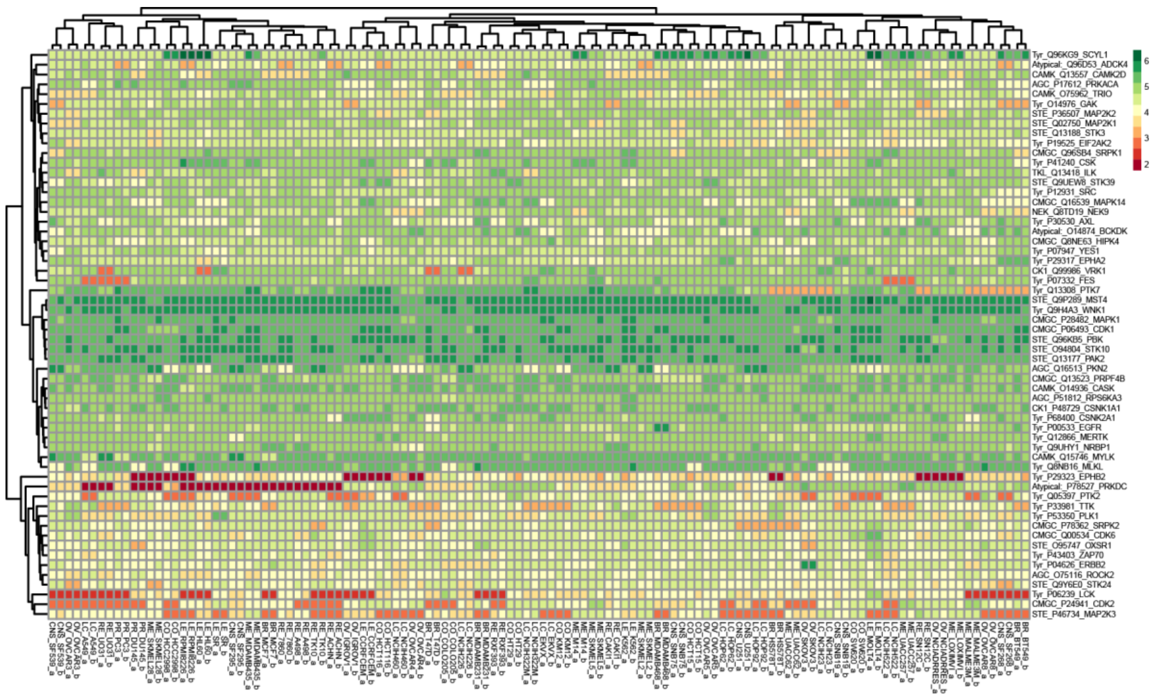


**Supplementary Figure 4. Fifty-eight protein kinases quantified in the NCI-60 cell panel.** Expression of 58 protein kinases, represented by log10 transformed protein intensity values, in the NCI-60. Values are clustered without supervision across both proteins and cell lines.

**Supplementary Figure 5**

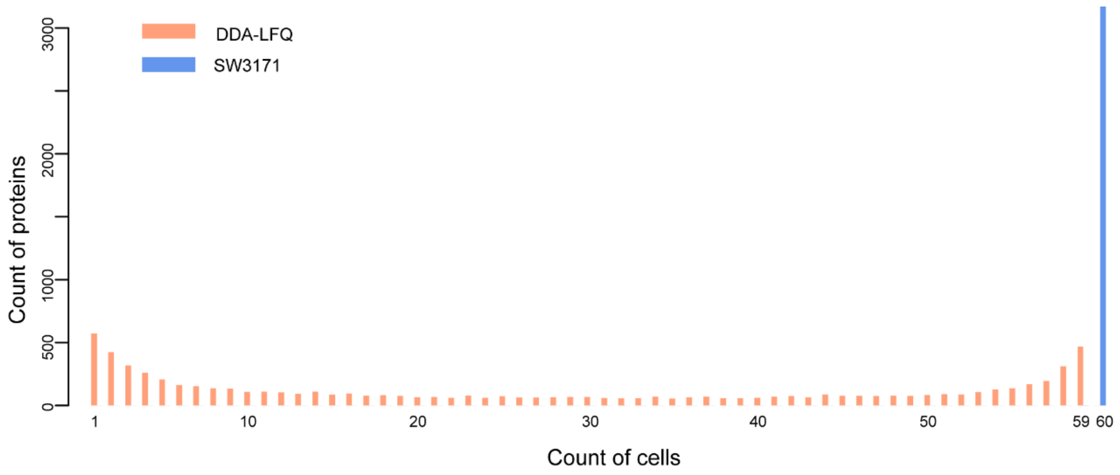


**Supplementary Figure 5. Count of proteins quantified in increasing number of cells.** This plot shows the number of proteins quantified in the NCI-60 cells. DDA-LFQ denotes the LFQ-processed DDA data of the NCI-60 cells. SW3171 means the SWATH data set presented in this study. Most of the SW3171 proteins were quantified in all 60 cells. In DDA-LFQ data set [2], highest numbers of IPI protein groups were quantified in 1 and 59 cells.

### Supplementary Figure 6

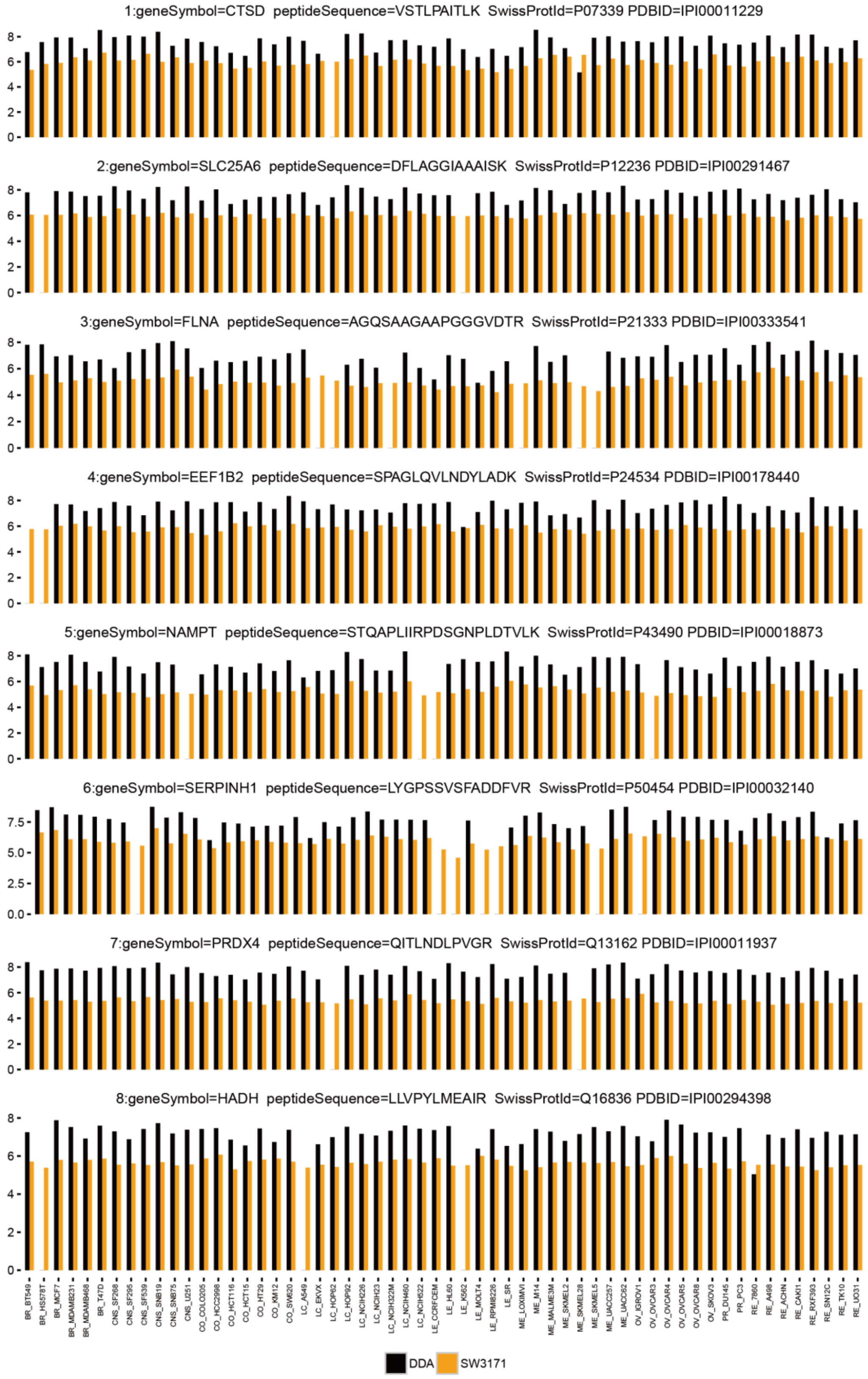

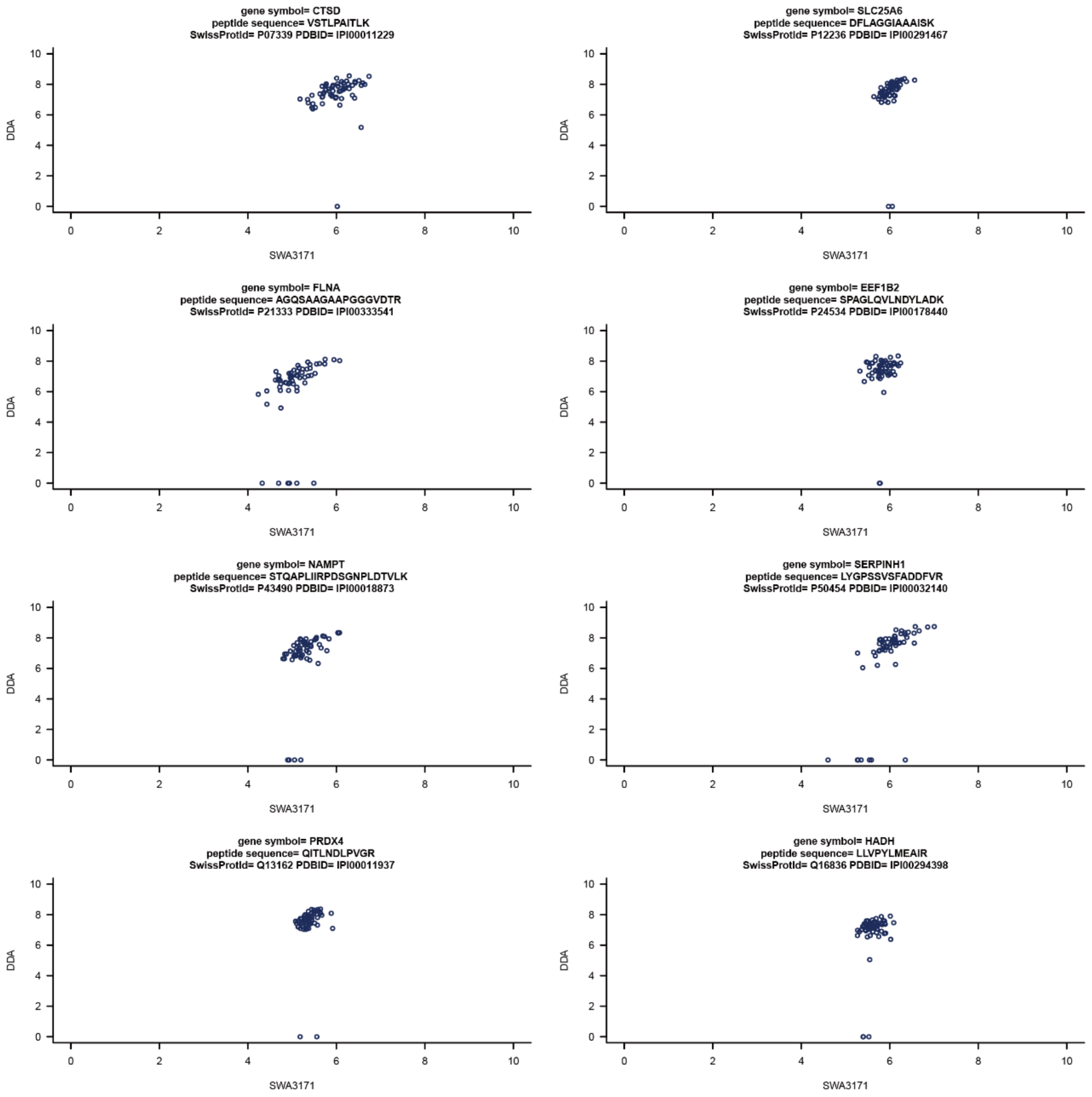


**Supplementary Figure 6. Comparison of 8 representative proteins which have been consistently quantified across nearly all NCI-60 cell lines by both DDA and SWATH data.** The data are shown in both bar plots and scatter plots. Protein intensity was using log10 scaled intensity values.

### Supplementary Figure 7

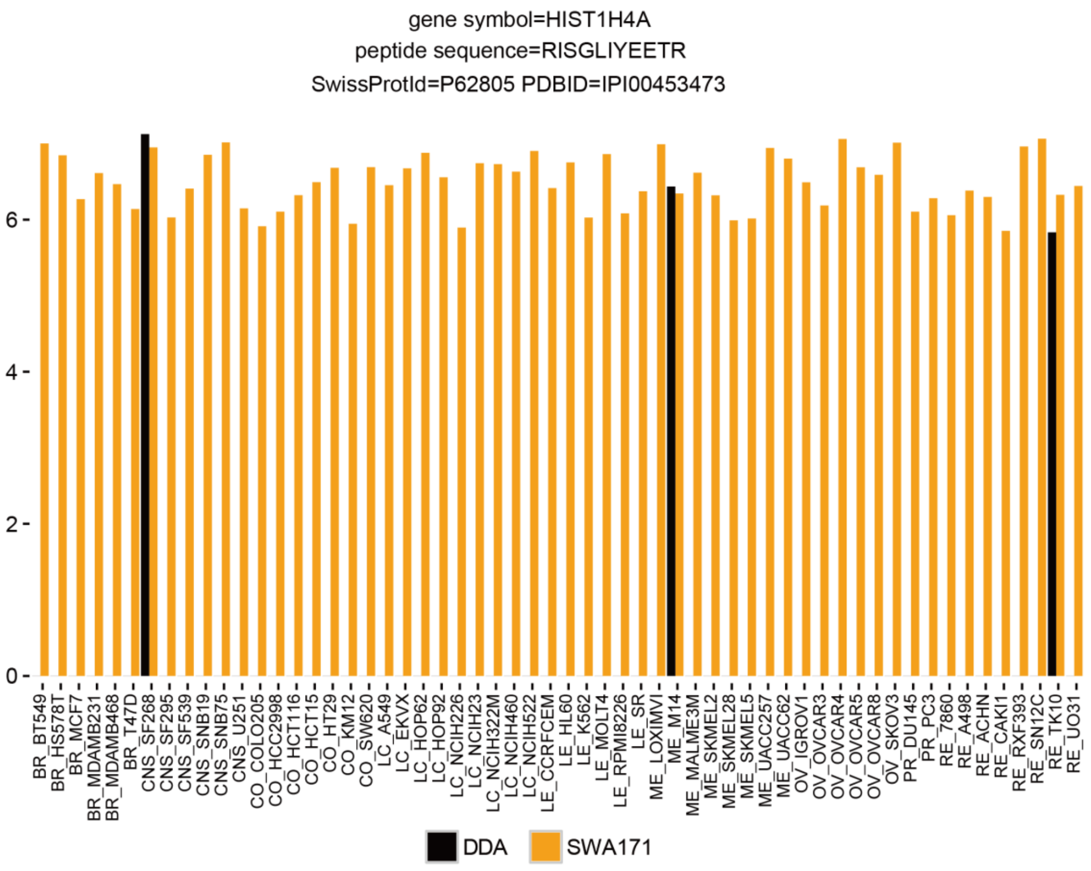

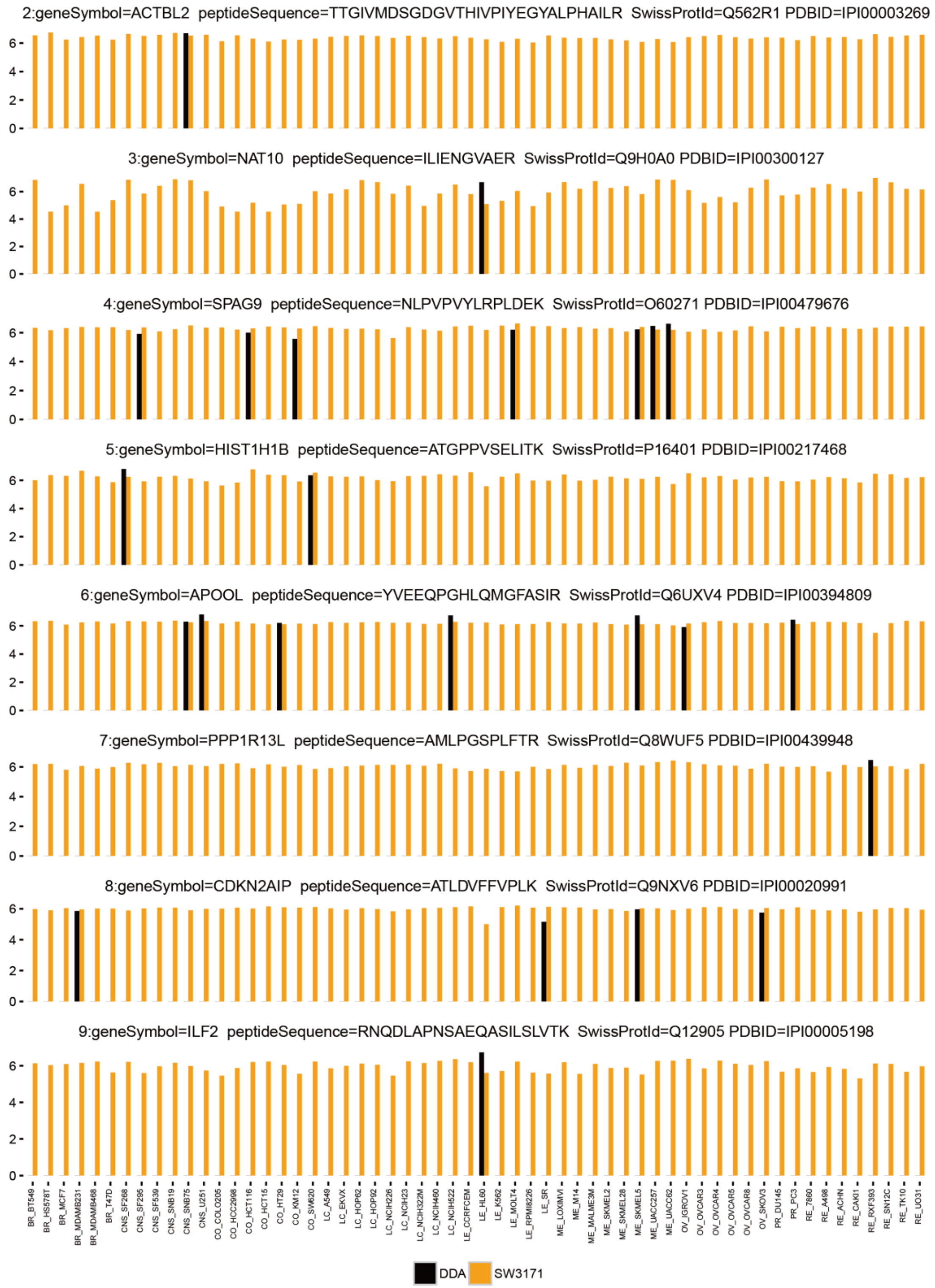
2:geneSymbol=ACTBL2 peptideSequence=TTGIVMDSGDGVTHIVPIYEGYALPHAILR SwissProtId=Q562R1 PDBID=IPI00003269
3:geneSymbol=NAT10 peptideSequence=ILIENGVAER SwissProtId=Q9H0A0 PDBID=IPI00300127
4:geneSymbol=SPAG9 peptideSequence=NLPVPVYLRPLDEK SwissProtId=O60271 PDBID=IPI00479676
5:geneSymbol=HIST1H1B peptideSequence=ATGPPVSELITK SwissProtId=P16401 PDBID=IPI00217468
6:geneSymbol=APOOL peptideSequence=YVEEQPGHLQMGFASIR SwissProtId=Q6UXV4 PDBID=IPI00394809
7:geneSymbol=PPP1R13L peptideSequence=AMLPGSPLFTR SwissProtId=Q8WUF5 PDBID=IPI00439948
8:geneSymbol=CDKN2AIP peptideSequence=ATLDVFFVPLK SwissProtId=Q9NXV6 PDBID=IPI00020991
9:geneSymbol=ILF2 peptideSequence=RNQDLAPNSAEQASILSLVTK SwissProtId=Q12905 PDBID=IPI00005198
0
2
4
6
BR_BT549
BR_HS578T
BR_MCF7
BR_MDAMB231
BR_MDAMB468
BR_T47D
CNS_SF268
CNS_SF295
CNS_SF539
CNS_SNB19
CNS_SNB75
CNS_U251
CO_COLO205
CO_HCC2998
CO_HCT116
CO_HCT15
CO_HT29
CO_KM12
CO_SW620
LC_A549
LC_EKVX
LC_HOP62
LC_HOP92
LC_NCIH226
LC_NCIH23
LC_NCIH322M
LC_NCIH460
LC_NCIH522
LE_CCRFCEM
LE_HL60
LE_K562
LE_MOLT4
LE_RPMI8226
LE_SR
ME_LOXIMVI
ME_M14
ME_MALME3M
ME_SKMEL2
ME_SKMEL28
ME_SKMEL5
ME_UACC257
ME_UACC62
OV_IGROV1
OV_OVCAR3
OV_OVCAR4
OV_OVCAR5
OV_OVCAR8
OV_SKOV3
PR_DU145
PR_PC3
RE_7860
RE_A498
RE_ACHN
RE_CAKI1
RE_RXF393
RE_SN12C
RE_TK10
RE_UO31
DDA
SW3171

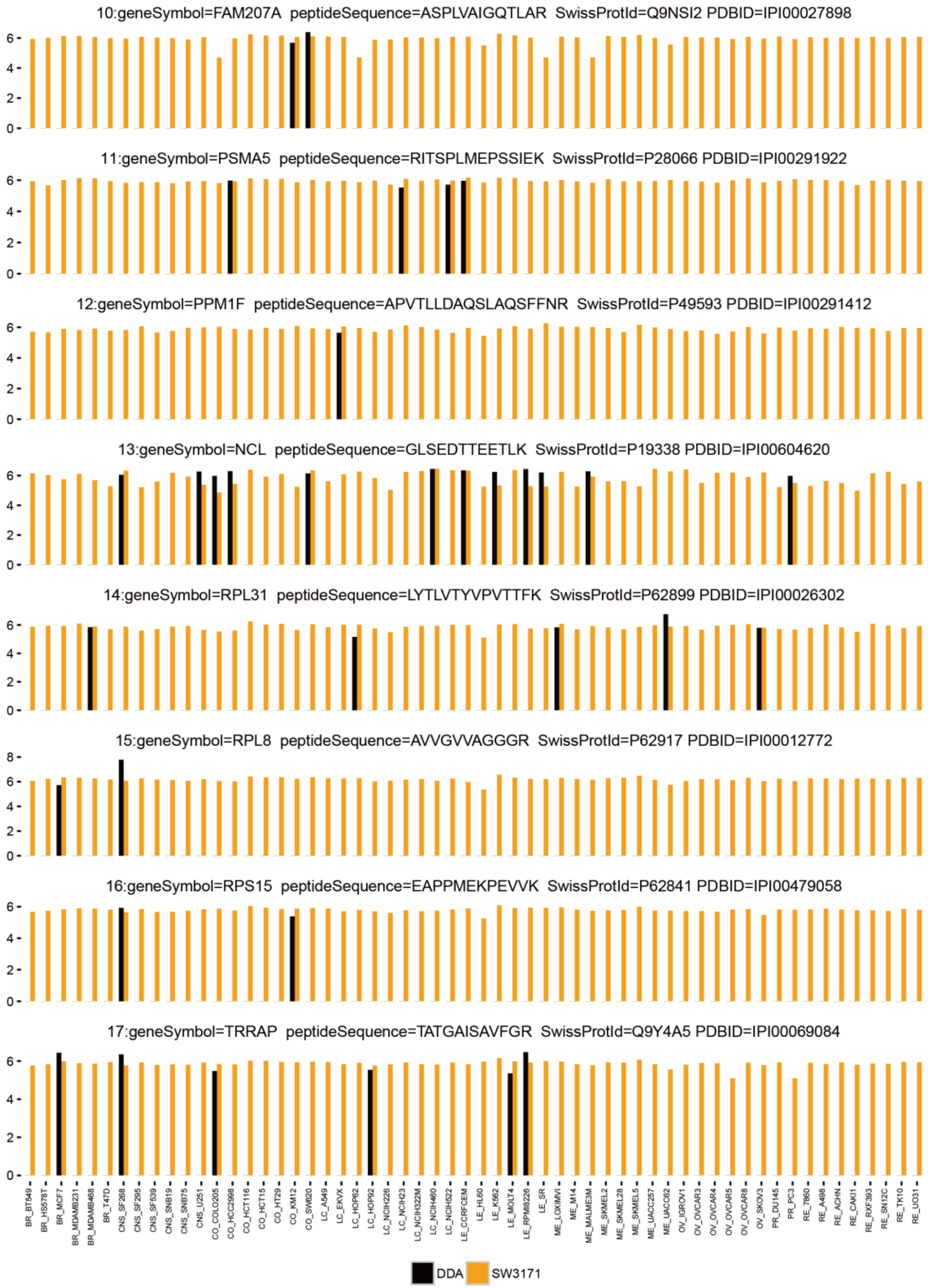

10:geneSymbol=FAM207A peptideSequence=ASPLVAIGQTLAR SwissProtId=Q9NSI2 PDBID=IPI00027898
11:geneSymbol=PSMA5 peptideSequence=RITSPLMEPSSIEK SwissProtId=P28066 PDBID=IPI00291922
12:geneSymbol=PPM1F peptideSequence=APVTLLDAQSLAQSFFNR SwissProtId=P49593 PDBID=IPI00291412
13:geneSymbol=NCL peptideSequence=GLSEDTTEETLK SwissProtId=P19338 PDBID=IPI00604620
14:geneSymbol=RPL31 peptideSequence=LYTLVTYVPVTTFK SwissProtId=P62899 PDBID=IPI00026302
15:geneSymbol=RPL8 peptideSequence=AVVGVVAGGGR SwissProtId=P62917 PDBID=IPI00012772
16:geneSymbol=RPS15 peptideSequence=EAPPMEKPEVVK SwissProtId=P62841 PDBID=IPI00479058
17:geneSymbol=TRRAP peptideSequence=TATGAISAVFGR SwissProtId=Q9Y4A5 PDBID=IPI00069084
BR_BT549 BR_HS578T BR_MCF7 BR_MDAMB231 BR_MDAMB468 BR_T47D CNS_SF268 CNS_SF295 CNS_SF539 CNS_SNB19 CNS_SNB75 CNS_U251 CO_COLO205 CO_HCC2998 CO_HCT116 CO_HCT15 CO_HT29 CO_KM12 CO_SW620 LC_A549 LC_EKVX LC_HOP62 LC_HOP92 LC_NCIH226 LC_NCIH23 LC_NCIH322M LC_NCIH460 LC_NCIH522 LE_CCRFCEM LE_HL60 LE_K562 LE_MOLT4 LE_RPMI8226 LE_SR ME_LOXIMVI ME_M14 ME_MALME3M ME_SKMEL2 ME_SKMEL28 ME_SKMEL5 ME_UACC257 ME_UACC62 OV_IGROV1 OV_OVCAR3 OV_OVCAR4 OV_OVCAR5 OV_OVCAR8 OV_SKOV3 PR_DU145 PR_PC3 RE_7860 RE_A498 RE_ACHN RE_CAKI1 RE_RXF393 RE_SN12C RE_TK10 RE_UO31
DDA
SW3171

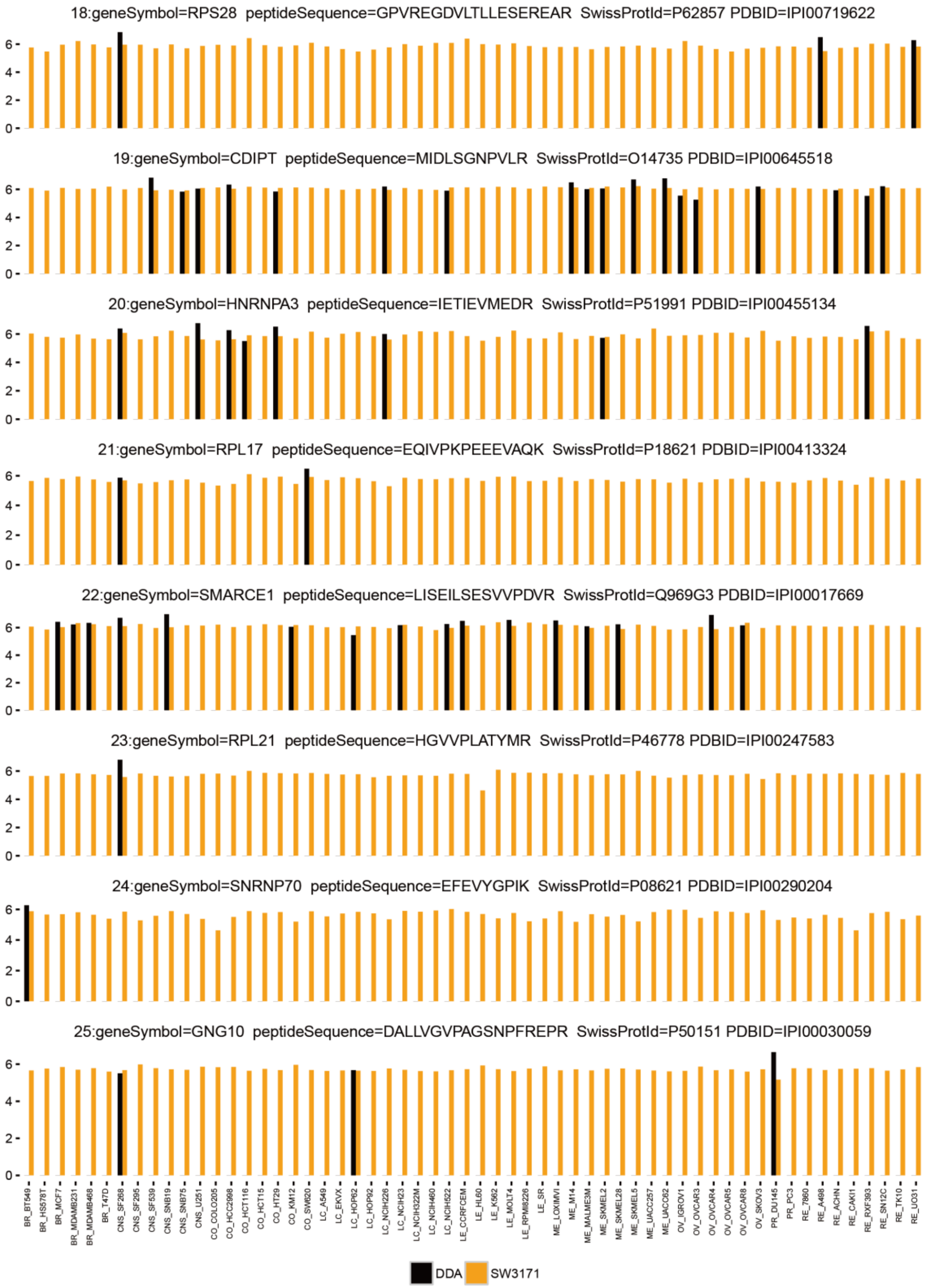
18:geneSymbol=RPS28 peptideSequence=GPVREGDVLTLLESEREAR SwissProtId=P62857 PDBID=IPI00719622
19:geneSymbol=CDIPT peptideSequence=MIDLSGNPVLR SwissProtId=O14735 PDBID=IPI00645518
20:geneSymbol=HNRNPA3 peptideSequence=IETIEVMEDR SwissProtId=P51991 PDBID=IPI00455134
21:geneSymbol=RPL17 peptideSequence=EQIVPKPEEEVAQK SwissProtId=P18621 PDBID=IPI00413324
22:geneSymbol=SMARCE1 peptideSequence=LISEILSESVVPDVR SwissProtId=Q969G3 PDBID=IPI00017669
23:geneSymbol=RPL21 peptideSequence=HGVVPLATYMR SwissProtId=P46778 PDBID=IPI00247583
24:geneSymbol=SNRNP70 peptideSequence=EFEVYGPIK SwissProtId=P08621 PDBID=IPI00290204
25:geneSymbol=GNG10 peptideSequence=DALLVGVPAGSNPFREPR SwissProtId=P50151 PDBID=IPI00030059
DDA
SW3171

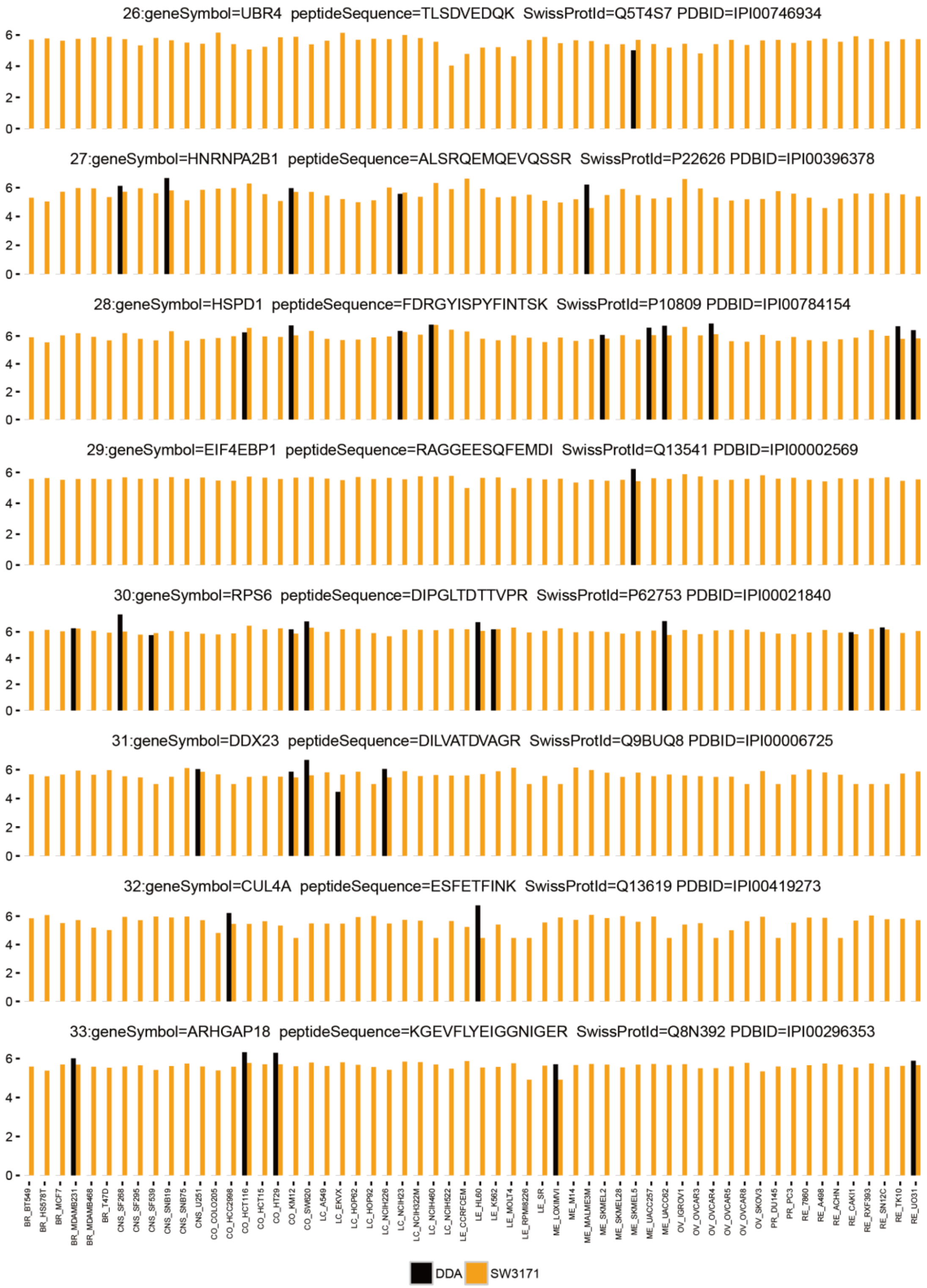
26:geneSymbol=UBR4 peptideSequence=TLSDVEDQK SwissProtId=Q5T4S7 PDBID=IPI00746934
27:geneSymbol=HNRNPA2B1 peptideSequence=ALSRQEMQEVQSSR SwissProtId=P22626 PDBID=IPI00396378
28:geneSymbol=HSPD1 peptideSequence=FDRGYISPYFINTSK SwissProtId=P10809 PDBID=IPI00784154
29:geneSymbol=EIF4EBP1 peptideSequence=RAGGEESQFEMDI SwissProtId=Q13541 PDBID=IPI00002569
30:geneSymbol=RPS6 peptideSequence=DIPGLTDTTVPR SwissProtId=P62753 PDBID=IPI00021840
31:geneSymbol=DDX23 peptideSequence=DILVATDVAGR SwissProtId=Q9BUQ8 PDBID=IPI00006725
32:geneSymbol=CUL4A peptideSequence=ESFETFINK SwissProtId=Q13619 PDBID=IPI00419273
33:geneSymbol=ARHGAP18 peptideSequence=KGEVFLYEIGGNIGER SwissProtId=Q8N392 PDBID=IPI00296353
DDA
SW3171

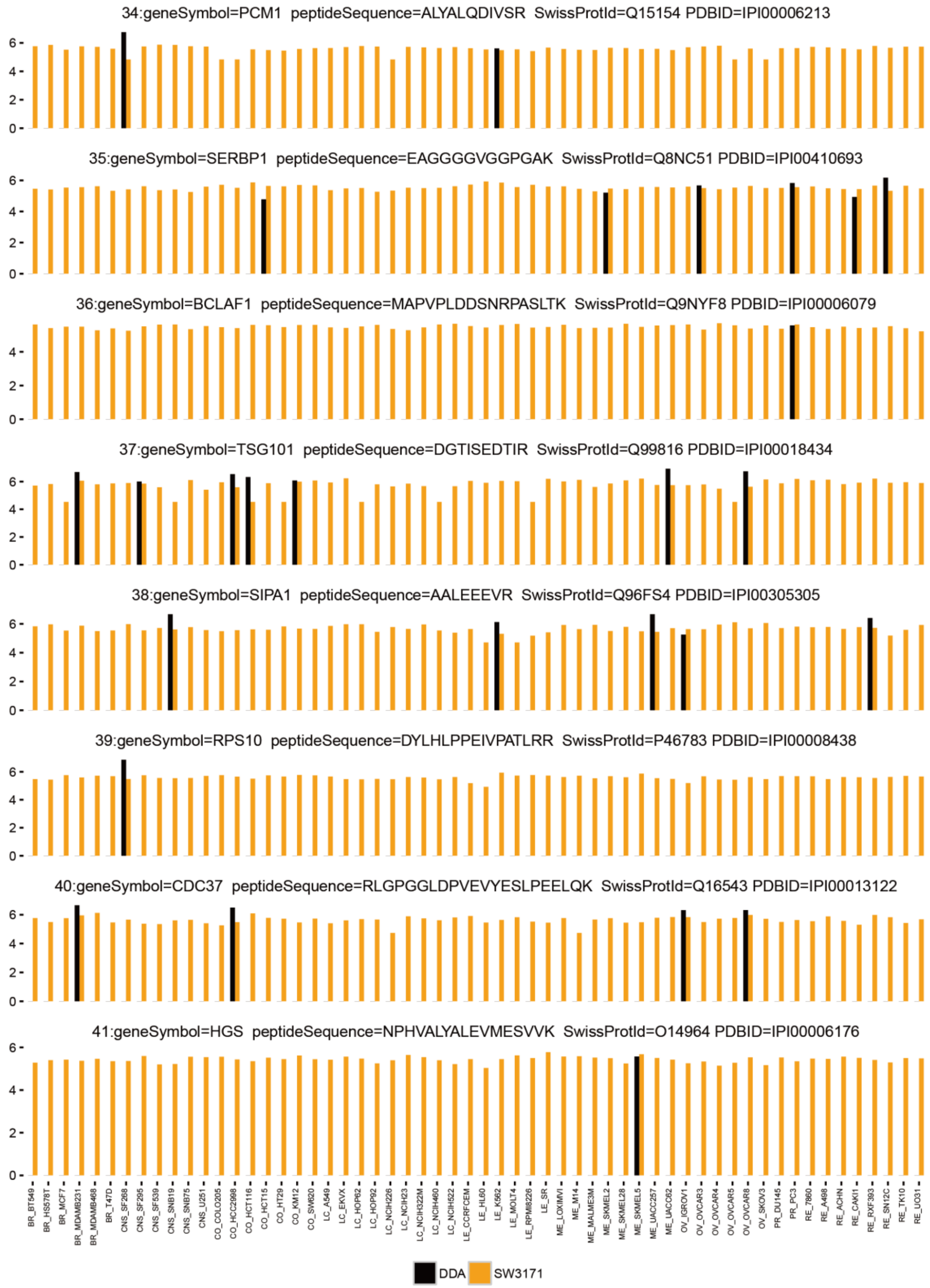
34:geneSymbol=PCM1 peptideSequence=ALYALQDIVSR SwissProtId=Q15154 PDBID=IPI00006213
35:geneSymbol=SERBP1 peptideSequence=EAGGGGVGGPGAK SwissProtId=Q8NC51 PDBID=IPI00410693
36:geneSymbol=BCLAF1 peptideSequence=MAPVPLDDSNRPASLTK SwissProtId=Q9NYF8 PDBID=IPI00006079
37:geneSymbol=TSG101 peptideSequence=DGTISEDTIR SwissProtId=Q99816 PDBID=IPI00018434
38:geneSymbol=SIPA1 peptideSequence=AALEEEVR SwissProtId=Q96FS4 PDBID=IPI00305305
39:geneSymbol=RPS10 peptideSequence=DYLHLPPEIVPATLRR SwissProtId=P46783 PDBID=IPI00008438
40:geneSymbol=CDC37 peptideSequence=RLGPGGLDPVEVYESLPEELQK SwissProtId=Q16543 PDBID=IPI00013122
41:geneSymbol=HGS peptideSequence=NPHVALYALEVMESVVK SwissProtId=O14964 PDBID=IPI00006176
DDA
SW3171

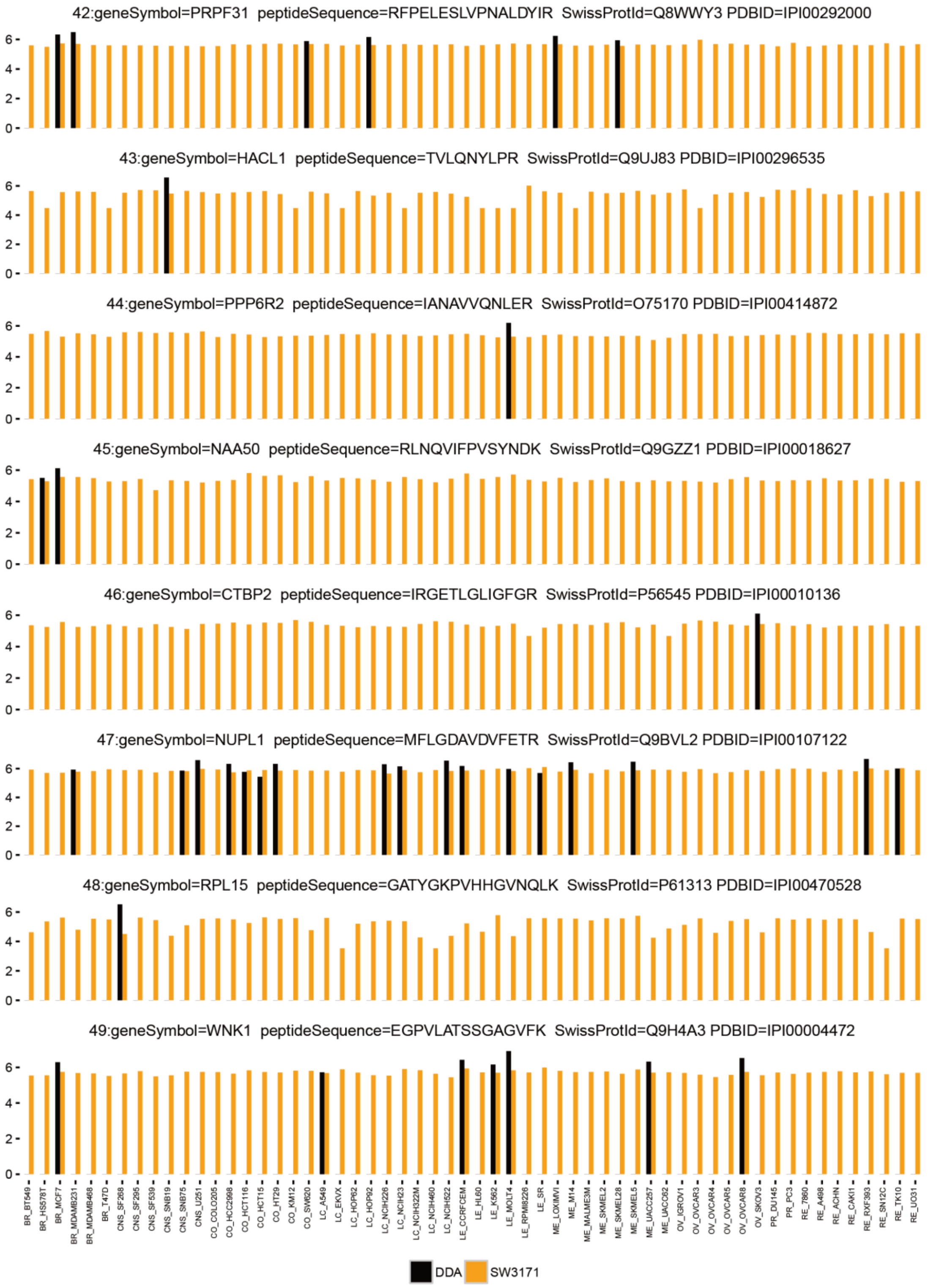
42:geneSymbol=PRPF31 peptideSequence=RFPELESLVPNALDYIR SwissProtId=Q8WWY3 PDBID=IPI00292000
43:geneSymbol=HACL1 peptideSequence=TVLQNYLPR SwissProtId=Q9UJ83 PDBID=IPI00296535
44:geneSymbol=PPP6R2 peptideSequence=IANAVVQNLER SwissProtId=O75170 PDBID=IPI00414872
45:geneSymbol=NAA50 peptideSequence=RLNQVIFPVSYNDK SwissProtId=Q9GZZ1 PDBID=IPI00018627
46:geneSymbol=CTBP2 peptideSequence=IRGETLGLIGFGR SwissProtId=P56545 PDBID=IPI00010136
47:geneSymbol=NUPL1 peptideSequence=MFLGDAVDVFETR SwissProtId=Q9BVL2 PDBID=IPI00107122
48:geneSymbol=RPL15 peptideSequence=GATYGKPVHHGVNQLK SwissProtId=P61313 PDBID=IPI00470528
49:geneSymbol=WNK1 peptideSequence=EGPVLATSSGAGVFK SwissProtId=Q9H4A3 PDBID=IPI00004472
DDA
SW3171

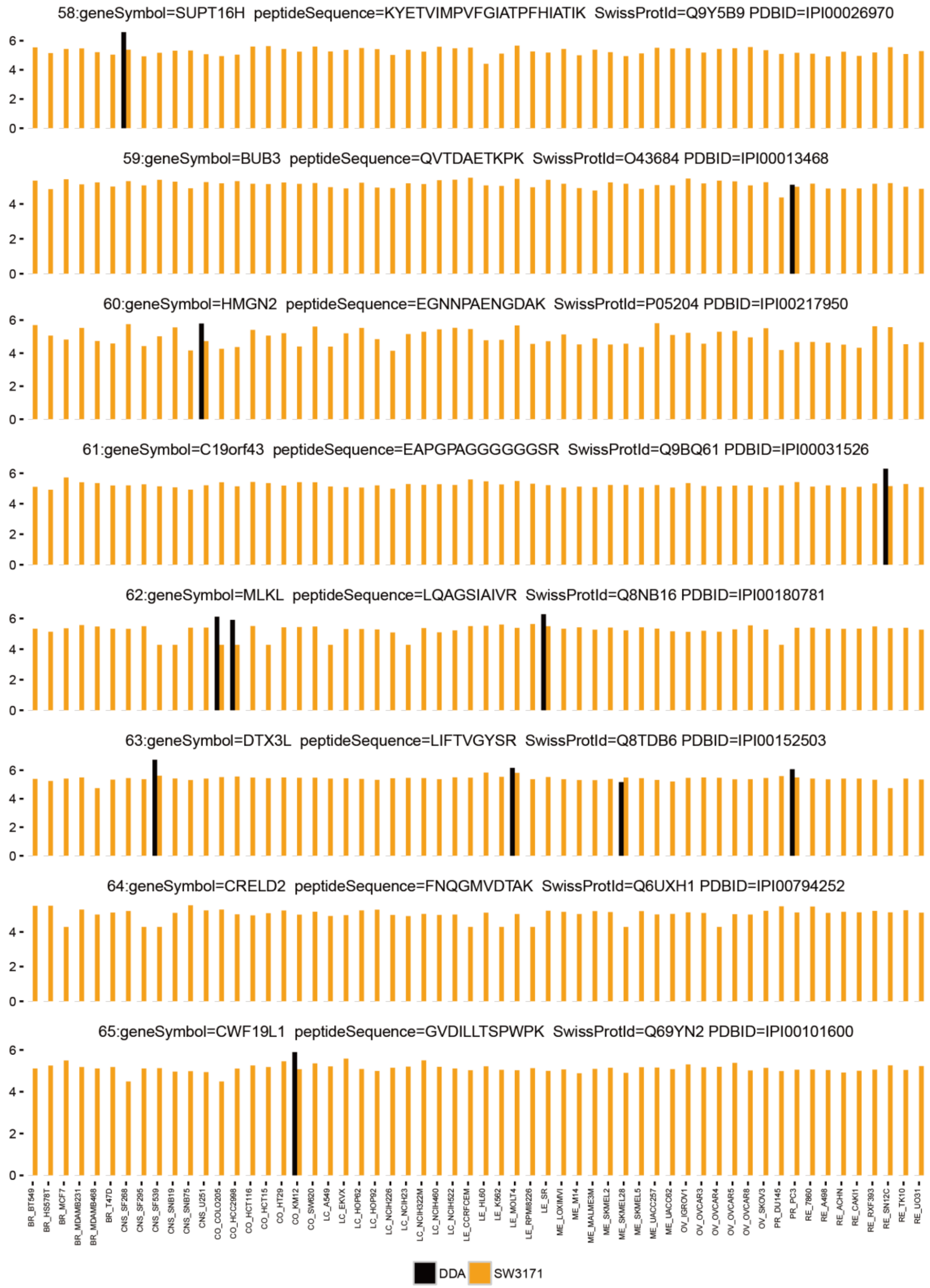
58:geneSymbol=SUPT16H peptideSequence=KYETVIMPVFGIATPFHIATIK SwissProtId=Q9Y5B9 PDBID=IPI00026970
59:geneSymbol=BUB3 peptideSequence=QVTDAETKPK SwissProtId=O43684 PDBID=IPI00013468
60:geneSymbol=HMGN2 peptideSequence=EGNNPAENGDAK SwissProtId=P05204 PDBID=IPI00217950
61:geneSymbol=C19orf43 peptideSequence=EAPGPAGGGGGGSR SwissProtId=Q9BQ61 PDBID=IPI00031526
62:geneSymbol=MLKL peptideSequence=LQAGSIAIVR SwissProtId=Q8NB16 PDBID=IPI00180781
63:geneSymbol=DTX3L peptideSequence=LIFTVGYSR SwissProtId=Q8TDB6 PDBID=IPI00152503
64:geneSymbol=CRELD2 peptideSequence=FNQGMVDTAK SwissProtId=Q6UXH1 PDBID=IPI00794252
65:geneSymbol=CWF19L1 peptideSequence=GVDILLTSPWPK SwissProtId=Q69YN2 PDBID=IPI00101600
DDA
SW3171

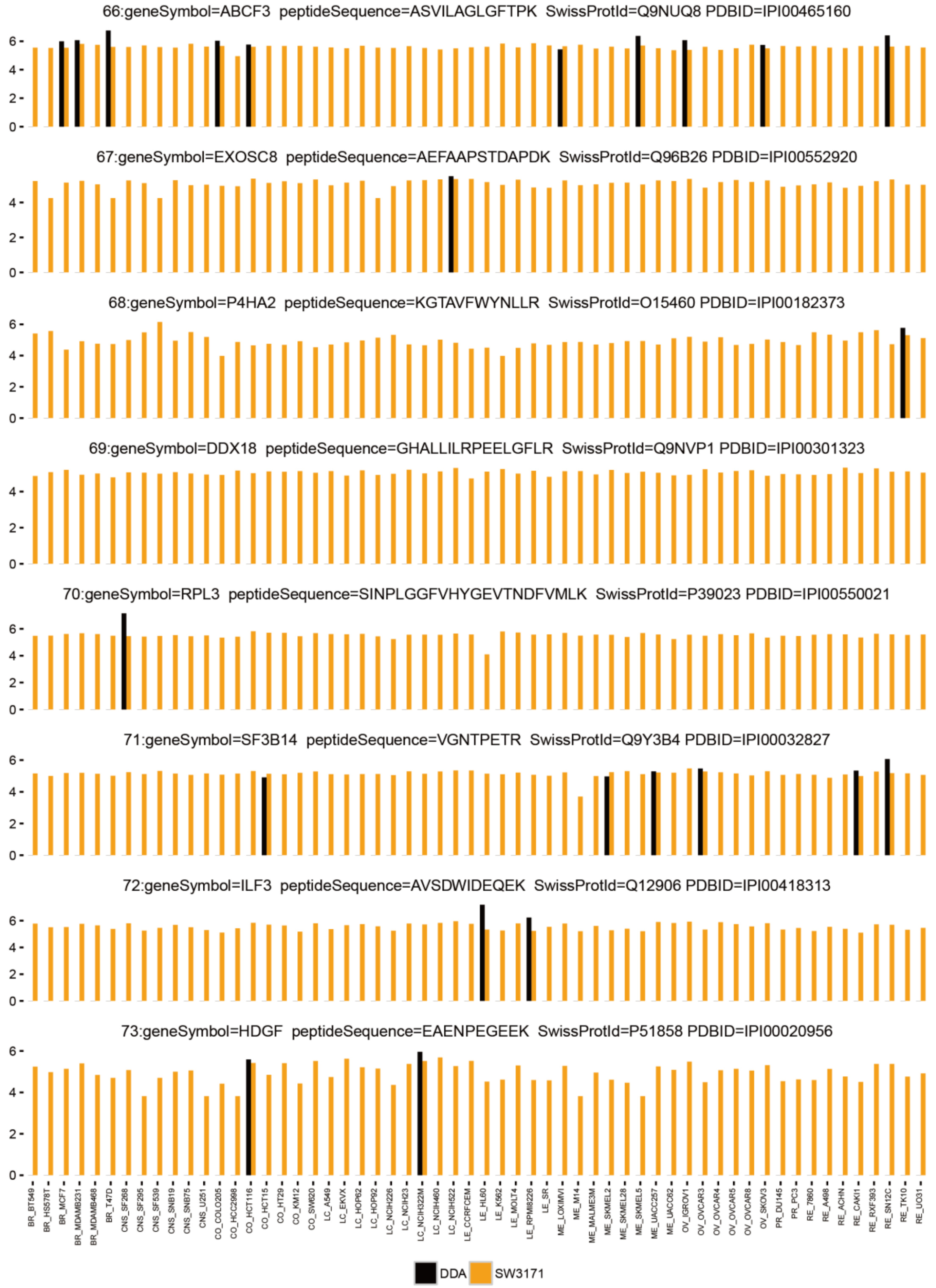
66:geneSymbol=ABCF3 peptideSequence=ASVILAGLGFTPK SwissProtId=Q9NUQ8 PDBID=IPI00465160
67:geneSymbol=EXOSC8 peptideSequence=AEFAAPSTDAPDK SwissProtId=Q96B26 PDBID=IPI00552920
68:geneSymbol=P4HA2 peptideSequence=KGTAVFWYNLLR SwissProtId=O15460 PDBID=IPI00182373
69:geneSymbol=DDX18 peptideSequence=GHALLILRPEELGFLR SwissProtId=Q9NVP1 PDBID=IPI00301323
70:geneSymbol=RPL3 peptideSequence=SINPLGGFVHYGEVTNDFVMLK SwissProtId=P39023 PDBID=IPI00550021
71:geneSymbol=SF3B14 peptideSequence=VGNTPETR SwissProtId=Q9Y3B4 PDBID=IPI00032827
72:geneSymbol=ILF3 peptideSequence=AVSDWIDEQEK SwissProtId=Q12906 PDBID=IPI00418313
73:geneSymbol=HDGF peptideSequence=EAENPEGEEK SwissProtId=P51858 PDBID=IPI00020956
BR_BT549 BR_HS578T BR_MCF7 BR_MDAMB231 BR_MDAMB468 BR_T47D CNS_SF268 CNS_SF295 CNS_SF539 CNS_SNB19 CNS_SNB75 CNS_U251 CO_COLO205 CO_HCC2998 CO_HCT116 CO_HCT15 CO_HT29 CO_KM12 CO_SW620 LC_A549 LC_EKVX LC_HOP62 LC_HOP92 LC_NCIH226 LC_NCIH23 LC_NCIH322M LC_NCIH460 LC_NCIH522 LE_CCRFCEM LE_HL60 LE_K562 LE_MOLT4 LE_RPMI8226 LE_SR ME_LOXIMVI ME_M14 ME_MALME3M ME_SKMEL2 ME_SKMEL28 ME_SKMEL5 ME_UACC257 ME_UACC62 OV_IGROV1 OV_OVCAR3 OV_OVCAR4 OV_OVCAR5 OV_OVCAR8 OV_SKOV3 PR_DU145 PR_PC3 RE_7860 RE_A498 RE_ACHN RE_CAKI1 RE_RXF393 RE_SN12C RE_TK10 RE_UO31
DDA
SW3171

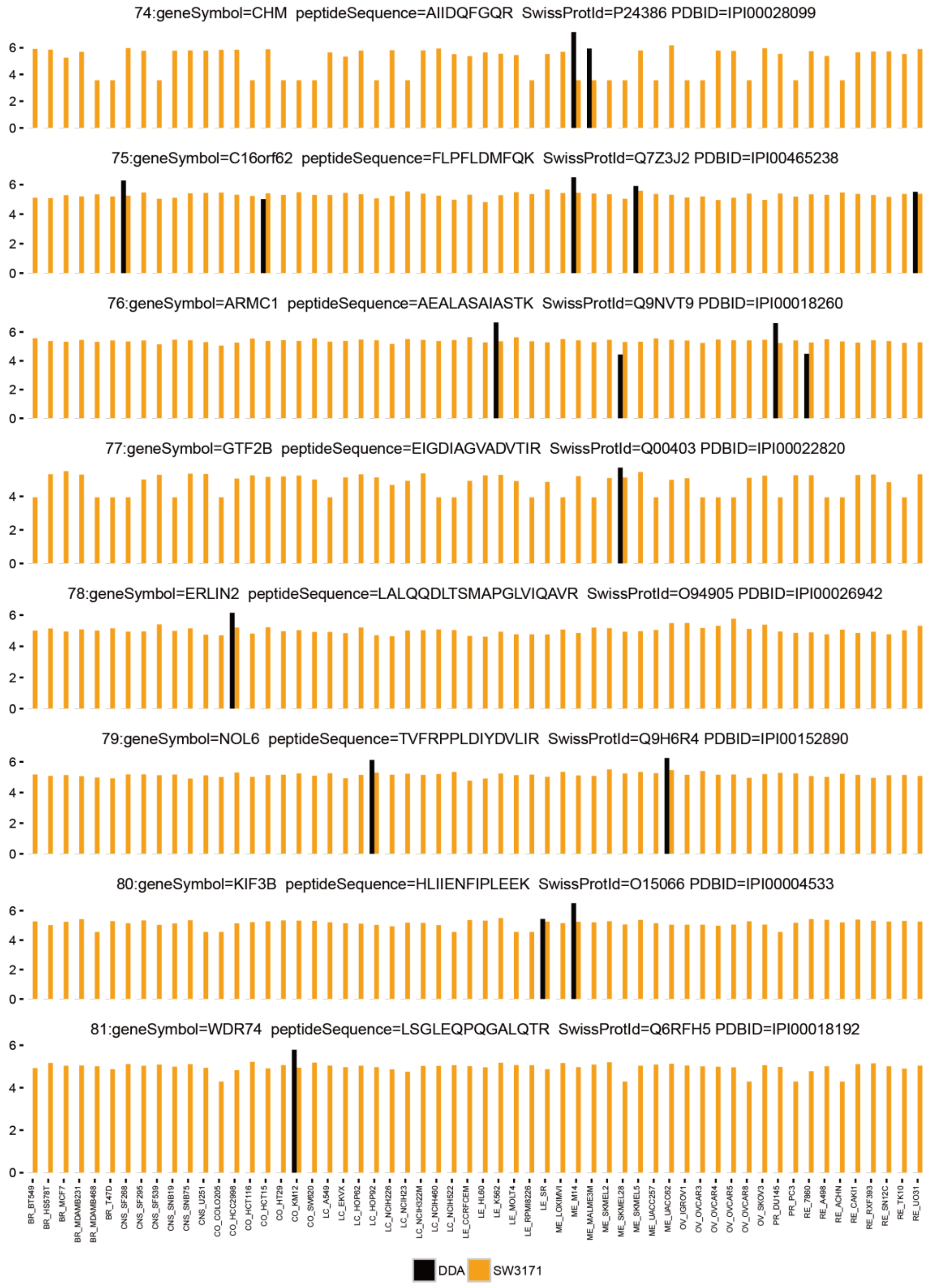
74:geneSymbol=CHM peptideSequence=AIIDQFGQR SwissProtId=P24386 PDBID=IPI00028099
75:geneSymbol=C16orf62 peptideSequence=FLPFLDMFQK SwissProtId=Q7Z3J2 PDBID=IPI00465238
76:geneSymbol=ARMC1 peptideSequence=AEALASAIASTK SwissProtId=Q9NVT9 PDBID=IPI00018260
77:geneSymbol=GTF2B peptideSequence=EIGDIAGVADVTIR SwissProtId=Q00403 PDBID=IPI00022820
78:geneSymbol=ERLIN2 peptideSequence=LALQQDLTSMAPGLVIQAVR SwissProtId=O94905 PDBID=IPI00026942
79:geneSymbol=NOL6 peptideSequence=TVFRPPLDIYDVLIR SwissProtId=Q9H6R4 PDBID=IPI00152890
80:geneSymbol=KIF3B peptideSequence=HLIIENFIPLEEK SwissProtId=O15066 PDBID=IPI00004533
81:geneSymbol=WDR74 peptideSequence=LSGLEQPQGALQTR SwissProtId=Q6RFH5 PDBID=IPI00018192
DDA
SW3171

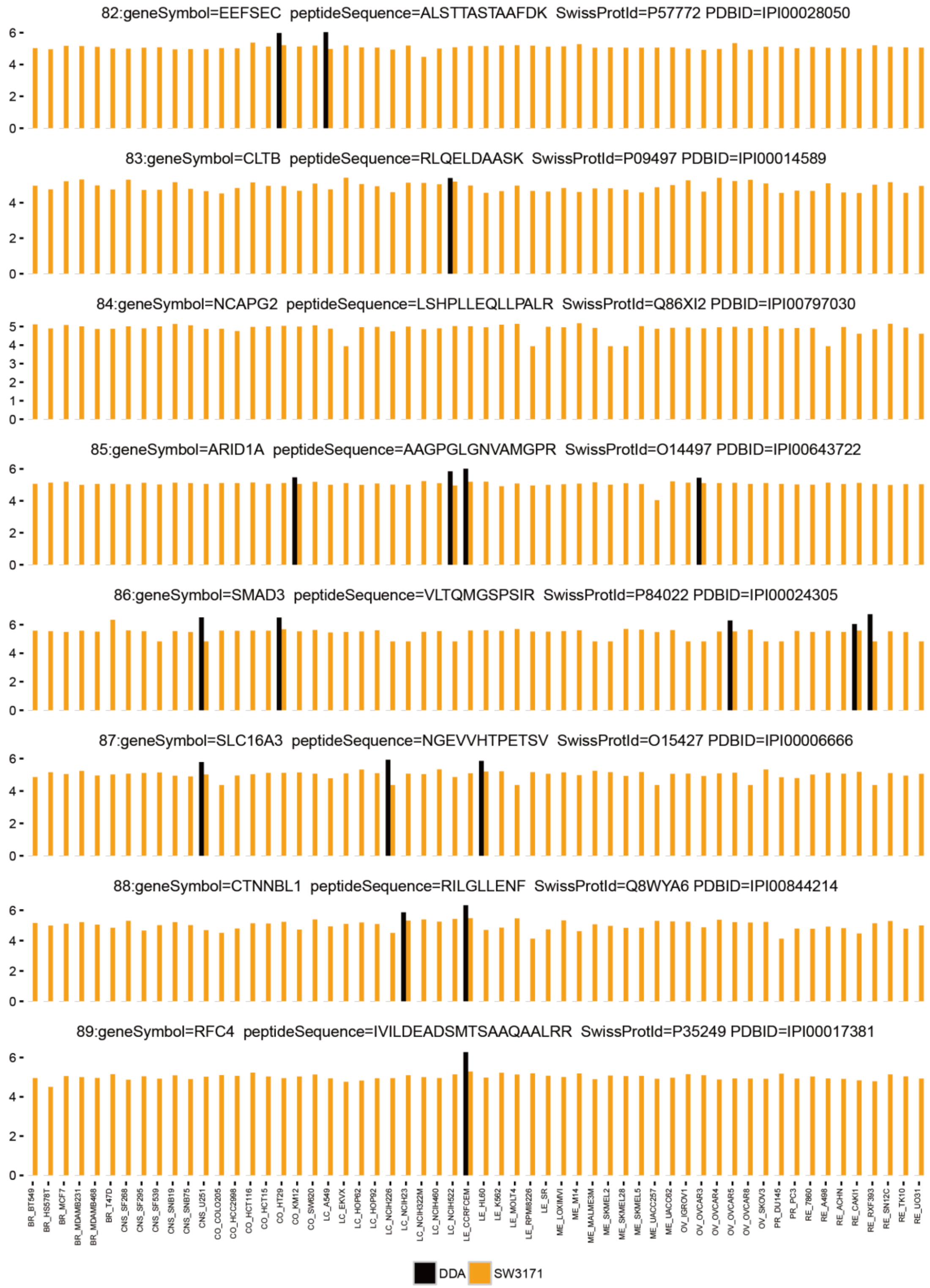
82:geneSymbol=EEFSEC peptideSequence=ALSTTASTAAFDK SwissProtId=P57772 PDBID=IPI00028050
83:geneSymbol=CLTB peptideSequence=RLQELDAASK SwissProtId=P09497 PDBID=IPI00014589
84:geneSymbol=NCAPG2 peptideSequence=LSHPLLEQLLPALR SwissProtId=Q86XI2 PDBID=IPI00797030
85:geneSymbol=ARID1A peptideSequence=AAGPGLGNVAMGPR SwissProtId=O14497 PDBID=IPI00643722
86:geneSymbol=SMAD3 peptideSequence=VLTQMGSPSIR SwissProtId=P84022 PDBID=IPI00024305
87:geneSymbol=SLC16A3 peptideSequence=NGEVVHTPETSV SwissProtId=O15427 PDBID=IPI00006666
88:geneSymbol=CTNNBL1 peptideSequence=RILGLLENF SwissProtId=Q8WYA6 PDBID=IPI00844214
89:geneSymbol=RFC4 peptideSequence=IVILDEADSMTSAAQAALRR SwissProtId=P35249 PDBID=IPI00017381
BR_BT549
BR_HS578T
BR_MCF7
BR_MDAMB231
BR_MDAMB468
BR_T47D
CNS_SF268
CNS_SF295
CNS_SF539
CNS_SNB19
CNS_SNB75
CNS_U251
CO_COLO205
CO_HCC2998
CO_HCT116
CO_HCT15
CO_HT29
CO_KM12
CO_SW620
LC_A549
LC_EKVX
LC_HOP62
LC_HOP92
LC_NCIH226
LC_NCIH23
LC_NCIH322M
LC_NCIH460
LC_NCIH522
LE_CCRFCEM
LE_HL60
LE_K562
LE_MOLT4
LE_RPMI8226
LE_SR
ME_LOXIMVI
ME_M14
ME_MALME3M
ME_SKMEL2
ME_SKMEL28
ME_SKMEL5
ME_UACC257
ME_UACC62
OV_IGROV1
OV_OVCAR3
OV_OVCAR4
OV_OVCAR5
OV_OVCAR8
OV_SKOV3
PR_DU145
PR_PC3
RE_7860
RE_A498
RE_ACHN
RE_CAKI1
RE_RXF393
RE_SN12C
RE_TK10
RE_UO31
DDA
SW3171

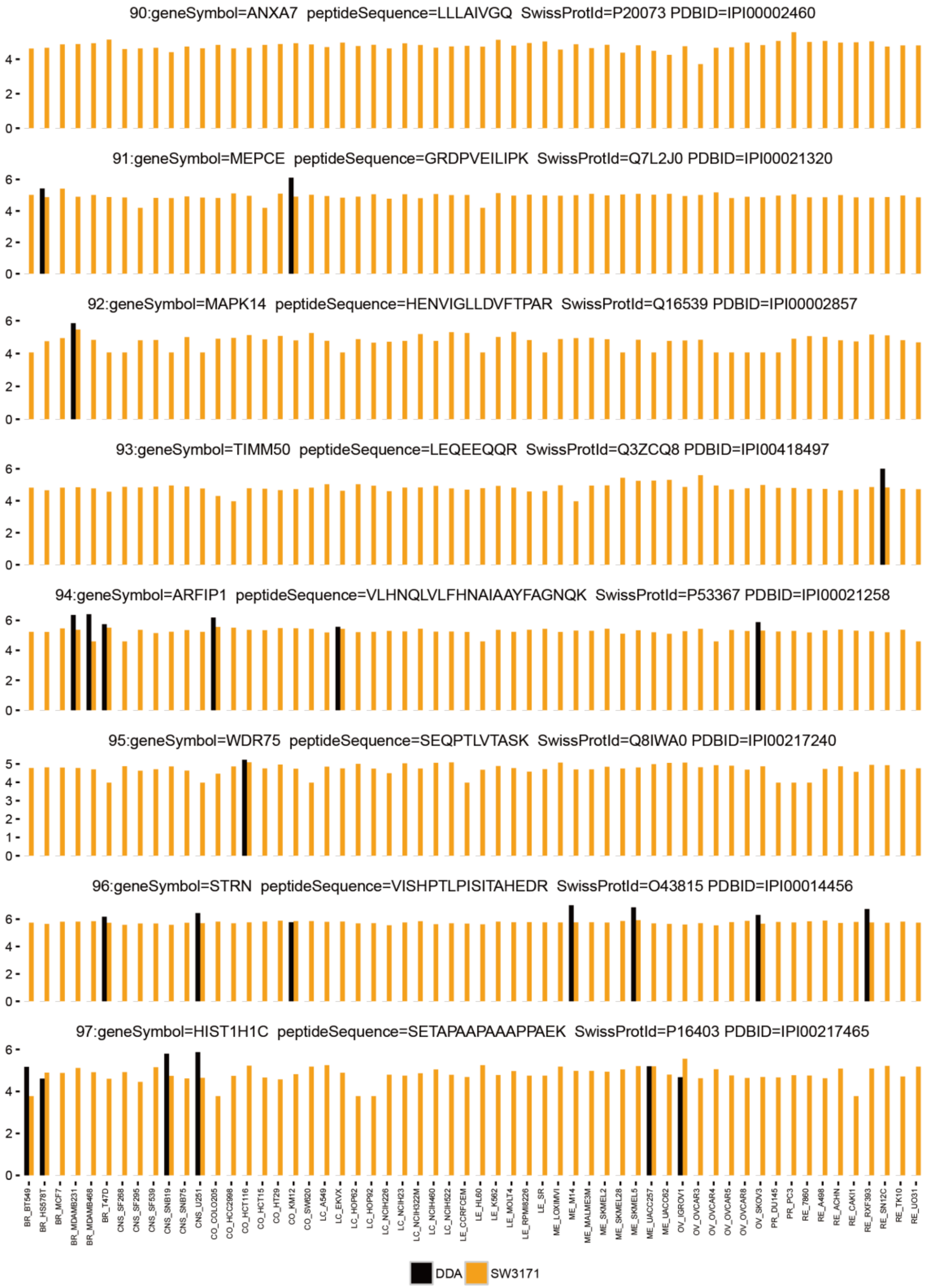
90:geneSymbol=ANXA7 peptideSequence=LLLAIVGQ SwissProtId=P20073 PDBID=IPI00002460
91:geneSymbol=MEPCE peptideSequence=GRDPVEILIPK SwissProtId=Q7L2J0 PDBID=IPI00021320
92:geneSymbol=MAPK14 peptideSequence=HENVIGLLDVFTPAR SwissProtId=Q16539 PDBID=IPI00002857
93:geneSymbol=TIMM50 peptideSequence=LEQEEQQR SwissProtId=Q3ZCQ8 PDBID=IPI00418497
94:geneSymbol=ARFIP1 peptideSequence=VLHNQLVLFHNAIAAYFAGNQK SwissProtId=P53367 PDBID=IPI00021258
95:geneSymbol=WDR75 peptideSequence=SEQPTLVTASK SwissProtId=Q8IWA0 PDBID=IPI00217240
96:geneSymbol=STRN peptideSequence=VISHPTLPISITAHEDR SwissProtId=O43815 PDBID=IPI00014456
97:geneSymbol=HIST1H1C peptideSequence=SETAPAAPAAAPPAEK SwissProtId=P16403 PDBID=IPI00217465
DDA
SW3171

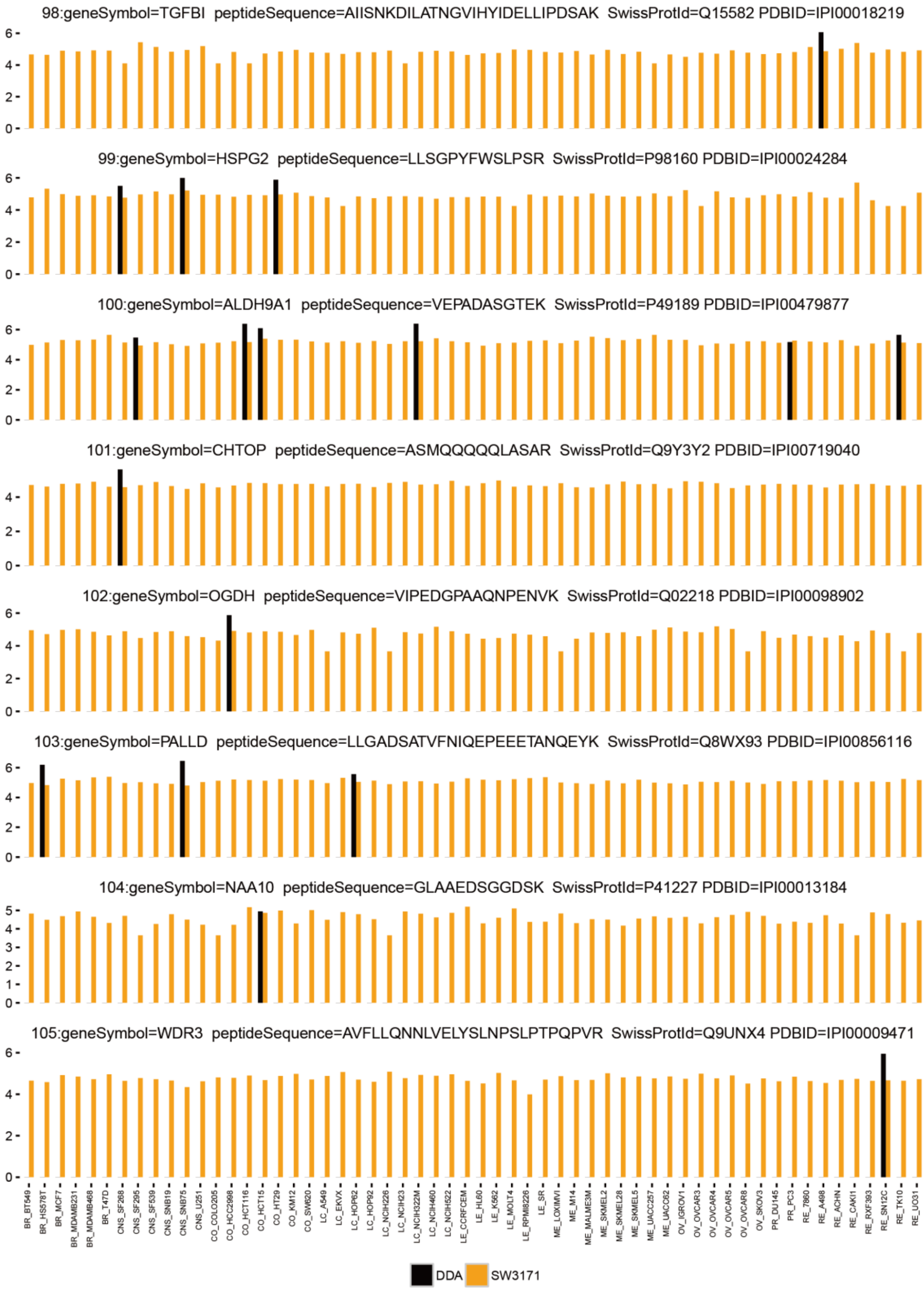
98:geneSymbol=TGFBI peptideSequence=AIISNKDILATNGVIHYIDELLIPDSAK SwissProtId=Q15582 PDBID=IPI00018219
99:geneSymbol=HSPG2 peptideSequence=LLSGPYFWSLPSR SwissProtId=P98160 PDBID=IPI00024284
100:geneSymbol=ALDH9A1 peptideSequence=VEPADASGTEK SwissProtId=P49189 PDBID=IPI00479877
101:geneSymbol=CHTOP peptideSequence=ASMQQQQQLASAR SwissProtId=Q9Y3Y2 PDBID=IPI00719040
102:geneSymbol=OGDH peptideSequence=VIPEDGPAAQNPENVK SwissProtId=Q02218 PDBID=IPI00098902
103:geneSymbol=PALLD peptideSequence=LLGADSATVFNIQEPEEETANQEYK SwissProtId=Q8WX93 PDBID=IPI00856116
104:geneSymbol=NAA10 peptideSequence=GLAAEDSGGDSK SwissProtId=P41227 PDBID=IPI00013184
105:geneSymbol=WDR3 peptideSequence=AVFLLQNNLVELYSLNPSLPTPQPVR SwissProtId=Q9UNX4 PDBID=IPI00009471
DDA
SW3171

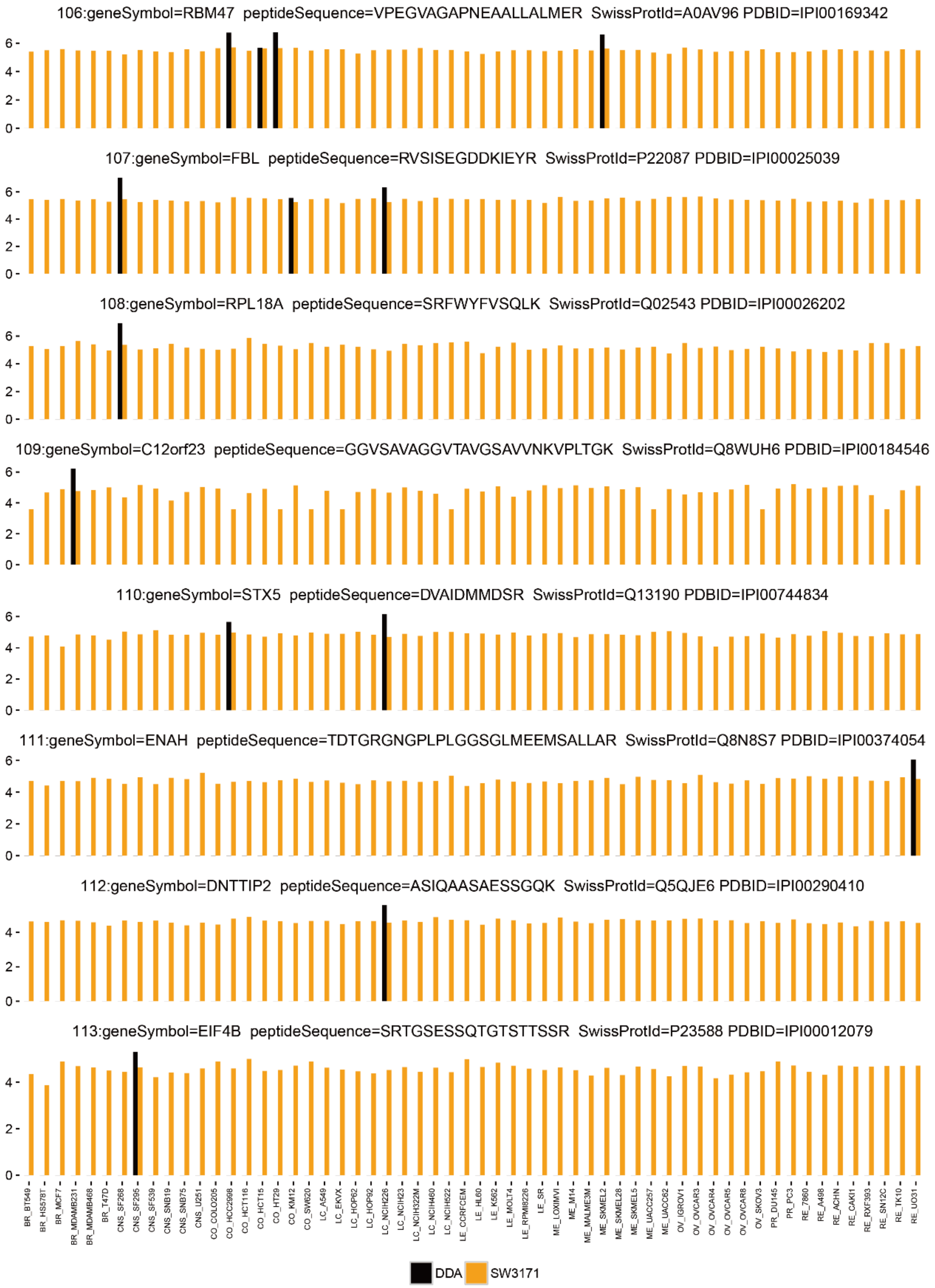
106:geneSymbol=RBM47 peptideSequence=VPEGVAGAPNEAALLALMER SwissProtId=A0AV96 PDBID=IPI00169342
107:geneSymbol=FBL peptideSequence=RVSISEGDDKIEYR SwissProtId=P22087 PDBID=IPI00025039
108:geneSymbol=RPL18A peptideSequence=SRFWYFVSQLK SwissProtId=Q02543 PDBID=IPI00026202
109:geneSymbol=C12orf23 peptideSequence=GGVSAVAGGVTAVGSAVVNKVPLTGK SwissProtId=Q8WUH6 PDBID=IPI00184546
110:geneSymbol=STX5 peptideSequence=DVAIDMMDSR SwissProtId=Q13190 PDBID=IPI00744834
111:geneSymbol=ENAH peptideSequence=TDTGRGNGPLPLGGSGLMEEMSALLAR SwissProtId=Q8N8S7 PDBID=IPI00374054
112:geneSymbol=DNTTIP2 peptideSequence=ASIQAASAESSGQK SwissProtId=Q5QJE6 PDBID=IPI00290410
113:geneSymbol=EIF4B peptideSequence=SRTGSESSQTGTSTTSSR SwissProtId=P23588 PDBID=IPI00012079
DDA
SW3171

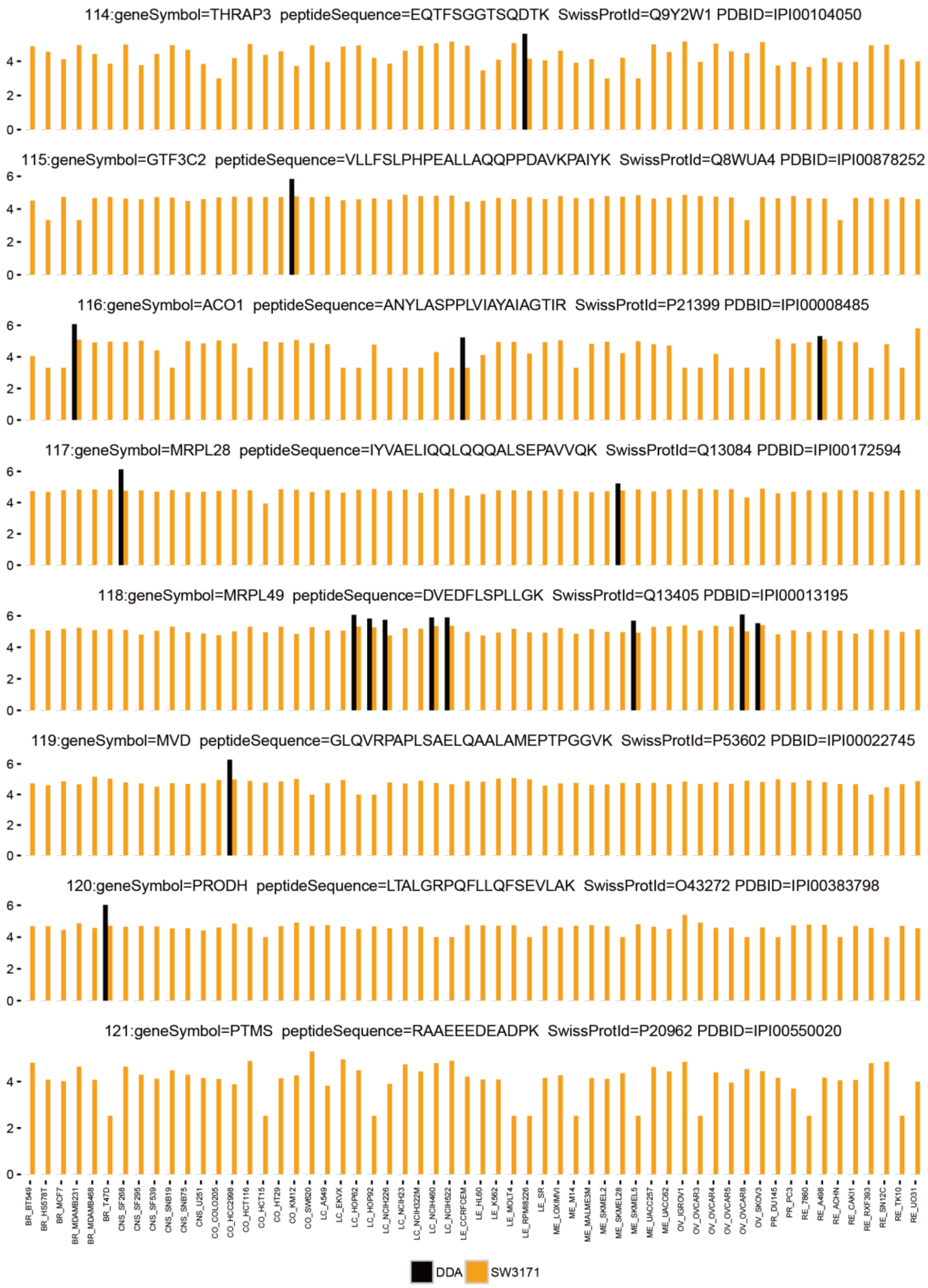
114:geneSymbol=THRAP3 peptideSequence=EQTFSGGTSQDTK SwissProtId=Q9Y2W1 PDBID=IPI00104050
115:geneSymbol=GTF3C2 peptideSequence=VLLFSLPHPEALLAQQPPDAVKPAIYK SwissProtId=Q8WUA4 PDBID=IPI00878252
116:geneSymbol=ACO1 peptideSequence=ANYLASPPLVIAYAIAGTIR SwissProtId=P21399 PDBID=IPI00008485
117:geneSymbol=MRPL28 peptideSequence=IYVAELIQQLQQQALSEPAVVQK SwissProtId=Q13084 PDBID=IPI00172594
118:geneSymbol=MRPL49 peptideSequence=DVEDFLSPLLGK SwissProtId=Q13405 PDBID=IPI00013195
119:geneSymbol=MVD peptideSequence=GLQVRPAPLSAELQAALAMEPTPGGVK SwissProtId=P53602 PDBID=IPI00022745
120:geneSymbol=PRODH peptideSequence=LTALGRPQFLLQFSEVLAK SwissProtId=O43272 PDBID=IPI00383798
121:geneSymbol=PTMS peptideSequence=RAAEEEDEADPK SwissProtId=P20962 PDBID=IPI00550020
0
2
4
6
BR_BT549
BR_HS578T
BR_MCF7
BR_MDAMB231
BR_MDAMB468
BR_T47D
CNS_SF268
CNS_SF295
CNS_SF539
CNS_SNB19
CNS_SNB75
CNS_U251
CO_COLO205
CO_HCC2998
CO_HCT116
CO_HCT15
CO_HT29
CO_KM12
CO_SW620
LC_A549
LC_EKVX
LC_HOP62
LC_HOP92
LC_NCIH226
LC_NCIH23
LC_NCIH322M
LC_NCIH460
LC_NCIH522
LE_CCRFCEM
LE_HL60
LE_K562
LE_MOLT4
LE_RPMI8226
LE_SR
ME_LOXIMVI
ME_M14
ME_MALME3M
ME_SKMEL2
ME_SKMEL28
ME_SKMEL5
ME_UACC257
ME_UACC62
OV_IGROV1
OV_OVCAR3
OV_OVCAR4
OV_OVCAR5
OV_OVCAR8
OV_SKOV3
PR_DU145
PR_PC3
RE_7860
RE_A498
RE_ACHN
RE_CAKI1
RE_RXF393
RE_SN12C
RE_TK10
RE_UO31
DDA
SW3171

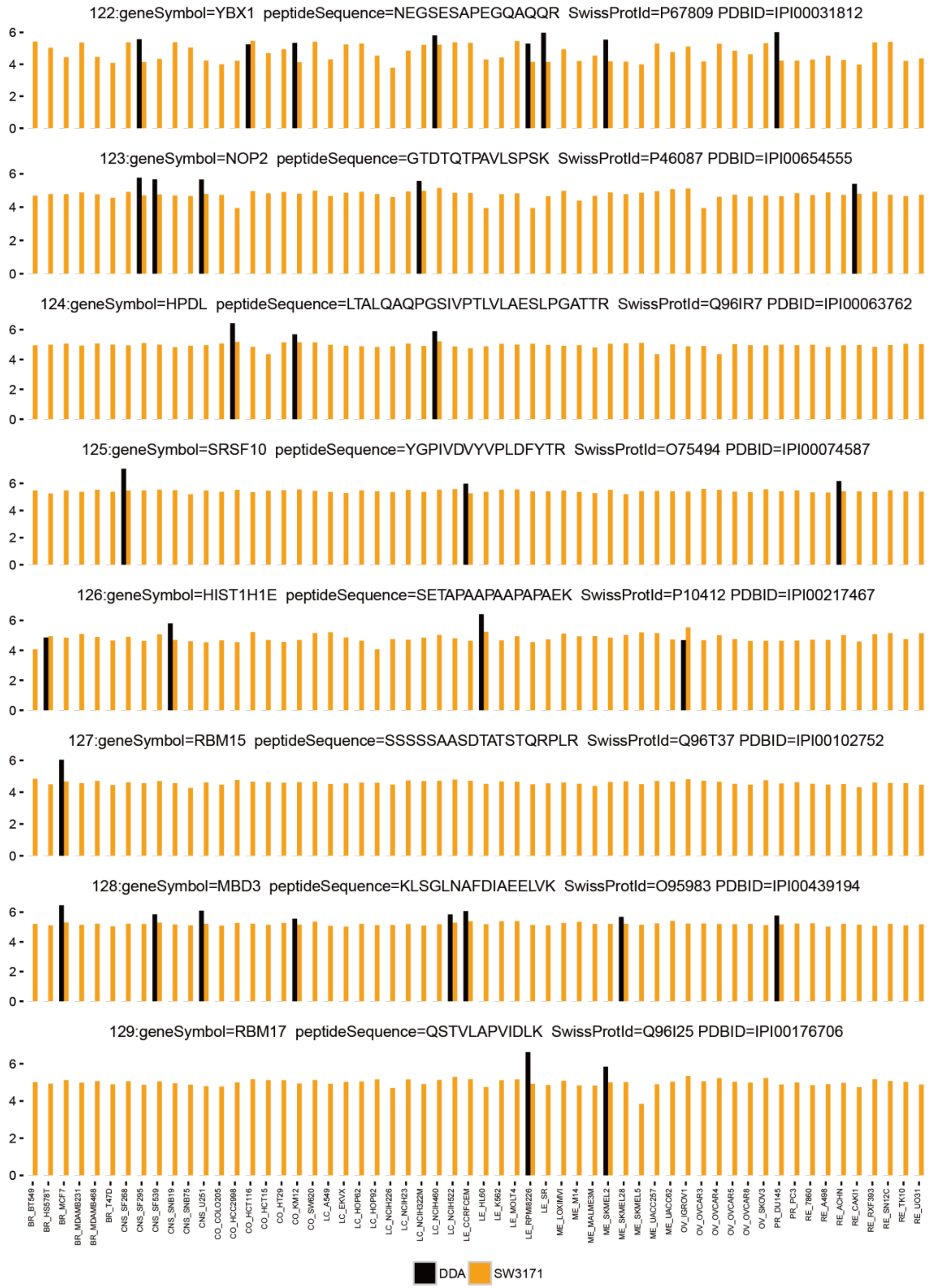
122:geneSymbol=YBX1 peptideSequence=NEGSESAPEGQAQQR SwissProtId=P67809 PDBID=IPI00031812
123:geneSymbol=NOP2 peptideSequence=GTDTQTPAVLSPSK SwissProtId=P46087 PDBID=IPI00654555
124:geneSymbol=HPDL peptideSequence=LTALQAQPGSIVPTLVLAESLPGATTR SwissProtId=Q96IR7 PDBID=IPI00063762
125:geneSymbol=SRSF10 peptideSequence=YGPIVDVYVPLDFYTR SwissProtId=O75494 PDBID=IPI00074587
126:geneSymbol=HIST1H1E peptideSequence=SETAPAAPAAPAPAEK SwissProtId=P10412 PDBID=IPI00217467
127:geneSymbol=RBM15 peptideSequence=SSSSSAASDTATSTQRPLR SwissProtId=Q96T37 PDBID=IPI00102752
128:geneSymbol=MBD3 peptideSequence=KLSGLNAFDIAEELVK SwissProtId=O95983 PDBID=IPI00439194
129:geneSymbol=RBM17 peptideSequence=QSTVLAPVIDLK SwissProtId=Q96I25 PDBID=IPI00176706
0
2
4
6
BR_BT549
BR_HS578T
BR_MCF7
BR_MDAMB231
BR_MDAMB468
BR_T47D
CNS_SF268
CNS_SF295
CNS_SF539
CNS_SNB19
CNS_SNB75
CNS_U251
CO_COLO205
CO_HCC2998
CO_HCT116
CO_HCT15
CO_HT29
CO_KM12
CO_SW620
LC_A549
LC_EKVX
LC_HOP62
LC_HOP92
LC_NCIH226
LC_NCIH23
LC_NCIH322M
LC_NCIH460
LC_NCIH522
LE_CCRFCEM
LE_HL60
LE_K562
LE_MOLT4
LE_RPMI8226
LE_SR
ME_LOXIMVI
ME_M14
ME_MALME3M
ME_SKMEL2
ME_SKMEL28
ME_SKMEL5
ME_UACC257
ME_UACC62
OV_IGROV1
OV_OVCAR3
OV_OVCAR4
OV_OVCAR5
OV_OVCAR8
OV_SKOV3
PR_DU145
PR_PC3
RE_7860
RE_A498
RE_ACHN
RE_CAKI1
RE_RXF393
RE_SN12C
RE_TK10
RE_UO31
DDA
SW3171

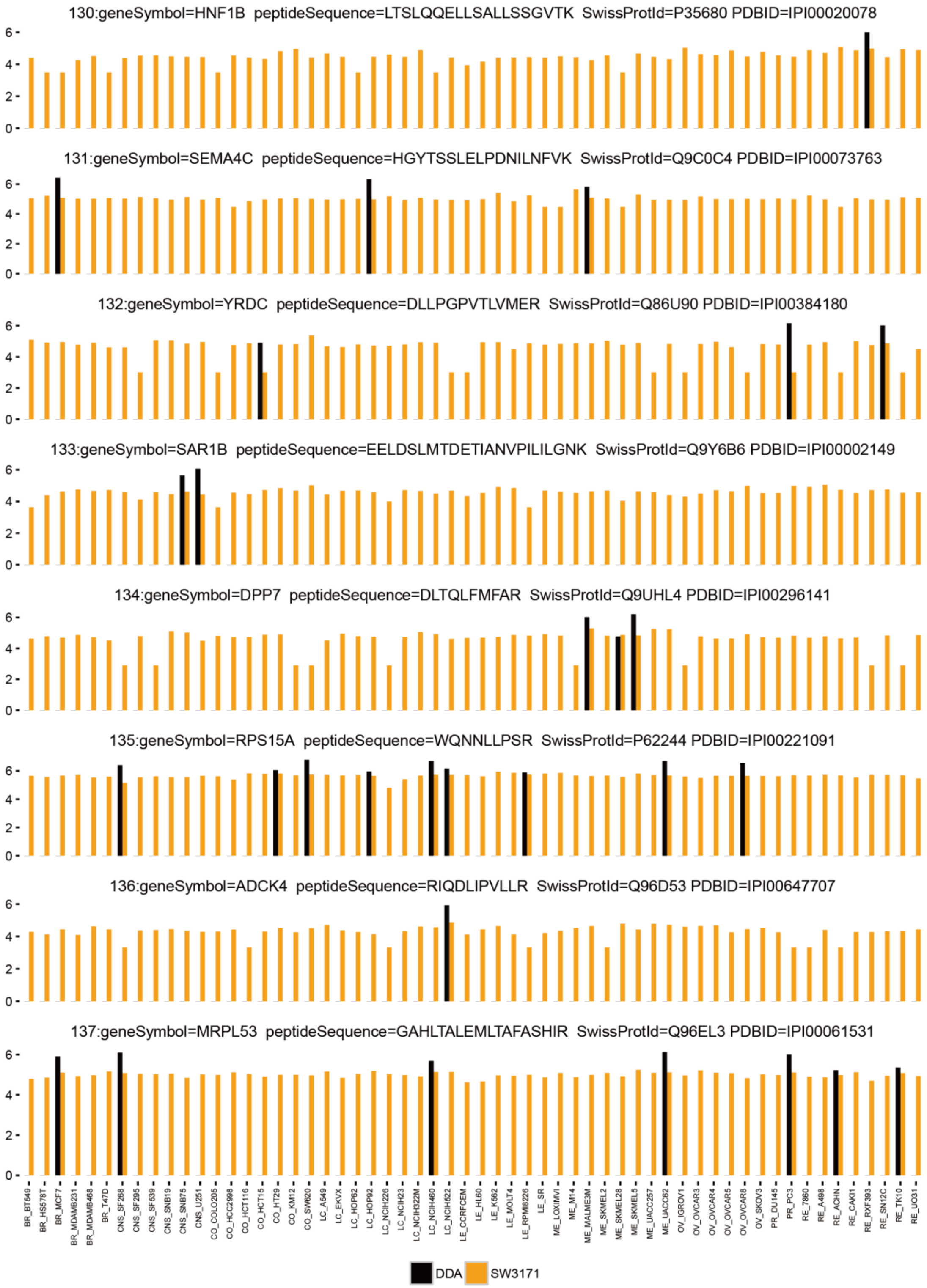
130:geneSymbol=HNF1B peptideSequence=LTSLQQELLSALLSSGVTK SwissProtId=P35680 PDBID=IPI00020078
131:geneSymbol=SEMA4C peptideSequence=HGYTSSLELPDNILNFVK SwissProtId=Q9C0C4 PDBID=IPI00073763
132:geneSymbol=YRDC peptideSequence=DLLPGPVTLVMER SwissProtId=Q86U90 PDBID=IPI00384180
133:geneSymbol=SAR1B peptideSequence=EELDSLMTDETIANVPILILGNK SwissProtId=Q9Y6B6 PDBID=IPI00002149
134:geneSymbol=DPP7 peptideSequence=DLTQLFMFAR SwissProtId=Q9UHL4 PDBID=IPI00296141
135:geneSymbol=RPS15A peptideSequence=WQNNLLPSR SwissProtId=P62244 PDBID=IPI00221091
136:geneSymbol=ADCK4 peptideSequence=RIQDLIPVLLR SwissProtId=Q96D53 PDBID=IPI00647707
137:geneSymbol=MRPL53 peptideSequence=GAHLTALEMLTAFASHIR SwissProtId=Q96EL3 PDBID=IPI00061531
0
2
4
6
BR_BT549
BR_HS578T
BR_MCF7
BR_MDAMB231
BR_MDAMB468
BR_T47D
CNS_SF268
CNS_SF295
CNS_SF539
CNS_SNB19
CNS_SNB75
CNS_U251
CO_COLO205
CO_HCC2998
CO_HCT116
CO_HCT15
CO_HT29
CO_KM12
CO_SW620
LC_A549
LC_EKVX
LC_HOP62
LC_HOP92
LC_NCIH226
LC_NCIH23
LC_NCIH322M
LC_NCIH460
LC_NCIH522
LE_CCRFCEM
LE_HL60
LE_K562
LE_MOLT4
LE_RPMI8226
LE_SR
ME_LOXIMVI
ME_M14
ME_MALME3M
ME_SKMEL2
ME_SKMEL28
ME_SKMEL5
ME_UACC257
ME_UACC62
OV_IGROV1
OV_OVCAR3
OV_OVCAR4
OV_OVCAR5
OV_OVCAR8
OV_SKOV3
PR_DU145
PR_PC3
RE_7860
RE_A498
RE_ACHN
RE_CAKI1
RE_RXF393
RE_SN12C
RE_TK10
RE_UO31
DDA
SW3171

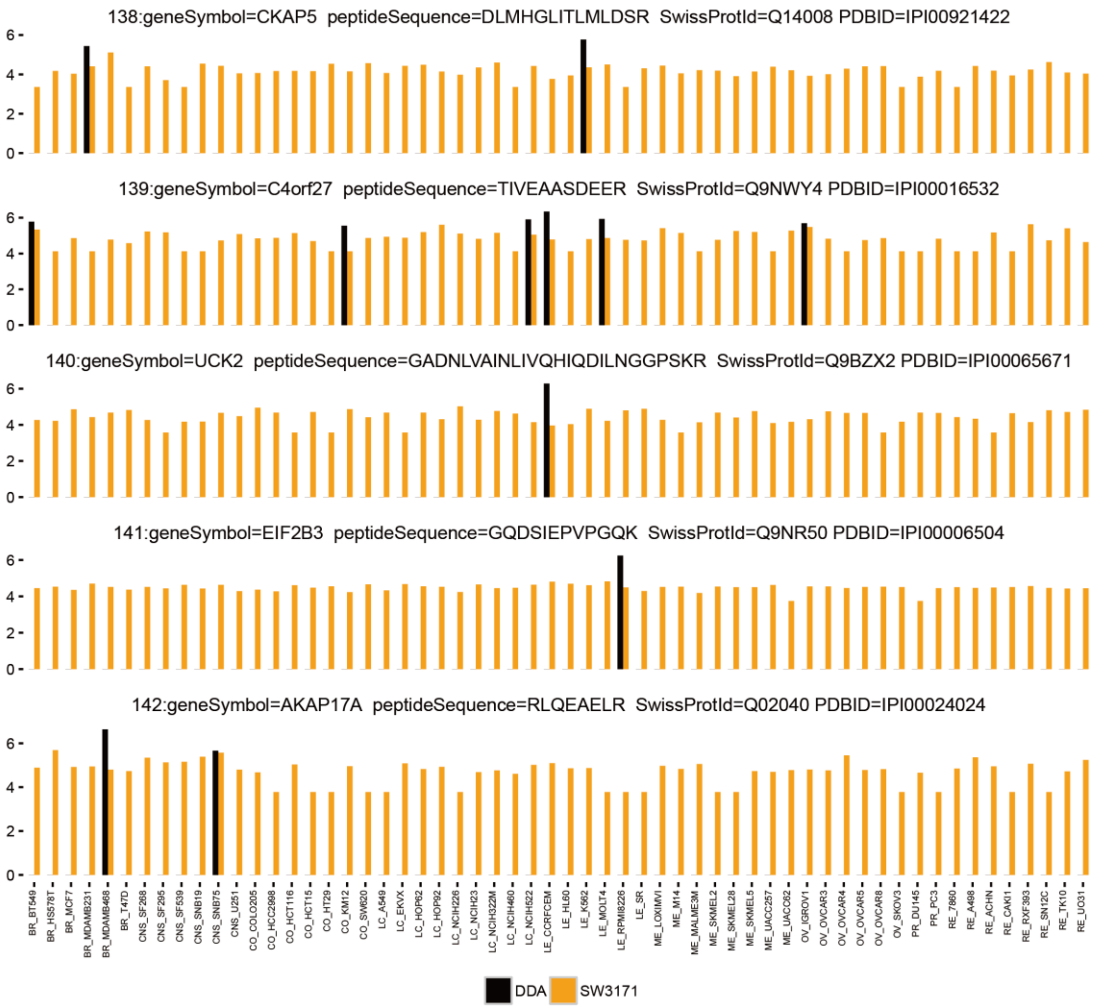
138:geneSymbol=CKAP5 peptideSequence=DLMHGLITLMLDSR SwissProtId=Q14008 PDBID=IPI00921422
139:geneSymbol=C4orf27 peptideSequence=TIVEAASDEER SwissProtId=Q9NWY4 PDBID=IPI00016532
140:geneSymbol=UCK2 peptideSequence=GADNLVAINLIVQHIQDILNGGPSKR SwissProtId=Q9BZX2 PDBID=IPI00065671
141:geneSymbol=EIF2B3 peptideSequence=GQDSIEPVPGQK SwissProtId=Q9NR50 PDBID=IPI00006504
142:geneSymbol=AKAP17A peptideSequence=RLQEAELR SwissProtId=Q02040 PDBID=IPI00024024
DDA
SW3171

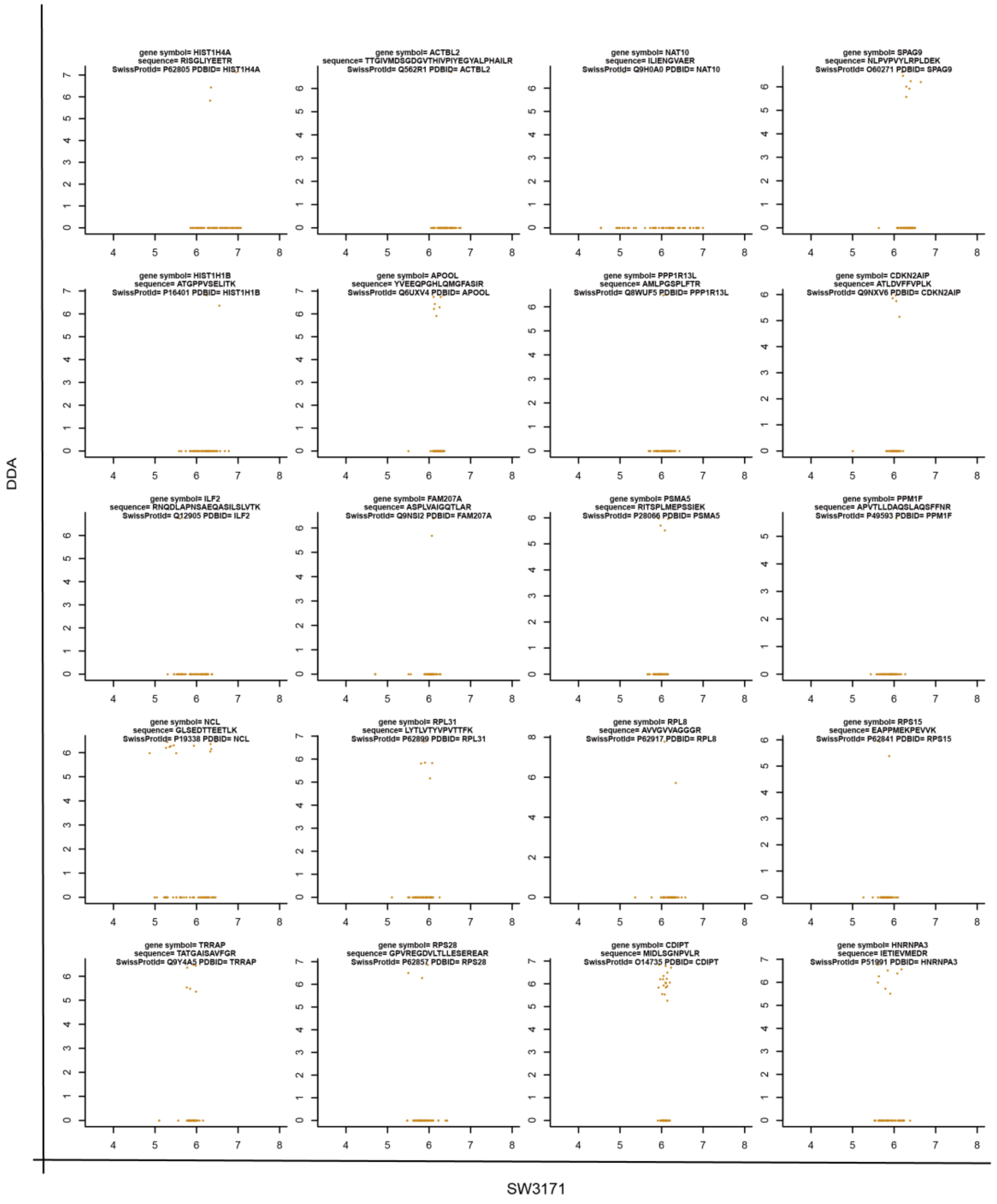
gene symbol= HIST1H4A
sequence= RISGLIYEETR
SwissProtId= P62805 PDBID= HIST1H4A
gene symbol= ACTBL2
sequence= TTGIVMDSGDGVTHIVPIYEGYALPHAILR
SwissProtId= Q562R1 PDBID= ACTBL2
gene symbol= NAT10
sequence= ILIENGVAER
SwissProtId= Q9H0A0 PDBID= NAT10
gene symbol= SPAG9
sequence= NLPVPVYLRPLDEK
SwissProtId= O60271 PDBID= SPAG9
gene symbol= HIST1H1B
sequence= ATGPPVSELITK
SwissProtId= P16401 PDBID= HIST1H1B
gene symbol= APOOL
sequence= YVEEQPGHLQMGFASIR
SwissProtId= Q6UXV4 PDBID= APOOL
gene symbol= PPP1R13L
sequence= AMLPGSPLFTR
SwissProtId= Q8WUF5 PDBID= PPP1R13L
gene symbol= CDKN2AIP
sequence= ATLDVFFVPLK
SwissProtId= Q9NXV6 PDBID= CDKN2AIP
gene symbol= ILF2
sequence= RNQDLAPNSAEQASILSLVTK
SwissProtId= Q12905 PDBID= ILF2
gene symbol= FAM207A
sequence= ASPLVAIGQTLAR
SwissProtId= Q9NSI2 PDBID= FAM207A
gene symbol= PSMA5
sequence= RITSPLMEPSSIEK
SwissProtId= P28066 PDBID= PSMA5
gene symbol= PPM1F
sequence= APVTLLDAQSLAQSFFNR
SwissProtId= P49593 PDBID= PPM1F
gene symbol= NCL
sequence= GLSEDTTEETLK
SwissProtId= P19338 PDBID= NCL
gene symbol= RPL31
sequence= LYTLVTYVPVTTFK
SwissProtId= P62899 PDBID= RPL31
gene symbol= RPL8
sequence= AVVGVVAGGGR
SwissProtId= P62917 PDBID= RPL8
gene symbol= RPS15
sequence= EAPPMEKPEVVK
SwissProtId= P62841 PDBID= RPS15
gene symbol= TRRAP
sequence= TATGAISAVFGR
SwissProtId= Q9Y4A5 PDBID= TRRAP
gene symbol= RPS28
sequence= GPVREGDVLTLLESEREAR
SwissProtId= P62857 PDBID= RPS28
gene symbol= CDIPT
sequence= MIDLSGNPVLR
SwissProtId= O14735 PDBID= CDIPT
gene symbol= HNRNPA3
sequence= IETIEVMEDR
SwissProtId= P51991 PDBID= HNRNPA3
DDA
SW3171

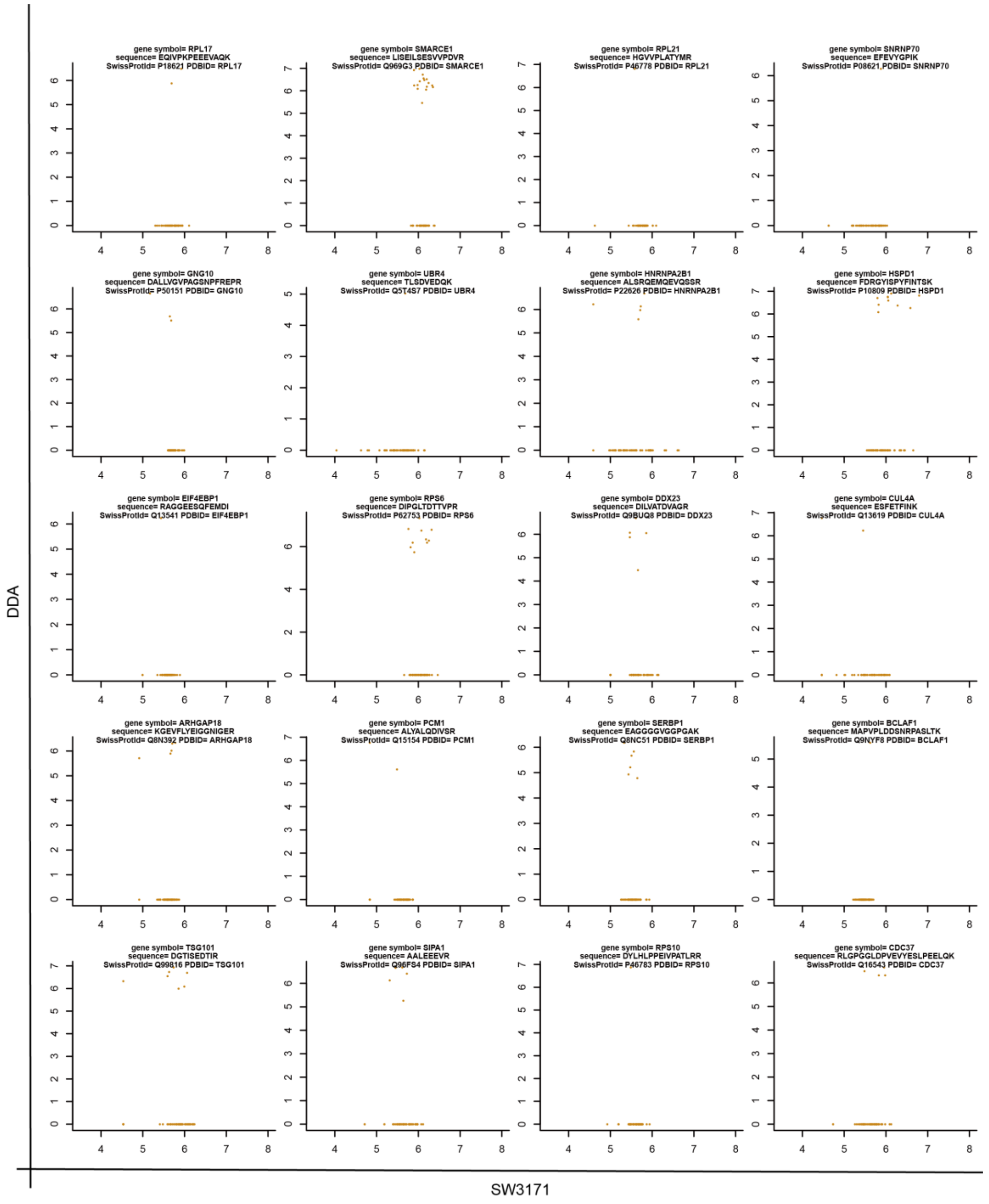
gene symbol= RPL17
sequence= EQIVPKPEEEVAQK
SwissProtId= P18621 PDBID= RPL17
gene symbol= SMARCE1
sequence= LISEILSESVVPDVR
SwissProtId= Q969G3 PDBID= SMARCE1
gene symbol= RPL21
sequence= HGVVPLATYMR
SwissProtId= P46778 PDBID= RPL21
gene symbol= SNRNP70
sequence= EFEVYGPIK
SwissProtId= P08621 PDBID= SNRNP70
gene symbol= GNG10
sequence= DALLVGVPAGSNPFREPR
SwissProtId= P50151 PDBID= GNG10
gene symbol= UBR4
sequence= TLSDVEDQK
SwissProtId= Q5T4S7 PDBID= UBR4
gene symbol= HNRNPA2B1
sequence= ALSRQEMQEVQSSR
SwissProtId= P22626 PDBID= HNRNPA2B1
gene symbol= HSPD1
sequence= FDRGYISPYFINTSK
SwissProtId= P10809 PDBID= HSPD1
gene symbol= EIF4EBP1
sequence= RAGGEESQFEMDI
SwissProtId= Q13541 PDBID= EIF4EBP1
gene symbol= RPS6
sequence= DIPGLTDTTVPR
SwissProtId= P62753 PDBID= RPS6
gene symbol= DDX23
sequence= DILVATDVAGR
SwissProtId= Q9BUQ8 PDBID= DDX23
gene symbol= CUL4A
sequence= ESFETFINK
SwissProtId= Q13619 PDBID= CUL4A
gene symbol= ARHGAP18
sequence= KGEVFLYEIGGNIGER
SwissProtId= Q8N392 PDBID= ARHGAP18
gene symbol= PCM1
sequence= ALYALQDIVSR
SwissProtId= Q15154 PDBID= PCM1
gene symbol= SERBP1
sequence= EAGGGGVGGPGAK
SwissProtId= Q8NC51 PDBID= SERBP1
gene symbol= BCLAF1
sequence= MAPVPLDDSNRPASLTK
SwissProtId= Q9NYF8 PDBID= BCLAF1
gene symbol= TSG101
sequence= DGTISEDTIR
SwissProtId= Q99816 PDBID= TSG101
gene symbol= SIPA1
sequence= AALEEEVR
SwissProtId= Q96FS4 PDBID= SIPA1
gene symbol= RPS10
sequence= DYLHLPPEIVPATLRR
SwissProtId= P46783 PDBID= RPS10
gene symbol= CDC37
sequence= RLGPGGLDPVEVYESLPEELQK
SwissProtId= Q16543 PDBID= CDC37
DDA
SW3171

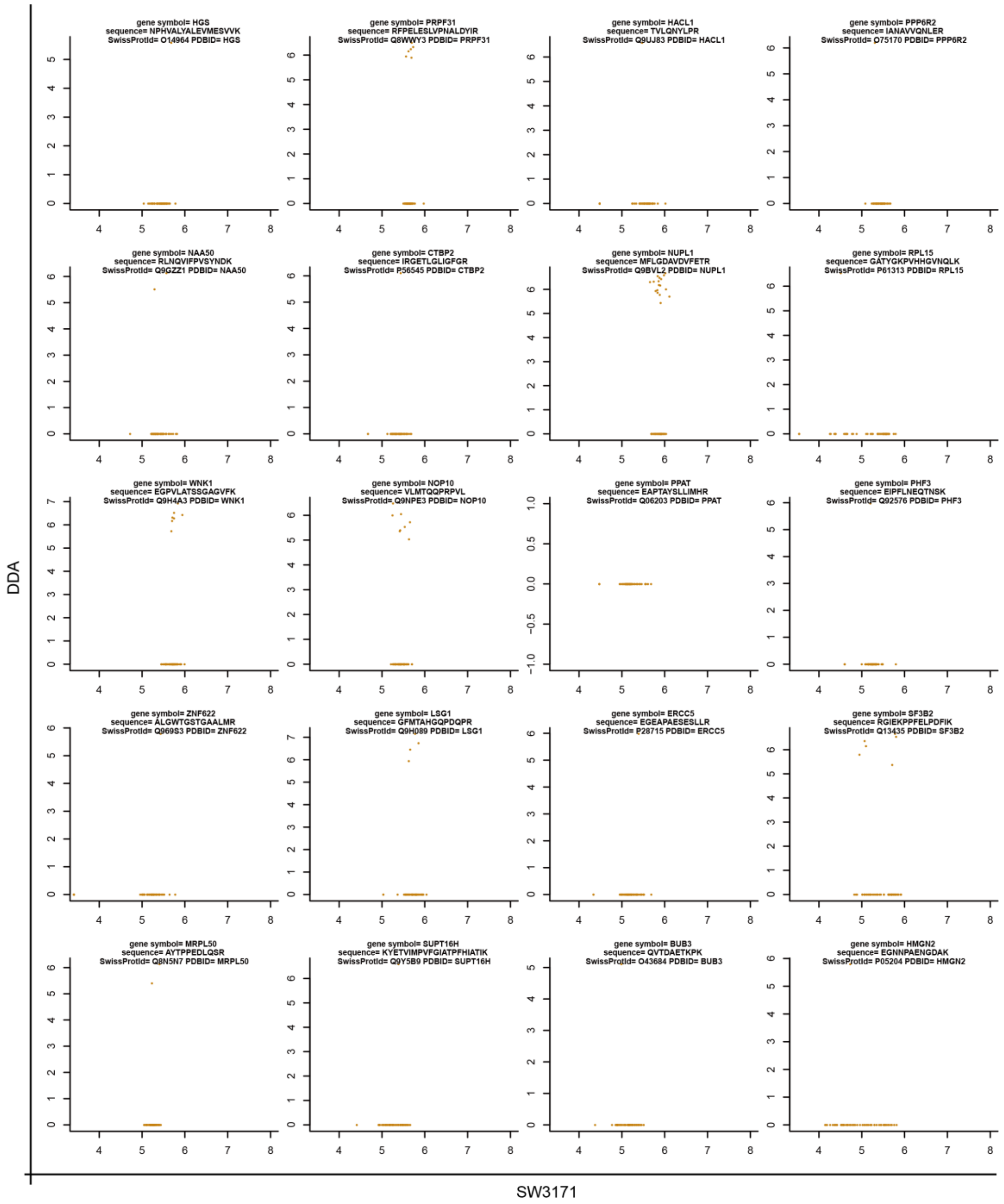
gene symbol= HGS
sequence= NPHVALYALEVMESVVK
SwissProtId= O14964 PDBID= HGS
gene symbol= PRPF31
sequence= RFPELESLVPNALDYIR
SwissProtId= Q8WWY3 PDBID= PRPF31
gene symbol= HACL1
sequence= TVLQNYLPR
SwissProtId= Q9UJ83 PDBID= HACL1
gene symbol= PPP6R2
sequence= IANAVVQNLER
SwissProtId= O75170 PDBID= PPP6R2
gene symbol= NAA50
sequence= RLNQVIFPVSYNDK
SwissProtId= Q9GZZ1 PDBID= NAA50
gene symbol= CTBP2
sequence= IRGETLGLIGFGR
SwissProtId= P56545 PDBID= CTBP2
gene symbol= NUPL1
sequence= MFLGDAVDVFETR
SwissProtId= Q9BVL2 PDBID= NUPL1
gene symbol= RPL15
sequence= GATYGKPVHHGVNQLK
SwissProtId= P61313 PDBID= RPL15
gene symbol= WNK1
sequence= EGPVLATSSGAGVFK
SwissProtId= Q9H4A3 PDBID= WNK1
gene symbol= NOP10
sequence= VLMTQQPRPVL
SwissProtId= Q9NPE3 PDBID= NOP10
gene symbol= PPAT
sequence= EAPTAYSLLIMHR
SwissProtId= Q06203 PDBID= PPAT
gene symbol= PHF3
sequence= EIPFLNEQTNSK
SwissProtId= Q92576 PDBID= PHF3
gene symbol= ZNF622
sequence= ALGWTGSTGAALMR
SwissProtId= Q969S3 PDBID= ZNF622
gene symbol= LSG1
sequence= GFMTAHGQPDQPR
SwissProtId= Q9H089 PDBID= LSG1
gene symbol= ERCC5
sequence= EGEAPAESESLLR
SwissProtId= P28715 PDBID= ERCC5
gene symbol= SF3B2
sequence= RGIEKPPFELPDFIK
SwissProtId= Q13435 PDBID= SF3B2
gene symbol= MRPL50
sequence= AYTPPEDLQSR
SwissProtId= Q8N5N7 PDBID= MRPL50
gene symbol= SUPT16H
sequence= KYETVIMPVFGIATPFHIATIK
SwissProtId= Q9Y5B9 PDBID= SUPT16H
gene symbol= BUB3
sequence= QVTDAETKPK
SwissProtId= O43684 PDBID= BUB3
gene symbol= HMGN2
sequence= EGNNPAENGDAK
SwissProtId= P05204 PDBID= HMGN2
DDA
SW3171

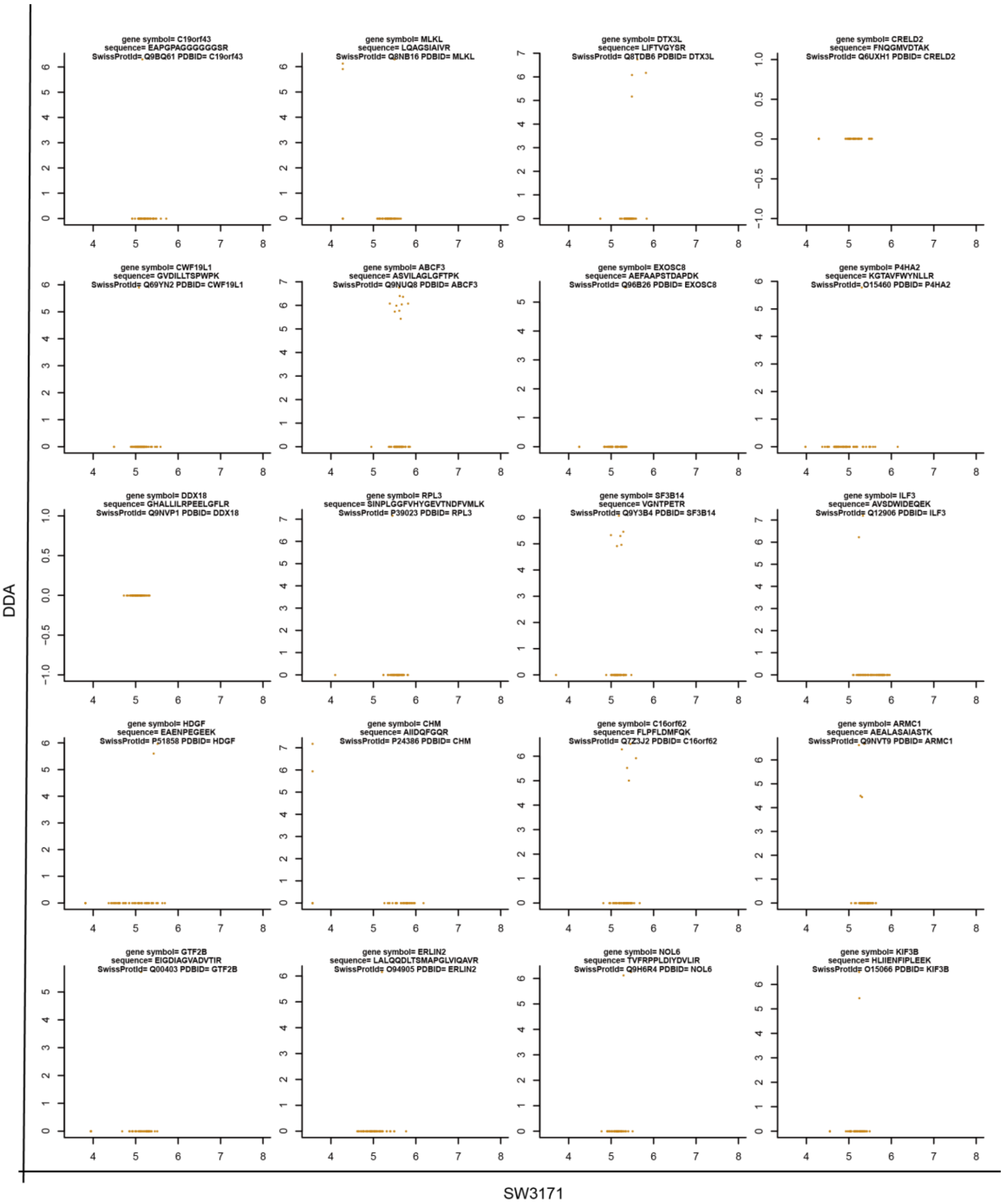
gene symbol= C19orf43
sequence= EAPGPAGGGGGGSR
SwissProtId= Q9BQ61 PDBID= C19orf43
gene symbol= MLKL
sequence= LQAGSIAIVR
SwissProtId= Q8NB16 PDBID= MLKL
gene symbol= DTX3L
sequence= LIFTVGYSR
SwissProtId= Q8TDB6 PDBID= DTX3L
gene symbol= CRELD2
sequence= FNQGMVDTAK
SwissProtId= Q6UXH1 PDBID= CRELD2
gene symbol= CWF19L1
sequence= GVDILLTSPWPK
SwissProtId= Q69YN2 PDBID= CWF19L1
gene symbol= ABCF3
sequence= ASVILAGLGFTPK
SwissProtId= Q9NUQ8 PDBID= ABCF3
gene symbol= EXOSC8
sequence= AEFAAPSTDAPDK
SwissProtId= Q96B26 PDBID= EXOSC8
gene symbol= P4HA2
sequence= KGTAVFWYNLLR
SwissProtId= O15460 PDBID= P4HA2
gene symbol= DDX18
sequence= GHALLILRPEELGFLR
SwissProtId= Q9NVP1 PDBID= DDX18
gene symbol= RPL3
sequence= SINPLGGFVHYGEVTNDFVMLK
SwissProtId= P39023 PDBID= RPL3
gene symbol= SF3B14
sequence= VGNTPETR
SwissProtId= Q9Y3B4 PDBID= SF3B14
gene symbol= ILF3
sequence= AVSDWIDEQEK
SwissProtId= Q12906 PDBID= ILF3
gene symbol= HDGF
sequence= EAENPEGEEK
SwissProtId= P51858 PDBID= HDGF
gene symbol= CHM
sequence= AIIDQFGQR
SwissProtId= P24386 PDBID= CHM
gene symbol= C16orf62
sequence= FLPFLDMFQK
SwissProtId= Q7Z3J2 PDBID= C16orf62
gene symbol= ARMC1
sequence= AEALASAIASTK
SwissProtId= Q9NVT9 PDBID= ARMC1
gene symbol= GTF2B
sequence= EIGDIAGVADVTIR
SwissProtId= Q00403 PDBID= GTF2B
gene symbol= ERLIN2
sequence= LALQQDLTSMAPGLVIQAVR
SwissProtId= O94905 PDBID= ERLIN2
gene symbol= NOL6
sequence= TVFRPPLDIYDVLIR
SwissProtId= Q9H6R4 PDBID= NOL6
gene symbol= KIF3B
sequence= HLIIENFIPLEEK
SwissProtId= O15066 PDBID= KIF3B
DDA
SW3171

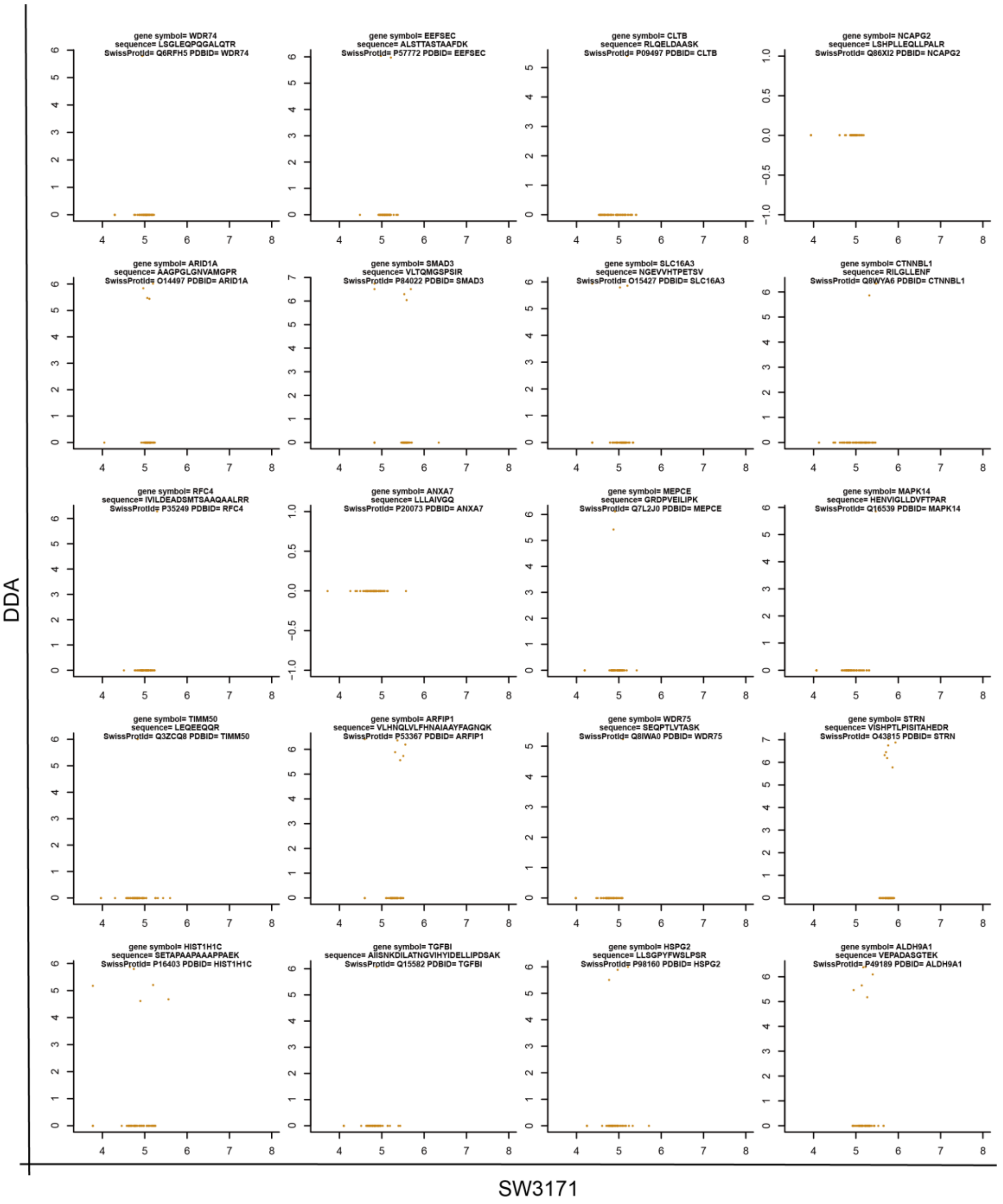
gene symbol= WDR74
sequence= LSGLEQPQGALQTR
SwissProtId= Q6RFH5 PDBID= WDR74
gene symbol= EEFSEC
sequence= ALSTTASTAAFDK
SwissProtId= P57772 PDBID= EEFSEC
gene symbol= CLTB
sequence= RLQELDAASK
SwissProtId= P09497 PDBID= CLTB
gene symbol= NCAPG2
sequence= LSHPLLEQLLPALR
SwissProtId= Q86XI2 PDBID= NCAPG2
gene symbol= ARID1A
sequence= AAGPGLGNVAMGPR
SwissProtId= O14497 PDBID= ARID1A
gene symbol= SMAD3
sequence= VLTQMGSPSIR
SwissProtId= P84022 PDBID= SMAD3
gene symbol= SLC16A3
sequence= NGEVVHTPETSV
SwissProtId= O15427 PDBID= SLC16A3
gene symbol= CTNNBL1
sequence= RILGLLENF
SwissProtId= Q8WYA6 PDBID= CTNNBL1
gene symbol= RFC4
sequence= IVILDEADSMTSAAQAALRR
SwissProtId= P35249 PDBID= RFC4
gene symbol= ANXA7
sequence= LLLAIVGQ
SwissProtId= P20073 PDBID= ANXA7
gene symbol= MEPCE
sequence= GRDPVEILIPK
SwissProtId= Q7L2J0 PDBID= MEPCE
gene symbol= MAPK14
sequence= HENVIGLLDVFTPAR
SwissProtId= Q16539 PDBID= MAPK14
gene symbol= TIMM50
sequence= LEQEEQQR
SwissProtId= Q3ZCQ8 PDBID= TIMM50
gene symbol= ARFIP1
sequence= VLHNQLVLFHNAIAAYFAGNQK
SwissProtId= P53367 PDBID= ARFIP1
gene symbol= WDR75
sequence= SEQPTLVTASK
SwissProtId= Q8IWA0 PDBID= WDR75
gene symbol= STRN
sequence= VISHPTLPISITAHEDR
SwissProtId= O43815 PDBID= STRN
gene symbol= HIST1H1C
sequence= SETAPAAPAAAPPAEK
SwissProtId= P16403 PDBID= HIST1H1C
gene symbol= TGFBI
sequence= AIISNKDILATNGVIHYIDELLIPDSAK
SwissProtId= Q15582 PDBID= TGFBI
gene symbol= HSPG2
sequence= LLSGPYFWSLPSR
SwissProtId= P98160 PDBID= HSPG2
gene symbol= ALDH9A1
sequence= VEPADASGTEK
SwissProtId= P49189 PDBID= ALDH9A1
DDA
SW3171

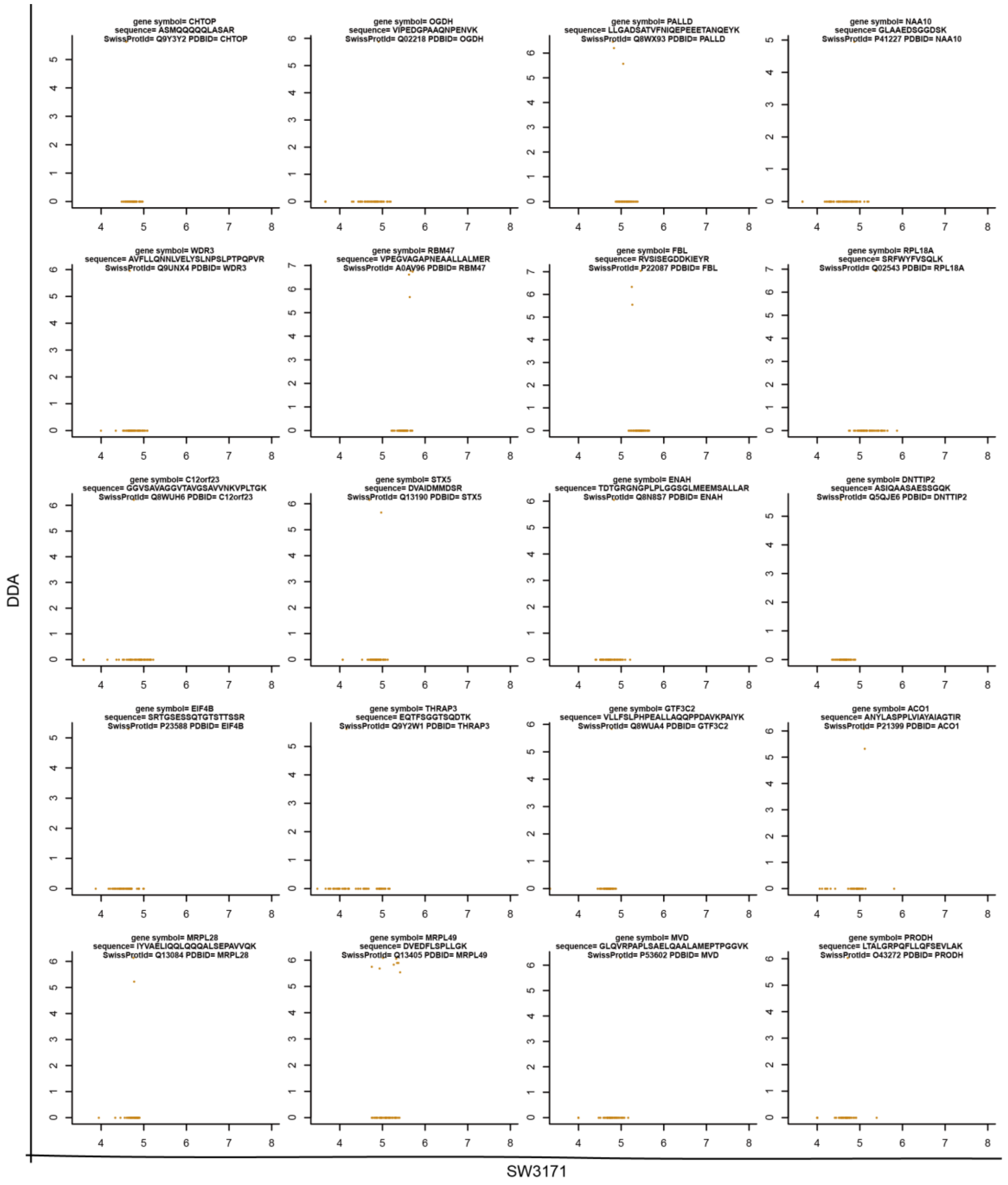
gene symbol= CHTOP
sequence= ASMQQQQQLASAR
SwissProtId= Q9Y3Y2 PDBID= CHTOP
gene symbol= OGDH
sequence= VIPEDGPAAQNPENVK
SwissProtId= Q02218 PDBID= OGDH
gene symbol= PALLD
sequence= LLGADSATVFNIQEPEEETANQEYK
SwissProtId= Q8WX93 PDBID= PALLD
gene symbol= NAA10
sequence= GLAAEDSGGDSK
SwissProtId= P41227 PDBID= NAA10
gene symbol= WDR3
sequence= AVFLLQNNLVELYSLNPSLPTPQPVR
SwissProtId= Q9UNX4 PDBID= WDR3
gene symbol= RBM47
sequence= VPEGVAGAPNEAALLALMER
SwissProtId= A0AV96 PDBID= RBM47
gene symbol= FBL
sequence= RVSISEGDDKIEYR
SwissProtId= P22087 PDBID= FBL
gene symbol= RPL18A
sequence= SRFWYFVSQLK
SwissProtId= Q02543 PDBID= RPL18A
gene symbol= C12orf23
sequence= GGVSAVAGGVTAVGSAVVNKVPLTGK
SwissProtId= Q8WUH6 PDBID= C12orf23
gene symbol= STX5
sequence= DVAIDMMDSR
SwissProtId= Q13190 PDBID= STX5
gene symbol= ENAH
sequence= TDTGRGNGPLPLGGSGLMEEMSALLAR
SwissProtId= Q8N8S7 PDBID= ENAH
gene symbol= DNTTIP2
sequence= ASIQAASAESSGQK
SwissProtId= Q5QJE6 PDBID= DNTTIP2
gene symbol= EIF4B
sequence= SRTGSESSQTGTSTTSSR
SwissProtId= P23588 PDBID= EIF4B
gene symbol= THRAP3
sequence= EQTFSGGTSQDTK
SwissProtId= Q9Y2W1 PDBID= THRAP3
gene symbol= GTF3C2
sequence= VLLFSLPHPEALLAQQPPDAVKPAIYK
SwissProtId= Q8WUA4 PDBID= GTF3C2
gene symbol= ACO1
sequence= ANYLASPPLVIAYAIAGTIR
SwissProtId= P21399 PDBID= ACO1
gene symbol= MRPL28
sequence= IYVAELIQQLQQQALSEPAVVQK
SwissProtId= Q13084 PDBID= MRPL28
gene symbol= MRPL49
sequence= DVEDFLSPLLGK
SwissProtId= Q13405 PDBID= MRPL49
gene symbol= MVD
sequence= GLQVRPAPLSAELQAALAMEPTPGGVK
SwissProtId= P53602 PDBID= MVD
gene symbol= PRODH
sequence= LTALGRPQFLLQFSEVLAK
SwissProtId= O43272 PDBID= PRODH
DDA
SW3171

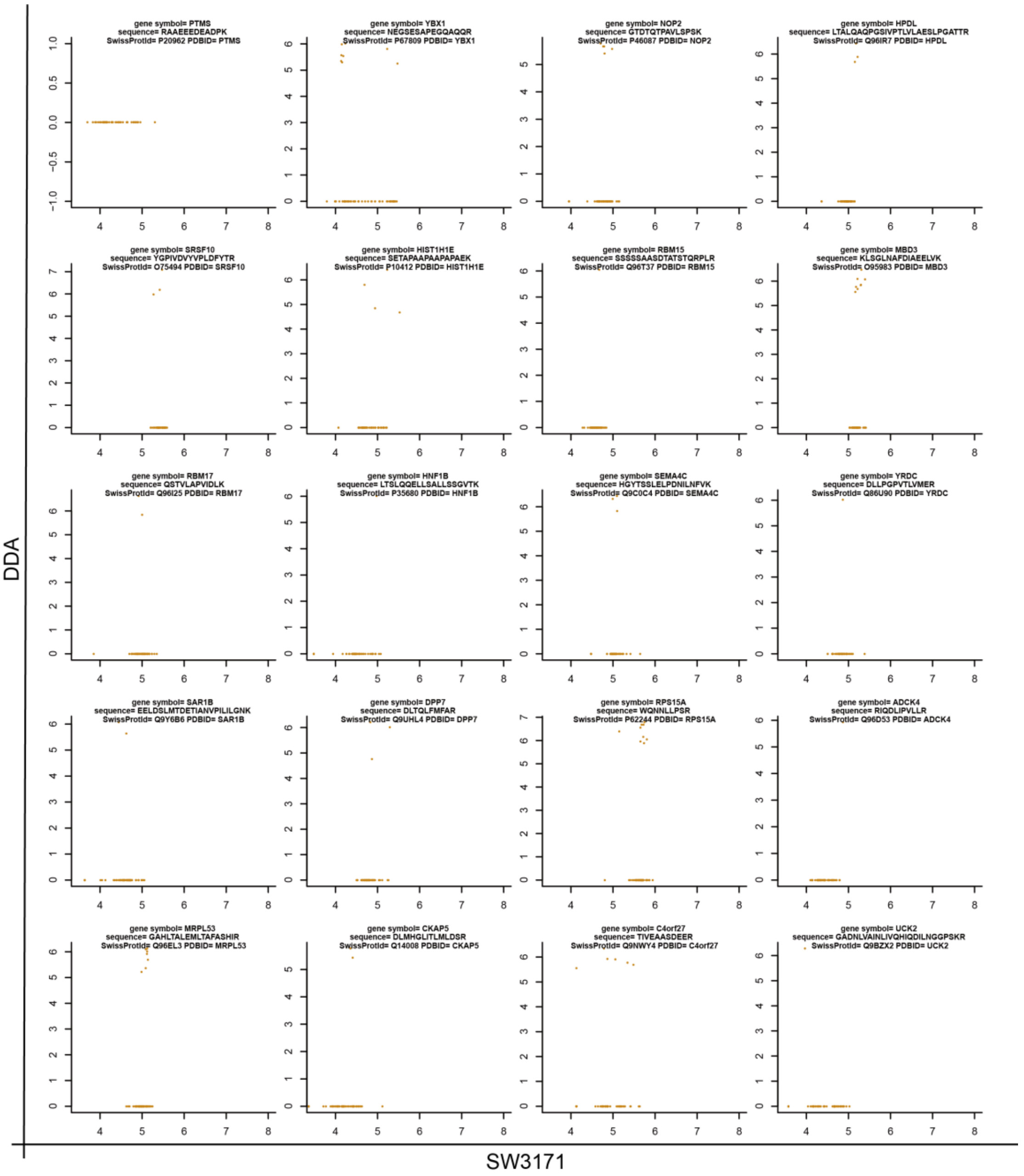
gene symbol= PTMS
sequence= RAAEEEDEADPK
SwissProtId= P20962 PDBID= PTMS
gene symbol= YBX1
sequence= NEGSESAPEGQAQQR
SwissProtId= P67809 PDBID= YBX1
gene symbol= NOP2
sequence= GTDTQTPAVLSPSK
SwissProtId= P46087 PDBID= NOP2
gene symbol= HPDL
sequence= LTALQAQPGSIVPTLVLAESLPGATTR
SwissProtId= Q96IR7 PDBID= HPDL
gene symbol= SRSF10
sequence= YGPIVDVYVPLDFYTR
SwissProtId= O75494 PDBID= SRSF10
gene symbol= HIST1H1E
sequence= SETAPAAPAAPAPAEK
SwissProtId= P10412 PDBID= HIST1H1E
gene symbol= RBM15
sequence= SSSSSAASDTATSTQRPLR
SwissProtId= Q96T37 PDBID= RBM15
gene symbol= MBD3
sequence= KLSGLNAFDIAEELVK
SwissProtId= O95983 PDBID= MBD3
gene symbol= RBM17
sequence= QSTVLAPVIDLK
SwissProtId= Q96I25 PDBID= RBM17
gene symbol= HNF1B
sequence= LTSLQQELLSALLSSGVTK
SwissProtId= P35680 PDBID= HNF1B
gene symbol= SEMA4C
sequence= HGYTSSLELPDNILNFVK
SwissProtId= Q9C0C4 PDBID= SEMA4C
gene symbol= YRDC
sequence= DLLPGPVTLVMER
SwissProtId= Q86U90 PDBID= YRDC
gene symbol= SAR1B
sequence= EELDSLMTDETIANVPILILGNK
SwissProtId= Q9Y6B6 PDBID= SAR1B
gene symbol= DPP7
sequence= DLTQLFMFAR
SwissProtId= Q9UHL4 PDBID= DPP7
gene symbol= RPS15A
sequence= WQNNLLPSR
SwissProtId= P62244 PDBID= RPS15A
gene symbol= ADCK4
sequence= RIQDLIPVLLR
SwissProtId= Q96D53 PDBID= ADCK4
gene symbol= MRPL53
sequence= GAHLTALEMLTAFASHIR
SwissProtId= Q96EL3 PDBID= MRPL53
gene symbol= CKAP5
sequence= DLMHGLITLMLDSR
SwissProtId= Q14008 PDBID= CKAP5
gene symbol= C4orf27
sequence= TIVEAASDEER
SwissProtId= Q9NWY4 PDBID= C4orf27
gene symbol= UCK2
sequence= GADNLVAINLIVQHIQDILNGGPSKR
SwissProtId= Q9BZX2 PDBID= UCK2
DDA
SW3171

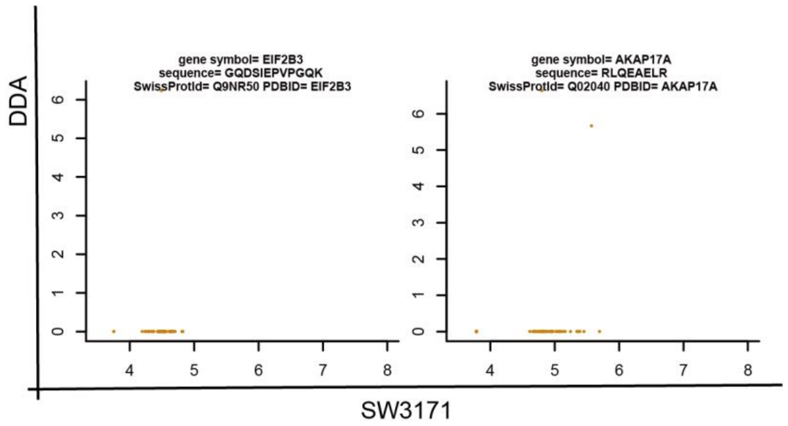


**Supplementary Figure 7. Bar plots and scatter plots for 142 representative peptides, which are all quantified cross all NCI-60 cell lines by SWATH but not quantified by DDA.** This figure shows the data completeness difference of the two data sets.

**Supplementary Figure 8**

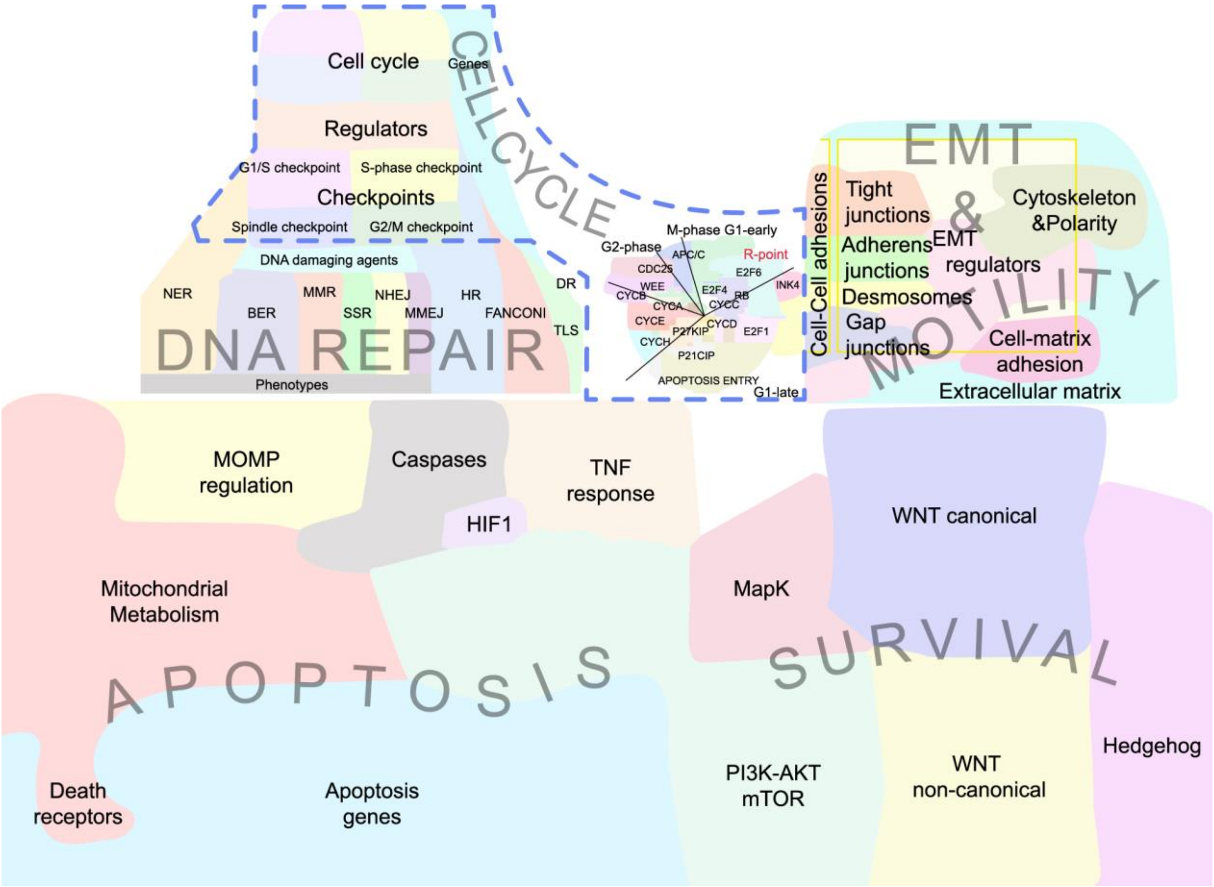


**Supplementary Figure 8. Global cancer signaling pathway maps based on Atlas of Cancer Signaling Network (ACSN) pathways** [3] (www. acsn.curie.fr). The annotations of the pathway map are shown.

### Supplementary Figure 9

**A**

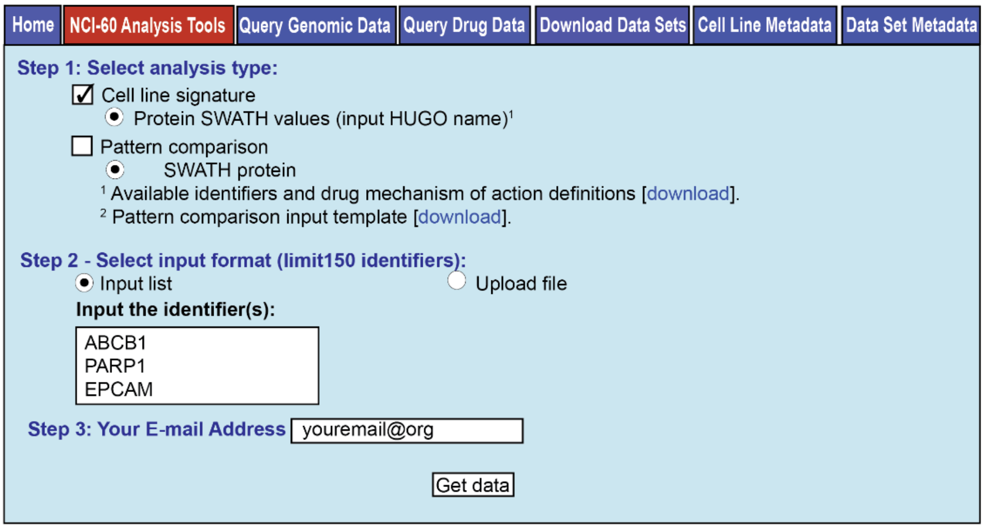


**B**

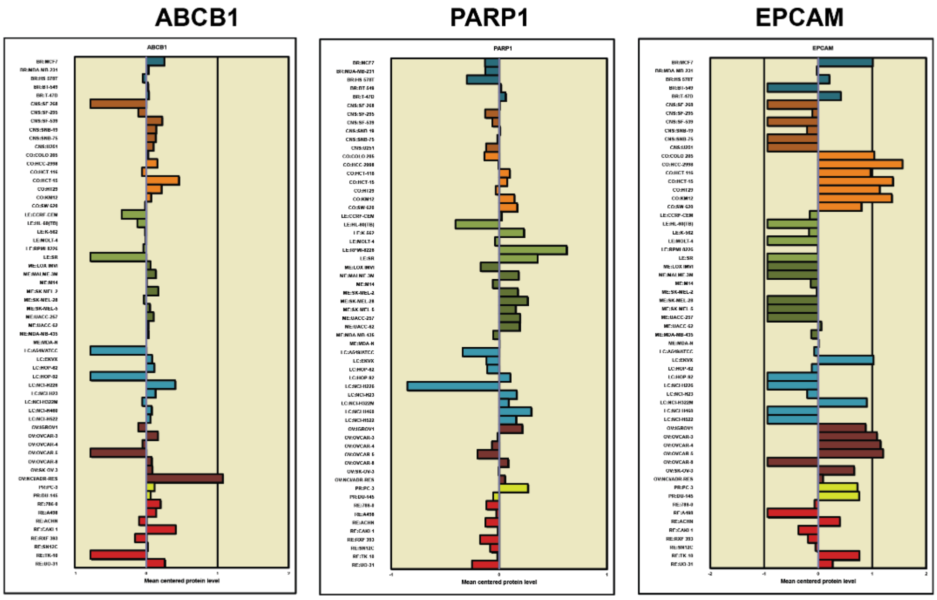


**Supplementary Figure 9. Access to NCI-60 proteotype in Cellminer.** To facilitate data access, visualization, and comparison with other forms of genomic and pharmacological data for the NCI-60 cancer cell lines, we have incorporated the SWATH data within CellMiner [4 5]. The CellMiner web site allows the data to be retrieved or used in several ways [6]. (**A**) The "Download Data Sets" tab allows either the total 3,171 proteins, or the 22,554 peptides data sets to be downloaded. This data will primarily be of use in computational biology pipelines. The "Query Genomic Data" tab allows up to 150 proteins or peptides to be accessed (using the "Gene" or "Peptide" pull downs), queryable by gene name or peptide peak identifier, chromosomal or genomic location. Data is sent in both Excel (.xls) and text (.txt) format. The

“NCI-60 Analysis Tools” tab (**A**) provides “Cell line signatures”. To obtain “Cell line signatures” for genes, select “Cell line signature” in Step 1, and then “Protein SWATH values”. In Step 2, up to 150 genes of interest may be input by either typing in the gene names in the “Input the identifier” box, or uploading them as a text or Excel file using the “Upload file” radio button. In Step 3, enter your e-mail address, and click “Get data”. Results will be sent by e-mail for each gene, with a link to download the results. This file contains three worksheets: i) tabular mean centered protein levels ratios as a both a bar plot and tabular data, and the peptide peak information for that gene ii) “Bin protein levels” with a histogram of the protein levels and iii) and “Footnotes”. (**B**) provides examples of three genes of interest. These “Cell line signatures” can also be used as input for the Pattern Comparison tool (also within the NCI-60 Analysis tools section) which provides correlated molecular and compound activity data. All available gene and peptide identifiers are available as a list within the “Available identifiers and drug mechanism of action definitions” as a download within the “NCI-60 Analysis Tools” tab.
